# CGGBP1 from higher amniotes restricts cytosine methylation and drives a GC-bias in transcription factor binding sites at repressed promoters

**DOI:** 10.1101/2024.11.14.623544

**Authors:** Praveen Kumar, Ishani Morbia, Aditi Lakshmi Satish, Subhamoy Datta, Umashankar Singh

**Affiliations:** Department of Biological Sciences and Engineering, Indian Institute of Technology Gandhinagar, Gandhinagar, India; Applied Tumor Genomics, Faculty of Medicine, University of Helsinki, Helsinki, Finland

**Keywords:** Transcription Factor-Binding Sites (TFBSs), promoter, cytosine methylation restriction, GC content, C-T transition, GC retention, amniote, conserved, evolution

## Abstract

CGGBP1, a 20 kDa protein, has several functions associated with its DNA-binding through a C2H2 zinc finger. A range of studies have shown that GC richness, inter-strand G/C-skew and low cytosine methylation are associated with CGGBP1 occupancy. The non-preference of any sequence motif as CGGBP1 binding site suggests widespread association of CGGBP1 with DNA including at potent transcription factor binding sites (TFBSs) in promoter regions. The evolutionary advantage of such a design remains unclear. The regulatory interference by human CGGBP1 at TFBSs is supported by purifying selection in the DNA-binding domain of CGGBP1 and its requirement for gene repression as well as restriction of cytosine methylation at GC-rich TFBSs. Here we describe an evolutionary trajectory of this property of CGGBP1 by combining global gene expression and cytosine methylation analyses on human cells expressing CGGBPs from four different vertebrates (representatives of coelacanth, reptiles, aves and mammals). We discover a potent cytosine methylation restriction by human CGGBP1 at some GC-rich TFBSs in repressed promoters. Further, we combine a high-throughput analysis of GC compositional bias of these CGGBP-regulated TFBSs from available orthologous sequences from a pool of over 100 species. We show that cytosine methylation restriction by CGGBP1 is tightly linked to GC retention in a set of TFBSs. Orthology analyses demonstrate that this property of CGGBPs has evolved in higher amniotes (aves and mammals) with lineage-specific heterogeneities in lower amniotes (reptiles). CGGBP1 ChIP-seq data suggest that occupancy of CGGBP1 at these target TFBSs plays a crucial role in their low methylation, GC-biased evolution and associated functions in gene repression.

**Highlights:**

- Resemblances in gene repression by overexpression of CGGBP1 from higher amniotes (*Homo sapiens* and *Gallus gallus*) is enhanced upon heat stress and differs from the non-repressive effects of lower amniotic CGGBPs (*Anolis carolinensis* and *Latimeria chalumnae*).
- Gene repression by higher amniotic CGGBP1 is associated with restriction of cytosine methylation at specific GC-rich TFBSs in 1 kb promoters of target genes. Lower amniotic CGGBPs allow TFBS cytosine methylation and C-T transitions.
- Orthologs of CGGBP1-repressed genes from >100 vertebrates show signs of accelerated C-T losses explicitly in the TFBSs at which higher amniotic CGGBP1 restricts cytosine methylation. Such a TFBS GC-loss difference between lower and higher amniotes is restricted to genes repressed by higher amniotic CGGBP1 at physiological temperature, not heat stress.
- This higher amniote-specific cytosine methylation restriction by CGGBP1 has likely influenced the differences between GC-rich TFBS composition and their abundance in target gene promoters throughout vertebrate evolution.

**Summary:** Evolution of transcription factor binding sites (TFBSs) depends on a variety of factors including cytosine methylation-associated C-T transition rates. Most of our understanding of TFBS evolution is based on omic-scale sequence comparisons with only circumstantial evidence for the relationship between the TFBSs and physiological adaptation. We report a TFBS landscaping function for CGGBP1 by expressing it’s different taxon-derived forms in human cells through profiling of global gene expression and cytosine methylation alongside a meta-analysis of C-T transition rates from over 100 vertebrae genomes. We show that CGGBP1 from higher amniotes restricts cytosine methylation and maintains GC-rich TFBSs in target gene promoters for repression. This epigenetic affection of TFBS evolution by CGGBP1 is selectively seen at genes repressed at physiological temperature only and not under heat stress when gene repression by CGGBP1 becomes largely transcription factor binding site independent. Our findings connect epigenetic mechanisms to cellular physiology through TFBS evolution linked with changes in CGGBP1.

## Introduction

CGGBP1 is a gene located on the short arm of human chromosome 3 coding for a 20 kDa protein. CGGBP1 protein contains a DNA-binding domain with clearly identifiable cysteine and histidine residues located in a manner consistent with the presence of a C2H2 Zinc finger domain (1). Multiple studies demonstrate that CGGBP1 is a DNA-binding protein with a preference for GC-rich double-stranded DNA (2–5). Although the C2H2 DNA-binding domain is clearly identifiable at the N-terminus of CGGBP1, the rest of CGGBP1 protein remains poorly understood. Except for a NLS at 55-80, there is no identifiable domain which can be used to deduce candidate functional regions for mutation and functional analyses (UniProt and NCBI Gene). There is absence of any known domains in the C-terminal half of CGGBP1 even as Alphafold shows that this part of CGGBP1 has helices interrupted by loops. A poorly conserved DBD from the Hermes transposons has been purported to bear some similarities to this C- terminal part of CGGBP1 (1,6).

With its relatively simple structure and small size, it is interesting that CGGBP1 has become associated with many different functions. It is involved in gene repression through CGG repeats (1–3), general gene repression through regulation of RNA Pol2 recruitment and Alu RNA transcription (4,5), mitigation of cytosine methylation (6–8), suppression of endogenous DNA damage response (9,10), repression of G-quadruplex formation (11) and regulation of chromatin barrier elements marked by CTCF occupancy (12). Most of the functions attributed to CGGBP1 involve DNA-binding (1,3,4,7–9,11–14) indicating that the DNA-binding domain and associated DNA-protein interactions between CGGBP1 and its target DNA are required for the functions of CGGBP1. The importance of CGGBP1 is demonstrated by the multitude of well-conserved processes it regulates as well as its unique evolutionary history (additional file 1) (15).

The Zn finger DNA binding domain of CGGBP1 shows similarity to the BED domain of the hAT transposase suggesting a transposon-associated origin of the protein (5,13). CGGBP1 is almost entirely conserved in mammals, strongly conserved in homeotherms and all amniotic forms of CGGBP1 seem to derive from a common ancestor of mammals and the extant archosauromorphs (additional file 1, fig1) (15). The latter seems to be derived from a common tetrapod ancestor, a coelacanth, although CGGBPs have been retained only in amniotes (15) and most amphibians have lost it (additional file 1, fig 2 shows the known amphibian forms from Newt and *Rhinatrema*) (13,16). The extant coelacanth *Latimeria* has multiple forms of CGGBPs of varying structures (13). By comparing the various forms of CGGBPs no singular functional or structural conservation can be deduced. The various CGGBPs, including the multiple forms present in parallel in coelacanthini, show different DNA-binding preferences with the avian and human CGGBP1 showing some similarity in vitro (13). Interestingly the N-terminal part of CGGBP1, which contains the DNA-binding domain is more conserved than the C-terminal part in which no known functional domains have been reported (additional file 1, fig 1-2) (15). Recent works suggest that the direct binding to CGG triplets is tightly linked with the DBD conserved between aves and mammals (15). However, the genome-wide occupancy of CGGBP1 defies this expectation and shows binding to sequences in which CGG repeats are not enriched (4,5). Some more recent CGGBP1 ChIP-seq (17) however have not been evaluated from this viewpoint. Thus the mechanism of DNA-binding of CGGBP1 and its dependence on the DBD remains unclear. The reason why a GC-rich DNA-binding protein would rapidly evolve in tetrapods, become conserved in amniotes with purifying selection in homeotherms and still maintain a very heterogeneous DNA-sequence preference is perplexing. Clearly, motif-specific DNA binding through the conserved C2H2 DBD is not a feature of CGGBP1 that can be deduced from these observations. Direct or indirect interactions with DNA through the less well conserved (15) C-terminal part of CGGBP1 most likely affects its overall function. The recent studies with C- or N-truncated parts of CGGBP1 shed light on the functional variability conferred by the C-terminal part of CGGBP1 (15). It seems that the C- and N-terminal parts of CGGBP1 cooperate to generate the grand functional outcomes in which the C-terminal part seems to be needed for the diversity of effects the N-terminal part can have through its default interactions with the DNA.

**Figure 1.**
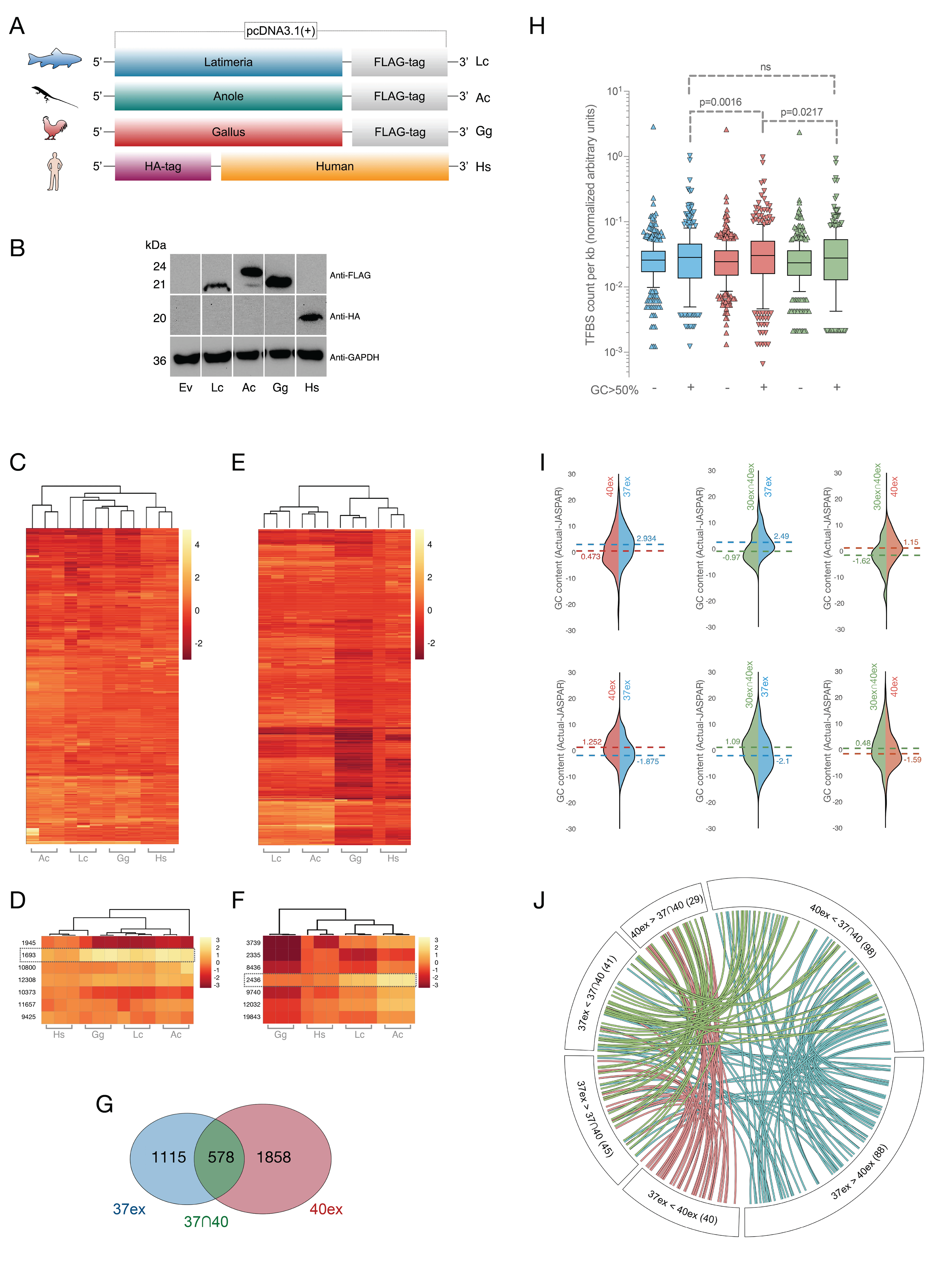
GC contents of TFBSs in target gene promoters as a major feature of gene repression by representative form of mammalian and avian CGGBP1s. A: Schematic (not to the scale) of the four different evolutionary forms of CGGBPs used in this study cloned into an constitutively active eukaryotic expression vector and represented as a two letter code: Lc (*Latimeria*, non-amniote), Ac (*Anole;* reptile), Gg (*Gallus*; ave) and Hs (Human; mammal) The C-terminal FLAG or an N-terminal HA tags are indicated. B: The Lc, Ac, Gg and Hs forms expressed in HEK293T cells at comparable levels and expected molecular weights with no indication of a lack of protein stability. An empty vector control is indicated as Ev. C: Heat map of global gene expression profiles (M values against Ev) using Agilent microarrays at 37°C show reproducible gene expression patterns in three replicate experiments. D: Classification of gene expression changes at 37°C against Ev using k-means identifies a cluster of 1693 genes which are repressed only by Hs (boxed row). E: Heat map of global gene expression profiles (M values against Ev) using Agilent microarrays at 40°C show that the large scale transcription repression is reproducibly captured in three independent replicates. F: A k-means clustering of genes, similar to that employed for the 37°C genes (D) captured 2436 genes repressed by Hs and Gg at 40°C (boxed row). G: The two gene sets highlighted in C and D have an unexpectedly high commonality with 1115 genes as 37°C-exclusive (37ex; blue), 1858 genes as 40°C-exclusive (40ex; red) and 578 genes commonly repressed at both the conditions (37∩40; green). The three gene sets were filtered to retain only unambiguous promoters: 37ex with 811 genes, 40ex with 1511 genes, and 37∩40 with 470 genes. H: The 40ex gene promoters (−1 kb from TSSs) have higher abundance of GC-rich TFBSs (n=310, a paired t-test between 40 ex and 37ex confirms this observation with a p-value of 0.0016. Additionally, the comparison of 40ex with 37∩40 showed a p-value of 0.0217). The 37ex and 37∩40 promoters do not show any quantitative enrichment of GC-rich TFBSs. I: The 37ex promoters show a qualitative difference in TFBSs in the form of unexpectedly high GC contents for most TFBSs. J: The circos plot shows the exact counts and overlaps of TFBS with higher or lower than expected GC content identities in various comparison sets. The > and < symbols depict the direction of GC content difference and the number of TFBSs showing a pattern of difference is indicated in parentheses. The arms connecting the six different comparison sets follow TFBSs common to different comparison sets.

**Figure 2.**
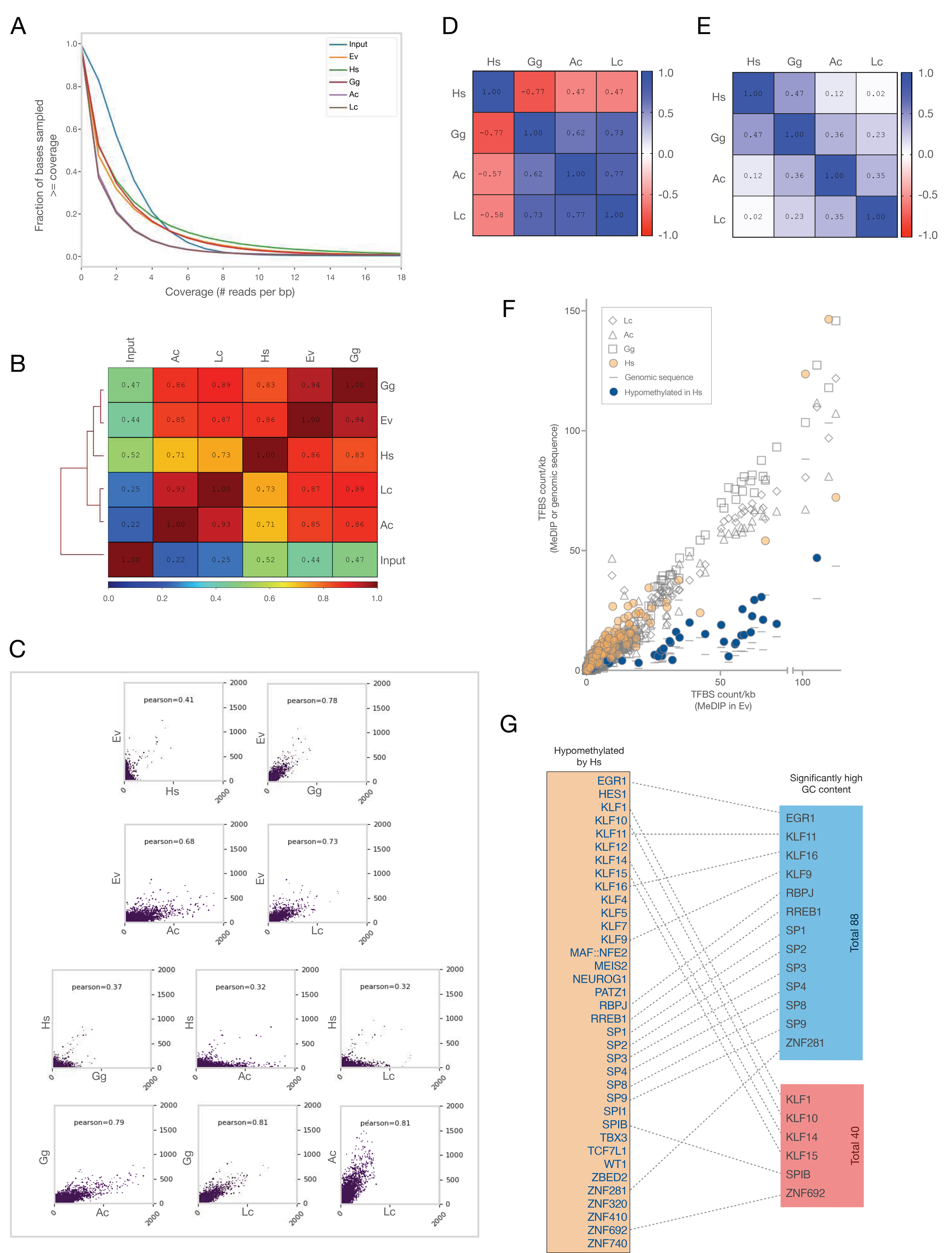
Differences in cytosine methylation patterns induced by Hs, Gg, Ac and Lc recapitulate interspecies differences in gene expression patterns. A: More genomic regions have lower methylation signals in Hs, Gg and Ev as compared to Ac and Lc. MeDIP coverage analysis for the five samples and the input shows two different enrichment patterns, one in Lc and Ac and another in Gg, Hs and Ev. The enrichment in Hs, Gg and Ev is majorly in the moderate signal range of 5 to <25. The enrichment in Ac and Lc is spread out in a large range of higher methylation signals starting from 25 and more (see genome-wide frequency distributions in Fig. S6A). The differences between the two patterns clearly shows that the areas under the curves for Hs, Gg, Ev are higher than those for Ac and Lc in the regions of no enrichment (signals up to 5) as well as in regions of enrichment in all samples (signals more than 8). B: Genome-wide correlation of normalized MeDIP signals and clustering demonstrate that the methylation pattern in Ev is the least affected by Hs and Gg. Expression of Ac and Lc change the methylation pattern distinctly. C: Scatter plots of methylation signals in common regions across different sample pairs show that Hs expression restricts methylation levels as compared to other forms of CGGBPs, including the endogenous CGGBP1. D and E: Correlations between methylation patterns at TFBSs in 1 kb promoters of genes repressed by CGGBP1 at 37°C (D) or 40°C (E) show differences between the two gene sets. The methylation restrictive effect of Hs CGGBP1 at TFBSs is stronger at the promoters of 37°C genes (D) than 40°C genes (E). F: TFBS occurrences in MeDIP reads mapping to the repressed gene promoters from Lc (empty diamonds), Ac (empty triangles), Gg (empty squares) and Hs (beige circles) when compared against those in Ev show a specific methylation restriction in Hs for 36 factors (>= 2 folds decrease compared to Ev; blue circles). The magnitudes of methylation levels inferred from TFBS occurrence per kb promoter sequence in each sample is benchmarked against the minimum expected occurrence of these TFBSs in the genomic sequence of these gene promoters (horizontal strokes). G: List of 36 hypomethylated TFBS by Hs show an unexpectedly high overlap with the TFBSs having significantly higher GC content in Hs repressed promoters (13 in 37ex promoters (blue box) and 6 in 40ex promoters (red box)). The actual JASPAR identities of the TFs are connected with dashed lines.

The C-terminal part of CGGBP1 has no known motifs or domains to target for functional studies. Also, the evolutionary influence of CGGBP1 lies in a combination of the functions of C- and N- termini. The outcome of interactions of different taxa-specific CGGBPs with different taxon group-specific DNA sequence motifs suggests that CGGBPs have evolved to target different motifs. However, the influence of CGGBPs from different taxa on the motifs they bind to remains unknown. A simple survey of the CGGBP1-regulated motifs reported earlier (15) in the light of the data presented here shows that many of these motifs are not as GC-rich in non-amniotes as compared to their orthologs in amniotes, especially homeotherms. This gives rise to a possibility that different sequence motifs are regulated by different taxon-specific forms of CGGBP1 through a combination of its N-term for binding and C-term for its effect. These recent findings (15) indicate a mechanism in which the CGGBP1 regulates the target site GC-richness and needs to be explored.

We report that different taxon-specific forms of CGGBPs bind to and regulate cytosine methylation of target sites differently. We employ CGGBPs from representative taxa expressed in human cells to study the effects of different evolutionary forms of CGGBPs on gene expression and cytosine methylation. Our experimental results, combined with genome-wide analyses of CGGBP1-regulated transcription factor binding sites, shed new light on genome, epigenome and transcriptome regulation through CGGBP1. We show that CGGBP1 has evolved in homeotherms, especially in mammals, to prevent cytosine methylation at GC-rich TFBSs associated with proximal promoters of its target genes. These results suggest that regulation of short GC-rich elements through prevention of cytosine methylation in gene promoters has been a crucial mechanism in amniote and homeotherm evolution and CGGBP1 is a key player in the process.

## Results

### Gene repression by CGGBP1 reveals a heat stress-dependent difference between non-amniotes, reptiles, aves and mammals

To explore the effects of various evolutionary forms of CGGBPs on gene expression, we expressed four distinct vertebrate forms of CGGBPs in HEK293T cells (Fig 1, A and B). The endogenous human CGGBP1 was knocked down first, followed by the introduction of the non-human CGGBPs forms (Fig S1A). Nuclear-cytoplasmic fractionation assays show that all vertebrate forms of CGGBPs are primarily localized to the nucleus following their synthesis in the cytoplasm, with minimal to no CGGBPs detected in the cytoplasmic fraction (Fig S1B).

Global gene expression analysis using Agilent arrays showed that overexpression of human or non-human forms of CGGBPs cause different patterns of gene expression disturbances (Fig 1C). As has been described earlier we could observe a general repression of gene expression by human CGGBP1. In comparison, the other forms of CGGBPs had two different effects; a large-scale repression of gene expression and a restricted induction of gene expression (Fig S2). Induction of gene expression was limited to non-amniotic and reptilian CGGBPs expression with different gene sets induced or repressed by different forms of CGGBP1.

The Hs CGGBP1 is predominantly repressive in nature. In order to identify species-specific differences in the mechanism of gene repression by different forms of CGGBPs we did not focus on different sets of genes induced or repressed by amniotic or non-amniotic CGGBPs. Instead we focused on a small set of genes which showed a clear loss of gene repression when human CGGBP1 was replaced with alternative forms. Using k-means clustering we could consistently identify a set of 1500-2000 genes with a human-specific repression. From these one such representative cluster of 1693 genes was further analyzed (Fig 1D). These genes did not show any functional group enrichment. To identify the mechanisms of gene repression specific to the human form, one possibility we focused on is the presence of proximal *cis* regulatory elements or DNA sequence features associated with these genes through which these genes were repressed by CGGBP1 in a proximal sequence-dependent manner. A JASPAR-wide motif search showed no obvious deviations from genome-wide motif enrichment in the 1 kb promoter regions of these genes (not shown).

Previous reports have shown that CGGBP1 participates in a sequence-directed repression which is sensitive to heat stress (18). To narrow down our search for any CGGBP1 target TFBSs in the repressed gene promoters we subjected the CGGBP overexpression system to heat stress (Fig S3). Previous studies have shown that upon heat stress there is a switch of CGGBP1 from Alu-SINEs to tRNA genes. The induction of Alu transcription generates Alu RNA which binds to and inhibits RNA Pol2 activity (4,19). Such a gene repression by CGGBP1 in response to heat stress takes place in trans and is independent of any *cis* regulatory motifs. So we used heat stress as a mechanism to override gene repression through *cis* regulatory elements by CGGBP1. Studying gene expression patterns at 40°C would hence help us compare the stress-free and heat stress-associated promoter profiles and thus establish the *cis* regulatory feature dependence of gene repression by various forms of CGGBPs. In order to reliably utilize the expression data from the 40°C experiments, we first needed to establish that the various forms of CGGBPs retained their functions at 40°C. If the various forms of CGGBPs were incapacitated by heat stress then we would not observe differences in heat stress-induced gene expression between them. We compared the changes in gene expression brought about by heat stress under the expression of each of the different forms of CGGBPs (Fig S4). There were striking patterns of similarity between heat stress-induced gene repression only between Hs and Gg CGGBP1 suggesting that the Gg CGGBP1 did not lose its function in human cells even under heat stress (Fig S5). In samples expressing Ac and Lc CGGBPs, there was a strong loss of gene repression similar to what was observed at 37°C (Fig 1C). These findings showed that even at 40°C the various forms of CGGBPs retained their functional integrity as expected at 37°C.

The change in gene expression pattern observed at 40°C was different from that at 37°C in a manner such that the CGGBP closest to the Hs form, Gg, was also able to regulate the gene expression pattern similar to that of Hs (Fig 1E and Table S1-S4). By k-means clustering of the 40°C gene expression data, we observed that under heat stress a distinct cluster of genes emerged for which the expression deregulation was similar to that observed at 37°C (Fig 1F). These genes were repressed by Hs CGGBP1 but not by Ac or Lc CGGBP. The only difference was that Gg CGGBP1, the only alternative homeotherm-derived CGGBP1 in our experiments which also has the least difference from the Hs form, became repressive as well. The repression of these genes by Hs CGGBP1 was expected to be independent of proximal *cis* regulatory elements (4) as upon heat stress CGGBP1 represses gene expression in trans through Alu RNA induction.

In order to fetch a set of genes which would reveal DNA sequence features of proximal *cis* regulatory sequences important for repression by CGGBP1, we extracted the genes exclusive to the 37°C Hs and 40°C Hs repression sets. Interestingly, there were 470 genes for which the non-repressive Gg CGGBP1 showed a heat stress-dependent repressive effect (Fig 1G).

Through this exclusion of genes which might be repressed due to alternative mechanisms, we retained a set of 811 genes exclusively repressed by Hs at 37°C to study further (Fig 1G). The motif features of the promoters of these 811 genes were compared against those of the 1512 genes exclusively repressed by Hs and Gg CGGBP1 at 40°C and the 470 genes repressed by Hs CGGBP1 regardless of the heat stress (Fig 1G) (Genes containing unambiguous promoter were retained and coordinates are mentioned in supplementary data Tables S5-S7).

Search for the entire JASPAR (vertebrate) motif set showed that there were no overall differences in motif occurrences between the three gene sets (not shown). However, the motifs with GC content of more than 50% were found to occur significantly more in 40ex promoters as compared to 37ex or 37∩40 (Fig 1H). These differences in GC-rich motif counts were restricted to the TFBSs as the promoter sets of these gene sets were not significantly different in overall GC contents (Table S8). Intriguingly, the high occurrence of GC-rich motifs in 40ex promoters did not affect the overall promoter GC content. We interpreted these findings as indicative of an inverse relationship between GC-rich motif occurrences and their actual GC contents in these promoters. These findings suggested that under the physiological conditions of 37°C (37ex) CGGBP1 represses genes through a combination of motifs occurrence and their GC contents regardless of the overall GC content of the promoter sequences.

To further understand the mechanism of differential gene repression by Hs and alternative forms of CGGBPs through TFBSs in target promoters, we analyzed motif GC contents in detail. The actual GC contents of the motif hits as well as their counts were compared for 716 JASPAR motifs in the three gene sets. Actual GC contents for several TFBSs were different in the different gene sets at the same threshold of motif detection (1e-5) (Tables S9-S14). For a much larger set of genes the actual motif GC content was significantly higher in 37°C (37ex) genes and 470 (37∩40) genes than that in the 40°C (40ex) genes (Fig 1, I and J, Tables S9 and S11).

The differences in GC contents for TFBSs present in 37ex and 40ex gene sets showed that Hs CGGBP1 has a larger spectrum of target motif GC content in its target repressed promoters than that of the alternative forms of CGGBPs. Of the alternative forms, Gg showed a heat stress-inducible facultative regulatory capacity for the high GC content TFBS regulation. The inability of Ac and Lc CGGBPs to repress the 37ex or 40ex genes suggested that the GC content-dependent regulation of TFBSs is an evolutionarily acquired feature of CGGBP1 manifested facultatively in aves and constitutively in mammals, considering that the Lc, Ac, Gg and Hs forms of CGGBPs represent coelacanthini, reptilia, aves and mammals respectively.

CGGBP1 binding to target GC-rich sequences exerts the dual effects of transcription repression and restriction of cytosine methylation. The mechanism of gene repression by the various CGGBP forms must be such that it would explain differential gene repression as well as differences in the TFBS GC contents. We investigated the possibility that cytosine methylation at TFBSs in target gene promoters is mitigated by CGGBP1 and through the differences in cytosine methylation mitigation properties the different forms of CGGBPs exert differential gene repression. The differences in TFBS cytosine methylation would also explain the passive mechanisms of GC content differences. The genomes of cells expressing the representative forms of CGGBPs were assayed for global methylation patterns by MeDIP-seq.

### Species origin of CGGBPs determines cytosine methylation restriction at some GC-rich TFBSs

The GC-content differences of TFBSs in promoters of target genes existed alongside no significant differences in GC contents of the promoters. Such a TFBS-specific GC content maintenance could be explained by mechanisms which affect mutation rates of cytosines. Cytosine methylation accelerates C-T transition. CGGBP1 restricts cytosine methylation and at the same time represses transcription (4,6,18). A TFBS-specific restriction of cytosine methylation by CGGBP1 could lead to the GC content differences observed. We performed methylcytosine DNA immunoprecipitation-sequencing from Hs, Gg, Ac and Lc expressing cells using a protocol described elsewhere. MeDIP summary is shown in Table S15.

A genome-wide comparison of cytosine methylation signals was performed using all the MeDIP- seq reads independently captured in various experiments. Different patterns of cytosine methylation enrichments were obtained in Hs, Gg, Ac and Lc. Recapitulating our observations with the gene expression patterns, of all the MeDIP samples the methylation enrichment between Hs and Gg were similar over a large dynamic range of MeDIP signals (Fig 2A). These comparisons showed that genome-wide methylation patterns were more similar between Hs and Gg which clustered together. In comparison, Ac and Lc showed a distinct pattern and clustered separately (Fig 2B; Fig S6, A and B). Noteworthily, the cytosine methylation patterns in Hs and Gg were most similar to Ev whereas Ac and Lc showed maximum difference from Ev and clustered together (Fig 2B; Fig S6, A and B). These genome-wide comparisons showed that there was a general mitigation of methylation in Hs (Fig 2C). For a definitive comparison we focussed on regions where methylation was captured in all the samples. It was confirmed that Hs expression exhibited a strong methylation mitigation. Whether the cytosine methylation mitigation takes place at regulatory elements in the genome, we analyzed MeDIP-seq signals at promoters, enhancers and conserved regulatory elements (UCSC). We found that Hs CGGBP1 maintains a level of cytosine methylation at these regulatory elements while other CGGBPs fail to (Fig S7, A, B and D). Methylation signals on array-wide gene promoters also tell the similar story, the closest form to the Hs CGGBP1 that is Gg appears to achieve the same but not as efficient in the alien environment (Fig S7C). While Ac and Lc CGGBPs completely fail to maintain cytosine methylation in the human cells.

The presence of JASPAR motifs in MeDIP-seq was used as a measure of TFBS methylation. 750 JASPAR Vertebrate motifs were detected in the randomly sampled MeDIP reads and were analyzed if their methylation was different between Hs, Gg, Ac and Lc. For the majority of TFBSs overexpression of different forms of CGGBP1 did not affect cytosine methylation (Fig S8). MeDIP signals were found 1 kb upstream of the 37ex, 40ex, and 37∩40 gene promoters, with the signal in the 37ex region mirroring gene expression changes at 37°C, while no such pattern was seen in the 40ex and 37∩40 promoters. (Fig S9). This indicates that methylation-dependent regulation plays a specific role in the expression of the 37ex genes. We then examined MeDIP reads from Hs, Gg, Ac, and Lc that mapped to the promoters of the 37ex, 40ex, and 37∩40 genes to identify transcription factor binding sites (TFBSs). Compared to Ev, only Hs CGGBP1 caused a significant alteration in TFBS methylation specifically at the 37ex gene promoters (Fig 2D). Interestingly, under heat stress conditions, Gg closely related to the homeothermic form of CGGBP1 exhibited a similar TFBS methylation pattern to that observed with Hs CGGBP1 at 40ex and 37∩40 gene promoters (Fig 2E and Fig S10). Hs turned out to be the only CGGBP1 which mitigated cytosine methylation of a set of 36 TFBSs only in the promoters of 37ex gene promoters (Fig 2F and table S16). Expression of the other forms of CGGBPs did not alter cytosine methylation of TFBSs in these sets of CGGBP1 target genes (Fig 2F).

We established that the reduced occurrence of these motifs were not due to skewed TFBS contents in the promoter sequences (Fig S11). The promoter and TFBS methylation findings mentioned above were further tested and verified by ruling out two major counter-possibilities: the lower abundance of these TFBSs in MeDIP signals at 37ex promoters could be simply due to the sequence features of the 37ex gene promoters which are deficient in these TFBSs and that there was a systemic bias due to which our MeDIP failed to capture these TFBSs as methylated. The genomic sequences and input data for these 37ex promoter genes were as rich in these TFBSs as expected thus proving that the deficiency of these TFBSs in MeDIP data could not be due to biased sequence properties of the regions of interest (Fig 2F). Additionally, the same TFBSs with low methylation capture in 37ex promoters were abundant in 40ex as well as in 37∩40 gene promoters in the same MeDIP datasets thereby nullifying any possibilities of a systemic bias against capture of some TFBSs.

The effect of Hs CGGBP1 on methylation restriction was clearly observed for TFBSs with a methylation signal above a threshold (Fig 2F). By filtering out the TFBSs with prominent methylation restriction we narrowed down to 36 TFBSs (Fig 2F; blue data points). 21 out of these 36 hypomethylated TFBSs overlapped with a set of 45 motifs independently reported as methylation restricted by full-length Hs CGGBP1 (15). These 36 motifs were also GC-rich, similar to the motifs discovered in the 37ex gene promoters (shown above in Fig 1, I and J). An unexpected 11 of these 36 motifs overlapped with the high GC-content motifs in 37ex gene promoters. These results showed that cytosine methylation mitigation of GC-rich TFBSs in a promoter context-dependent manner is tightly associated with gene repression by Hs CGGBP1 and that it differently marks genes for repression upon heat stress. Cytosine methylation restriction by CGGBP1 is a key feature of gene repression by Hs or a related avian CGGBP1 but not the reptilian or other distant forms of CGGBPs.

Put together these findings led us to the following conclusions: A set of GC rich TFBSs are methylation restricted by Hs CGGBP1. At conditions of no heat stress Hs CGGBP1 maintains these TFBSs at low methylation levels as well as in an expression repressed state. The GC rich TFBSs occur with varying actual GC content in different promoters. Their occurrence with higher GC content is linked to low cytosine methylation and repression by Hs CGGBP1. Based on these results we proposed a possibility that Hs CGGBP1 acts as a methylation restrictor at the promoters of a specific set of target genes and the high GC content of the TFBSs is a consequence of low C-T transition rates specific to some GC-rich TFBSs. We put this possibility to a rigorous test by analyzing the promoters of orthologs of CGGBP1 target genes in multiple species across different taxa.

### Large-scale orthology comparisons show that CGGBP1-repressed genes have a distinct evolutionary trajectory of TFBS GC-content

The inability of non-Hs CGGBPs in restricting TFBS methylation could be due to two different reasons: (i) the non-Hs CGGBPs are equally restrictive towards TFBS methylation as Hs CGGBP1 but fail to exert their function in the alien environment of a human cell, or (ii) TFBS methylation restriction is a function that has gradually evolved in mammals and other vertebrate CGGBPs do not have this function. If the latter possibility is true, then the orthologous promoter sequences of the 37ex, 40ex and 37∩40 genes shall show TFBS occurrence and GC-content in accordance with our findings that Hs has the strongest methylation restriction effect. Thus the actual effect of non-human CGGBPs on TFBS methylation restriction could be best studied by analyzing the TFBSs in orthologous promoter sequences from different taxa. This would also help us define if the single species representative forms of CGGBPs used in our experiments represent their respective taxon groups in general or not.

The transcription factors corresponding to 36 hypomethylated motifs were found to have orthologs or paralogs in most vertebrate taxa (additional file 2). We first compared the occurrence of the 36 hypomethylated TFBSs in 1 kb promoter regions genomewide from 105 species (see methods for details; species information is given in Table S17). As the GC contents vary between different species, we filtered out sets of promoters which fall in the maximal representation zone of 30-70% (Fig S12A). This range allows us to study TFBS occurrences in a GC content range that is comparable in the promoters of all species. We found that unlike non-amniotes, reptiles or aves, the majority of the 57 mammals analyzed clustered together with high occurrences of most of these TFBSs (Fig S12B). A similar enrichment of TFBSs in mammals was observed in a heterogeneous set of orthologs (orthologs were not always gene-matched in each species) for the 37ex gene promoters available from 75 species (heat map cluster 37ex ortholog genes; see method for details) (Fig 3A). These findings aligned with our proposition that the high occurrence of hypomethylated TFBSs correlates with methylation restriction by human CGGBP1, a representative of the highly conserved mammalian CGGBP1s.

**Figure 3.**
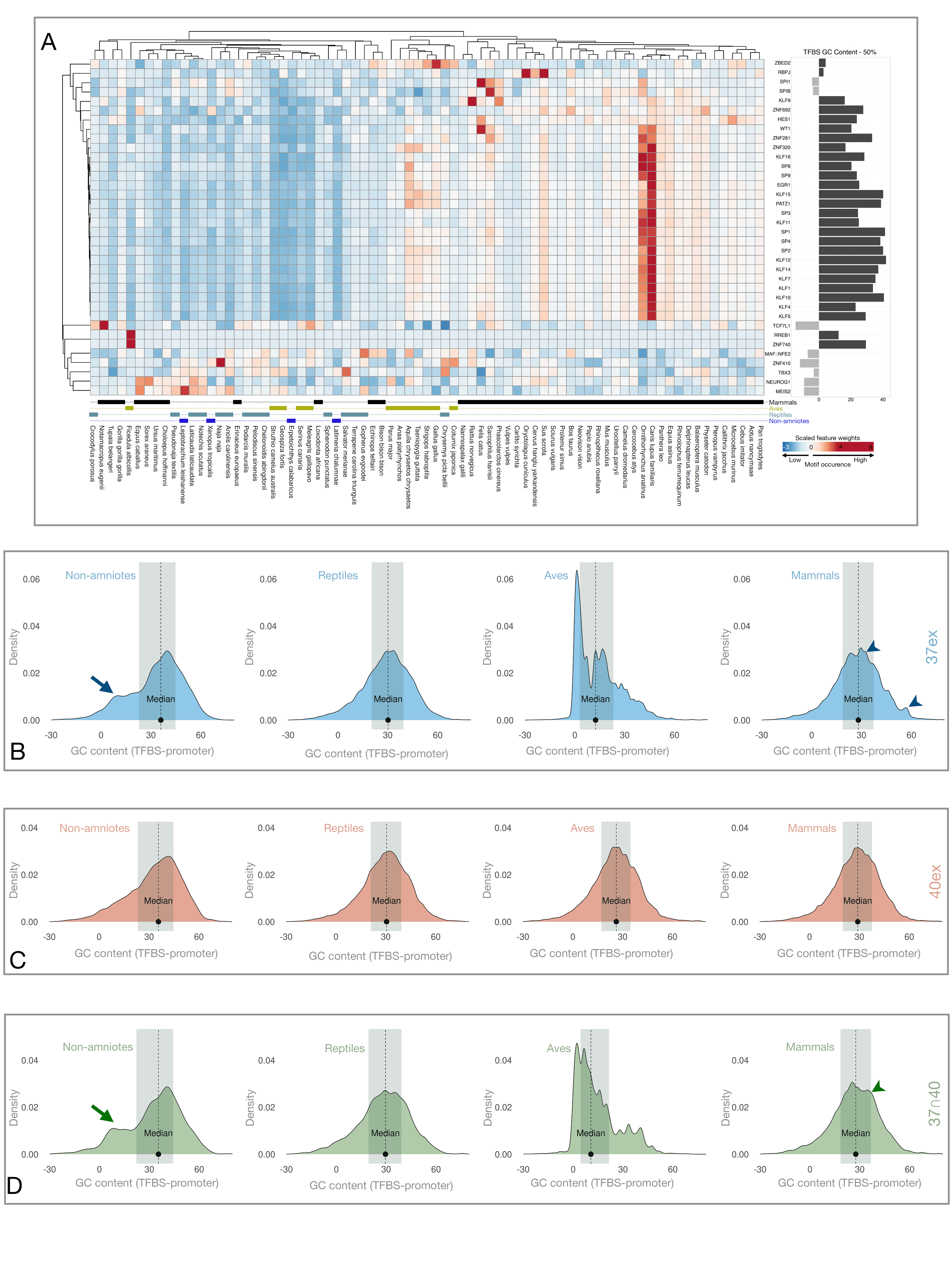
Promoters of 37ex ortholog genes show abundance and GC contents consistent with cytosine methylation protection by CGGBP1. A: Heatmap showing occurrence of 36 hypomethylated motifs in a heterologous set of 37ex-orthologous promoters from 75 vertebrate species selected by presence of CGGBP1 (non-amniotes 4, reptiles 14, aves 12 and mammals 45). The species names and taxon group assignments are indicated under the heatmap. The largest category of orthologs from mammals cluster together due to high occurrence of GC-rich TFBSs. GC-richness of the TFBSs is indicated in the panel to the right of the heatmap (dark gray and light gray respectively represent TFBSs with GC contents higher or lower than 50%). The clustering of mammals is primarily due to the abundance of the high GC content TFBSs only. B: The TFBS-specific GC-content shifts in promoters of 37ex orthologs show that the low GC content TFBSs in non-amniotes (blue arrow in non-amniotes) have shifted towards high GC content in amniotes with some lineage-specific variation in aves (high overall promoter GC content lowering the TFBS-promoter GC content differentials, Fig. S13B). In mammals, the taxon group with methylation restrictive CGGBP1 and highest abundance of these TFBSs, the GC-richness of a group of TFBSs resists lowering of TFBS-promoter GC content differentials (blue arrowheads in mammals). C: The specificity of the patterns of TFBS-promoter GC content differentials for 37ex genes (shown in B) was established by repeating the analyses for 40ex genes and their orthologous promoters. The 40ex genes do not show the TFBS GC content retention as discovered for 37ex genes. D: The same analysis for 37∩40 genes showed that the genes commonly repressed at 37°C and 40°C recapitulate the patterns of TFBS GC content retention observed for 37ex genes (green arrow in non-amniotes and arrowhead in mammals). A subset of the genes repressed at 40°C retain the promoter features of 37ex genes and these features are evolutionarily conserved in their orthologs highlighting the functional relevance of methylation restriction, TFBS GC content enrichment and transcriptional repression by CGGBP1. B-D: The dotted vertical line in all plots represents the median and the shaded region is the IQR.

Methylation states of promoters and TFBSs in such a large and disparate set of organisms is difficult to analyze experimentally. So we used the species specific retention of GC content in TFBSs as a surrogate for cytosine retention aided by methylation restriction. A targeted analysis of the CGGBP-regulated TFBSs in promoters of orthologs of CGGBP-repressed genes allowed us to assess how methylation restriction by CGGBPs in different species led to different levels of GC retention in TFBSs. We specifically analyzed GC retention in TFBSs over and above the background GC retention in the promoters harboring them by calculating differentials of GC contents between two different entities: the actual TFBSs and respective promoters. We established that this strategy avoided any false positive detection of GC-retention in mammals through a GC-rich k-mer assessment with or without the motif-promoter differential calculation (additional file 3). Using this method we calculated the differential GC content of TFBSs (details in methods) in the orthologous sequences from 75 species and compared them between four groups with different levels of CGGBP conservation; non-amniotes, reptiles, aves and mammals. For a control comparison set, we performed these calculations also for the hypomethylated TFBSs in all the known promoters (Ensembl) from 105 species which are a non-redundant superset of the species for which orthology data were available (Fig S13B). A relatively higher GC retention in TFBSs would yield a positive differential whereas a relatively high GC in promoters sequences would yield a negative differential. These calculations were normalized per unit length of the DNA sequence analyzed thus eliminating any artifacts due to different counts of orthologous sequences available and represented.

The GC differentials of the orthologous promoters were spread over a large range in all groups except aves (Fig 3, B and D). Specifically in non-amniotes a large subset of orthologous promoters showed a low GC bias in TFBSs, because these TFBSs were located in GC-rich promoters (Fig 3, B and D). The promoter GC contents increased in reptiles and mammals (Fig S13A) for several promoters but the TFBS contents did not decrease concomitantly. Aves however showed two different types of promoters one with extremely high GC-content and another with similar GC content as observed in reptiles and mammals and this was reflected in two different modes of TFBS-Promoter GC content differentials (Fig 3, B and D). The avian orthologs with similar GC content as those in reptiles and mammals appear to have resisted increase in genome-wide GC-content and associated changes in promoter GC-content and allow relatable comparisons across the taxon groups. Unlike these, the GC-rich avian promoters exhibited lineage specific variations over a large range of GC content differentials. Thus, the increase in TFBS GC content in amniotes has been higher than that at the promoters containing them, with some lineage-specific variations observed in aves (Fig S13B).

Such a maintenance of TFBS GC contents even when the promoter GC-contents change was not a generic phenomenon. Comparisons of all TSSs genome-wide showed that the TFBS- specific GC retention is restricted to a subset of all genes which are regulated by CGGBP1 or their orthologs. Also, this is a mechanism exerted by CGGBP1 at the native physiological conditions of 37°C (37ex) (Fig 3, B and D). By analyzing the genes repressed at 40°C (40ex) we did not observe any such shifts in promoter GC-contents (Fig 3C). Most interestingly, the strongest shift in promoter GC content resisted by the TFBSs was observed for a set of 470 genes (37∩40) which were repressed by Hs only at 37°C and by Hs as well as Gg at 40°C and remained non-repressed by Ac and Lc at either temperature (Fig 3D). These findings suggested that GC content of motifs with respect to promoters is a parameter relevant to repression by CGGBPs. It also shows that the same TFBSs which are methylation restricted by Hs are maintained at high GC content in relatively GC-poor promoters in non-amniotes compared to amniotes (Fig S13A).

Cytosine methylation restriction could affect C-T transition rates thereby maintaining C at TFBSs and give rise to high TFBS-promoter GC-content differences. If true, the higher G/C abundance in TFBSs would reflect as a T-C transition from non-amniotes to mammals. The CpG context allows confident measurements of cytosine methylation associated with pyrimidine transitions changing from CpG dinucleotides to TpG. We thus analyzed the hypomethylated TFBSs in CGGBP-regulated promoters for CpG to TpG transition rate differences between non-amniotes, reptiles, aves and mammals as a surrogate for differences in cytosine methylation profiles.

### CGGBP1-dependent methylation restriction at target gene promoters is predominantly at non-CpG sites with different effects on TFBS composition in amniotes and non-amniotes

We next analyzed the cytosine sequence context at which methylation restriction exerted by CGGBP1 leads to selective GC retention in TFBSs. CpG methylation leading to TpG is a telltale sign of cytosine methylation associated with base transition in the CpG context. Genome-wide, without restricting to the CGGBP1-repressed genes, the CpG contents of amniotic promoters were found to be clearly higher than those of the non-amniotes (Fig S13C). While CpG contents increased from non-amniotes to amniotes, TpG contents decreased slightly (Fig S13D). Put together, these results showed that methylation protection most likely plays a role in the enhanced CpG content in amniotic promoters that is the highest in mammals.

Previous studies have shown that CGGBP1 restricts cytosine methylation predominantly in the CHH context genome-wide with only weak effects on CpG methylation (6,7). However, the current analysis is concentrated at promoter sequences which are richer in CpG including comparisons between species with different levels of CpG content in promoters; mammalian promoters have progressively concentrated CpG dinucleotides (Fig S13C). In vertebrate promoters cytosine methylation is low in CpG context and methylation restriction by CGGBP1 can be exercised only in those sequence contexts wherein cytosine is prone to become methylated, that is non-CpG. To rule out the possibility of CpG methylation as a driver of C-T transition at promoters containing these hypomethylated TFBSs, we asked if (i) the promoter sequences used in our analyses conform to the expected patterns of CpG content change from non-amniotes to mammals, (ii) if there are any evidences of CpG to TpG transition rates indicative of cytosine methylation in promoter sequences and (iii) whether, of the hypomethylated TFBSs, any CpG containing ones, show retention of CpG over TpG. These questions were answered by analyzing the CpG-TpG transition rate differentials between TFBSs and respective promoters. This calculation also employed a differential of motif-promoter values to eliminate any false positive detections due to base compositional biases between different taxon groups (additional file 3).

The CpG-TpG differentials of the orthologs of 37ex, 40ex and 37∩40 genes showed taxon group-specific differences (Fig 4, A-D). In non-amniotes, the promoters of the orthologs of these genes showed a CpG-TpG differential pattern which was expected from the respective genome-wide patterns (Fig 4A). Two distinct groups of promoters were visible with different levels of CpG-TpG differentials genome-wide as well as in the 37ex, 40ex and 37∩40 groups. These findings showed that the non-amniotic orthologs of CGGBP1-repressed genes do not conform to distinct CpG methylation patterns different from the respective genome-wide patterns (Fig 4A). Interestingly, the non-amniotic representative CGGBP from Lc also failed to exert repression on these genes. While a cause-effect relationship between CpG-TpG differentials and gene repression is hard to establish, we analyzed if this association between CpG-TpG differentials and gene repression held true for all the forms of CGGBPs tested or not.

**Figure 4.**
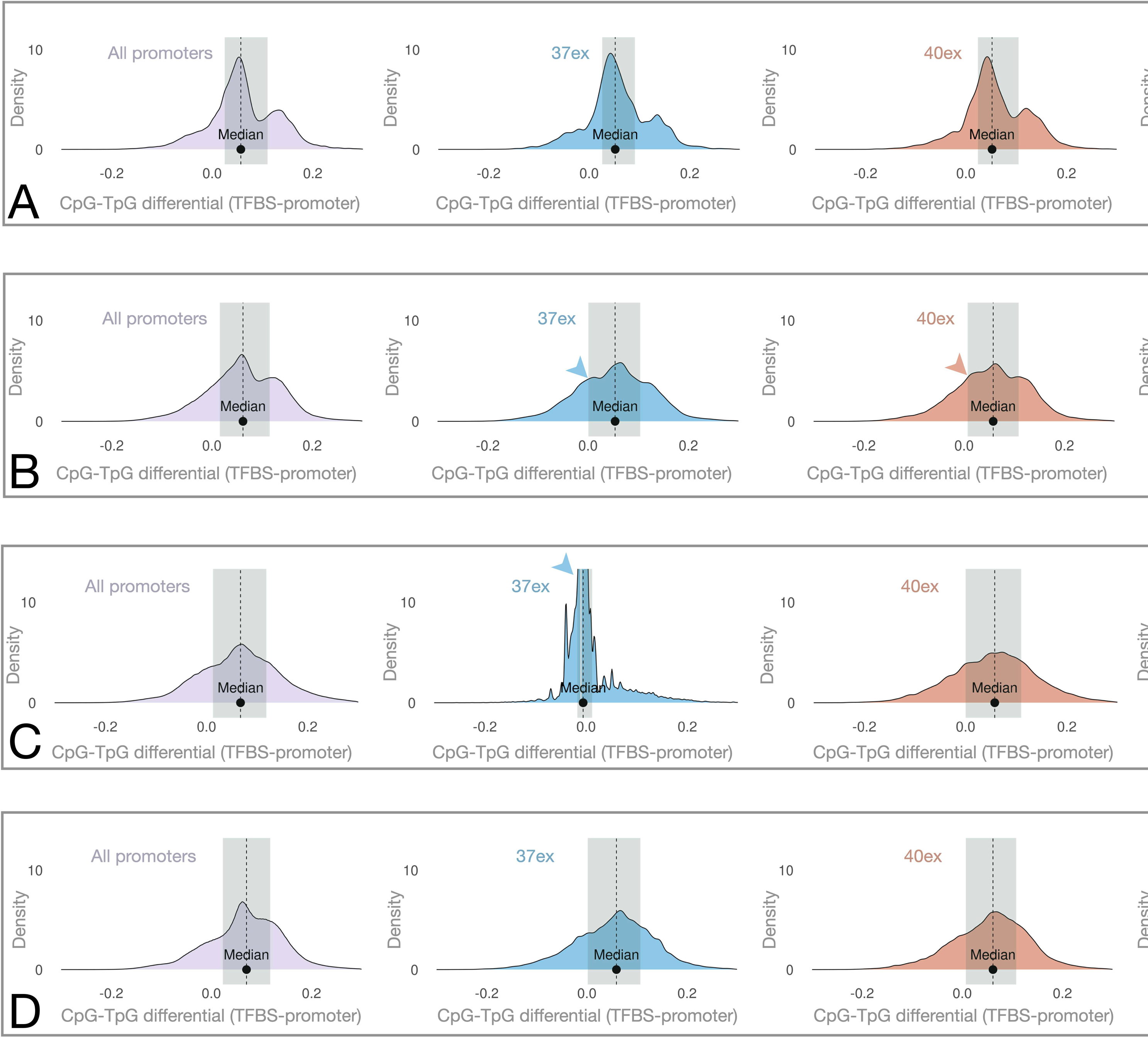
The methylation restriction at GC-rich TFBSs is independent of CpG methylation as estimated by CpG to TpG transition rates in orthologs of CGGBP1 repressed promoters against all promoters genome-wide in different taxon groups. A and B: The non-amniotic (A) and reptilian (B) orthologs of 37ex, 40ex or 37∩40 gene promoters show no major differences in TFBS-specific CpG retention patterns between expected (all promoters genome-wide) and observed (data from different temperature gene sets). The differences between all the analyzed promoter sets of non-amniotes and reptiles are due to broad changes on promoter CpG contents between non-amniotes and reptiles. The indicated (arrowheads) region of the CpG- TpG density distributions in reptiles (B) show that the reptilian orthologs of CGGBP1 represses gene promoters have a higher CpG retention not just in the TFBSs but in the promoters overall. C: In aves CpG retention was observed exclusively in orthologs of 37°C genes (blue and green arrows). D: In mammalian orthologs of CGGBP1-repressed genes no TFBS specific CpG retention was observed. The most consistently CGGBP1-repressed promoters however showed a consolidation of CpG-TpG transition rates to a narrow range, likely due to the high CpG richness of these promoters.

Like Lc, the reptilian representative CGGBP1 from Ac had also failed to repress these genes. The orthologs of these genes from 14 reptilian representatives showed a remarkably similar CpG-TpG differential distribution (Fig 4B); something unexpected due to the multiple lineage-specific diversities in reptiles. However, the 37ex, 40ex and 37∩40, all gene sets showed nearly identical distribution patterns of CpG-TpG differentials across the orthologs (Fig 4B). The reptiles are evolutionarily distant from non-amniotes; such a similarity of gene regulation and promoter CpG methylation pattern with non-amniotes was unexpected. Apparently, like the non-amniotic orthologs, the reptilian orthologs also seemed to escape any CpG methylation restriction of the promoters as well as repression of the associated gene as a linked phenomenon. If these associations between lack of repression and CpG methylation restrictions in non-amniotes and reptiles were not coincidental, one would expect the avian and mammalian orthologs to exhibit a CpG-TpG differential pattern compatible with the repressions observed due to Gg and Hs CGGBP1s respectively.

Promoters have become CpG-rich in the avian and mammalian lineages (Fig S13C). In these promoters if the hypomethylated TFBSs have resisted such an increase in CpG content, they would give rise to a decrease in the CpG-TpG differentials. Such a decrease of CpG-TpG differentials was observed in all the genes repressed by avian CGGBP1 except those which were repressed exclusively upon heat stress (Fig 4C). Thus, the avian CGGBP1 preferentially targets CpG-rich promoters for repression at 37°C of which only some retain repression at 40°C. This gene repressive function of avian CGGBP1 at CpG-rich promoters observed in human cells was not random. The avian orthologs of 1281 human genes repressed by Gg CGGBP1 obtained from 12 avian genomes showed a CpG retention pattern specifically for 37ex and 37∩40 promoters. CpG-rich promoters are generally poor in cytosine methylation (including at the TFBSs harbored in the promoters) and such promoters gave rise to the patterns we observed in Gg 37ex and 37∩40 (Fig 4C). In comparison, the mammalian CGGBP1s also showed a strong preference for repression of CpG rich promoters. Unlike the avian CGGBP1 however, the CpG rich promoters were repressed by mammalian CGGBP1 at 37°C (37ex) as well as 40°C (40ex) (Fig 4D). In both the cases the repression of CpG-rich promoters does not depend on TFBSs methylation harboring, best represented by the avian and mammalian (37∩40) genes and their orthologs (Fig 4, C and D). The CpG richness as a factor determining repression by CGGBP1 is a feature of aves and mammals. The differences in the consistency with which CpG rich promoters are repressed by these two forms of CGGBP1 in human cells under different conditions of heat stress (Hs CGGBP1 being more consistent than Gg CGGBP1) could partly be due to the differences in the CGGBP1 from Hs and Gg and in part due to constraints of inter-species differences between Gg CGGBP1 and the Hs cells as its non-native environment.

Unlike our prior observations that GC retention in hypomethylated TFBSs was specific to 37ex promoters, no such specificity was observed for CpG-TpG differentials. Moving from non-amniotes to mammals, the 37∩40 gene promoters clearly showed a trend for no difference in CpT-TpG transition rates between TFBSs and promoters. The enrichment for GC but not as much for CpG-TpG differentials indicated that non-CpG methylation might be responsible for C- T transition in promoters leading to low promoter GC content. The same pattern of CpG-TpG differentials were observed for 40ex gene promoters and their orthologs. Intermediates of these CpG-TpG transition rate differences were observed in reptiles and aves (Fig 4, B and C). These observations were aligned with the generally low cytosine methylation of CpG rich promoter sequences in mammals. It also became clear that the methylation restriction function of CpG is concentrated at non-CpG cytosines. These results suggest that (i) the orthologs of 37ex gene promoters in non-amniotes are under two different kinds of methylation regulation such that methylation protection offered by non-amniotic CGGBPs is limited only to a subset of the promoters, (ii) CpG-TpG transition rate differences between TFBSs and respective promoters for CGGBP1-repressed genes and their orthologs has undergone a change during evolution and stabilized with the emergence of mammalian CGGBP1, and (iii) the cytosine methylation restriction at TFBSs by CGGBP1 is a non-CpG phenomenon.

So far our analyses utilized gene sets fetched through a k-means clustering exercise as it was necessary to compare sequence properties from orthologs of the same gene set as much as available (Fig 1, D and F). As indicated by the results so far, the repression by CGGBPs, especially the mammalian CGGBP1 represented by Hs, is tightly linked to a methylation restriction of specific TFBSs. This methylation restriction is manifested as a higher non-CpG GC-content of the TFBSs as compared to the respective promoters. To further establish the robustness of association between of non-CpG GC retention in TFBSs at CGGBP1-repressed genes we compared GC and CpG-TpG differentials between TFBSs and promoters in different sets of genes consistently repressed by each form of CGGBP1 at 37°C in three replicate experiments. Unlike the gene sets derived through a k-means clustering, these genes would be independently derived for each CGGBP with no constraints of occurrence in multiple comparisons (Tables S18-S21). The GC differential analysis of TFBSs and promoters of these genes showed the most striking feature of mammalian CGGBP1. A separate cluster of GC-poor TFBSs was found in the independently derived genes from Lc, Ac and Gg CGGBPs whereas only Hs CGGBP1 showed no such presence of GC-poor TFBSs (Fig 5A). The large cluster of TFBSs with a higher GC content was commonly present for all these gene sets. This property of Hs CGGBP1 was restricted only to the genes repressed at 37°C. A similar analysis of genes repressed at 40°C (top 1000 genes with the lowest p-values are enlisted in supplementary data (Tables S22-S25) showed no evidence for methylation restriction and associated GC retention at TFBSs (Fig 5B). Consistent with the results presented above, the GC retention in TFBSs by Hs CGGBP1 was lost when the analysis was repeated on the same promoters (40°C) and TFBSs for CpG-TpG transitions reinforcing that the methylation restrictions and associated GC retention is a strong feature of Hs CGGBP1 in non-CpG context and is a feature of mammalian CGGBP1 which is exhibited to different extents by reptilian and avian CGGBP1 as well (Fig S14, A and B).

**Figure 5.**
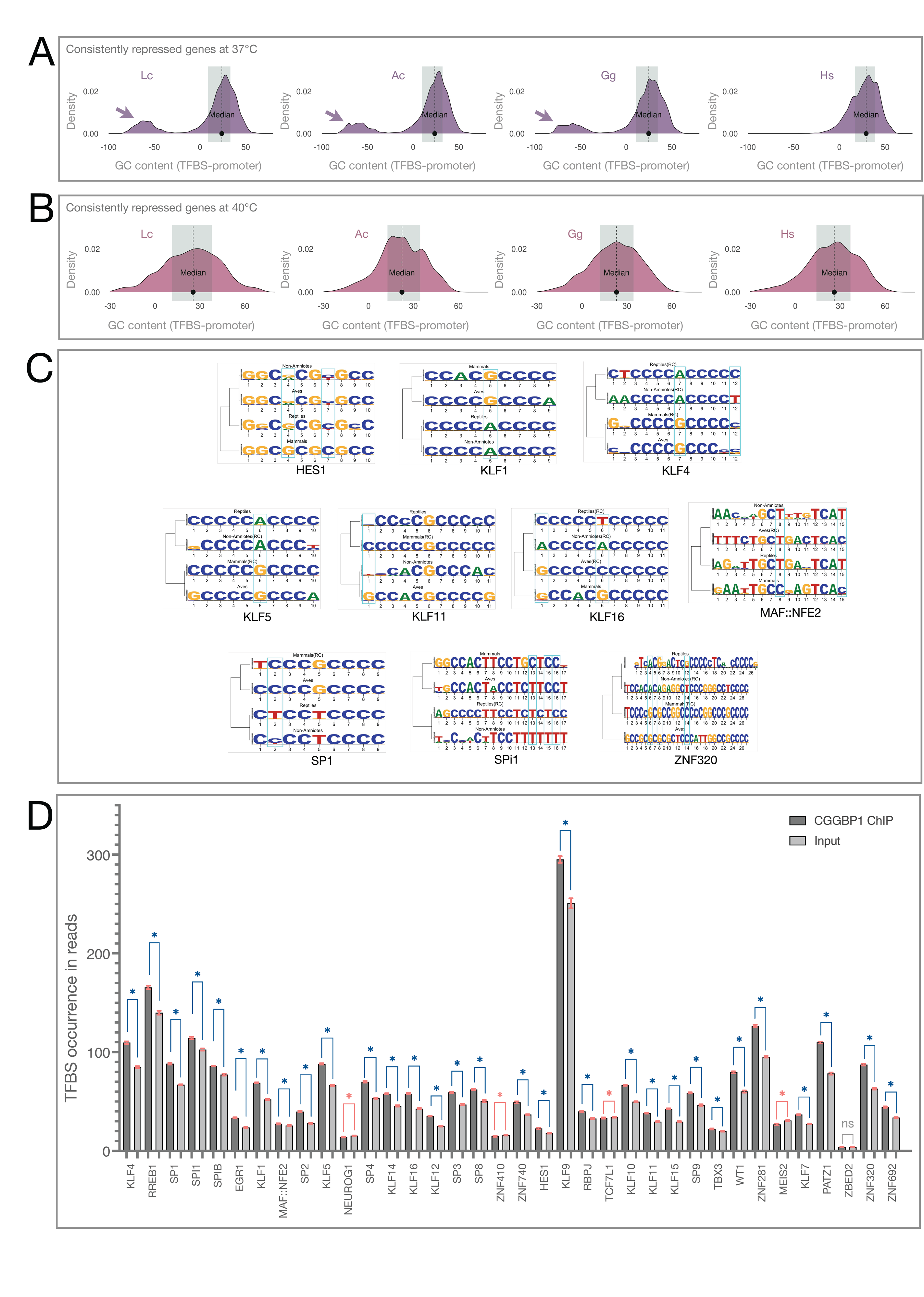
GC-rich TFBSs repressed by Hs CGGBP1 show evidence of lower C to T transition rates in mammals and enrichment in CGGBP1 ChIP-seq. A and B: Independent gene sets repressed by Lc, Ac, Gg and Hs reveal TFBS GC content bias as a feature of promoters repressed by Hs and mammalian CGGBP1 only at 37°C (A) not at 40°C (B). The low GC- content of Hs repressed promoters is contributed to by the absence of a gene cluster with GC- rich promoters in the Lc, Ac and Gg repressed promoters only (arrow heads). The Hs repressed promoters show a TFBS-driven high GC content differential (arrow) as compared to the non-Hs samples. The median values of the GC differentials in Lc, Ac and Gg are highly significantly different from those in Hs with all p-values <1e-40). C: Actual TFBSs with higher GC retention in CGGBP1 repressed gene promoters and their orthologs from various taxon groups show a G or C retention at specific positions in all amniotes (including reptiles) and in some cases in aves and mammals (higher amniotes). These specific instances were discovered by performing motif searches on actual TFBS strings in various orthologous sequences of CGGBP1-repressed genes against TFBS strings discovered in all promoters genome-wide (background). Specific instances of such C retentions, indicative of methylation restriction, are boxed. D: Of the 36 TFBSs with methylation restriction and associated GC retention, 29 are significantly enriched in CGGBP1 ChIP-seq (p-value < 0.01, blue asterisks). For four TFBSs there was a significant depletion in the ChIP-seq data (red asterisks) and for one TFBS there was no significant difference (gray “ns”). All comparisons were performed against input. Thus, CGGBP1 occupies and is physically associated with a vast majority of TFBSs exhibiting methylation restriction, GC enrichment and presence in CGGBP1-repressed promoters.

These taxon group-specific patterns of GC-content differentials between TFBSs and promoters from the orthologs of CGGBP1 target genes implied that different forms of CGGBPs might affect the TFBS composition differently. A strong methylation protection of the target TFBSs by mammalian CGGBP1, and to a lesser extent the avian CGGBP1, would favor emergence of GC-rich TFBSs. On the other hand a weaker methylation protection by non-amniotic CGGBPs would not favor TFBSs with GC-richness. As these TFBSs are representatives of protein binding sites pan-vertebrates, sweeping changes in TFBS composition are not expected. Any drastic changes in TFBS compositions would also be limited by the selection pressures operating through the effects of TF-TFBS bindings. So we expected to observe marginal but clear levels of GC enrichment in TFBSs between non-amniotes and amniotes. Some of these GC enrichments in TFBSs would indicate a cytosine methylation restriction. We specifically analyzed GC enrichments in the actual TFBS strings returned by FIMO (p-value < 0.00001) with or without indications of C-T transitions and compared them between non amniotes, reptiles, aves and mammals.

The differences in C-T transitions were qualitatively visible in 10 hypomethylated TFBSs (Table S26) using the promoter sets of the respective genome sets as backgrounds (Fig 5C). The position probability matrices from actual FIMO hits for the hypomethylated TFBSs from each taxon group and compared it against each other (motif alignments). We also found that for 7 motifs there were GC retentions (Table S27) without any indications of C-T transitions in TFBSs specifically in amniotes or mammals as compared to non amniotes (Fig S15). A simple inference from these findings was that in the non-amniotes and to some extent in early amniotes (reptiles) GC contents in TFBSs were favored by methylation restriction of cytosines thereby counteracting their loss through C-T transitions. In other cases where GC presence in amniotes was not likely to have occurred through a C-T transition, it could have been a methylation-independent GC-biased retention presumably due to an interaction between the TFBSs and CGGBP1. These inferences were supported by the observation that in none of the hypomethylated TFBSs there was any evidence of GC-loss in aves and mammals (Fig S15).

Such biased GC retentions could be favored by mitigation of methylation. There is evidence that CGGBP1 occupancy mitigates with cytosine methylation at many of these TFBSs (15). Methylation restriction by CGGBP1 has been shown to be dependent on its DNA-binding (15). It follows from our presented observations that if methylation restriction at the hypomethylated TFBSs is directly affected by CGGBP1 then we shall expect a specific occupancy of CGGBP1 at these TFBSs. By using publicly available CGGBP1 ChIP-seq data (GSE187851 and data available through PMID 39605320) we compared CGGBP1 occupancy at these TFBSs as enrichment against the input. Interestingly, the CGGBP1-binding regions identified in these experiments through peak-calling did not show enrichment of CG, CGG or CCG sequences (additional file 4). A *de novo* pattern search also did not reveal the presence of any CG or CGG or CCG-rich motifs (additional file 5), even if these are known in vitro binding sites for CGGBP1.

However, a read-level analysis using 1 million randomly drawn reads from these datasets over 10 iterations showed a mild yet highly significant enrichment of CG, CGG and CCG sequences (additional file 6). These analyses showed that CGGBP1 binds to a diverse set of sequences which interfere with pattern identifications in peaks but the properties of the sequences bound to CGGBP1 can be reliably analyzed by read-level comparisons between ChIP and respective inputs. We thus analyzed these data for subtle enrichment of the hypomethylated TFBSs at read level. The input and ChIP-seq sequences were subjected to FIMO search using the PWMs for the hypomethylated TFBSs. As additional controls ten different rounds of randomly sampled sequence reads were subjected to FIMO. As compared to the input we could observe a significant enrichment in CGGBP1-ChIP at these TFBSs (Fig 5D and Fig S16). A synthesis of these results is that CGGBP1 occupancy at these motifs restricts cytosine methylation and drives GC retention at the TFBSs. Through methylation restriction CGGBP1 plays a role in evolution of GC-biased TFBSs.

## Discussion

Cytosine methylation is a ubiquitous epigenetic mechanism implicated in gene silencing, heterochromatinization as well as genomic integrity. It is prevalent in genomes with varying levels of cytosines and although it is the most well understood in CpG context, there is increasing evidence for its functionally relevant occurrence in non-CpG context. There is mounting evidence that the same *de novo* methyltransferases which methylate cytosine in CpG context can also carry out cytosine methylation in non-CpG contexts (7,20–22). Cytosine methylation plays a pivotal role in the evolution of genomic regulatory elements by influencing methylation-associated base changes (6,8). This role of cytosine methylation has been widely investigated in the proximal promoters of RNA-Pol2 transcribed genes which are regulated by TFBSs. Cytosine methylation contributes to TFBS evolution in two different ways: Methylation facilitates C-T transitions thereby creating new motifs; also, methylation interferes with the binding of proteins to DNA thereby dissociating TFs from cognate TFBSs even without stably mutating the TFBS (23,24). The complexity of this phenomenon is enhanced by observations that in specific sequence contexts, especially CpG, methylation can facilitate DNA-protein interactions (25,26). A general theme which emerges from cytosine methylation in TFBSs is that stable retention of TFBSs would benefit from a lack of cytosine methylation. However, such a lack of methylation for the retention of TFBSs can not persist in conflict with the major gene repressive effect of cytosine methylation. Gene repressive mechanisms which prevent TFBS methylation are foundational to such a mechanism connecting cytosine methylation, TFBS evolution and gene repression.

Our recent works on the functions of a GC-rich DNA-binding and repeat-regulatory protein CGGBP1 have made the ground for the possibility that CGGBP1 has influenced TFBS evolution through cytosine methylation restriction. Unpublished results demonstrate that steric hindrances created by the occupancy of CGGBP1, dependent on its DNA-binding domain, negatively affect cytosine methylation at GC-rich TFBSs (15). This peculiar feature of CGGBP1 appears to exert its influence more predictably on TFBSs conserved in terrestrial vertebrates, especially amniotes. The differences in evolutionary variations between the N-terminal and C-terminal halves (15) are recapitulated using representative forms of CGGBPs from different vertebrate species. The N-terminal DNA-binding domain is relatively well conserved than the C-terminal part of CGGBP1 indicating that the DNA-binding-associated cytosine methylation protection is a feature that has resisted devolving in all the vertebrates sharing ancestry with the extant coelacanths.

Using high-throughput gene expression assays under the influence of overexpression of representative forms of CGGBPs from different evolutionary lineages, we find that gene repression is a common feature of CGGBPs that has been retained during evolution. The microarray-based gene expression assays demonstrate that there is an overall repressive effect of all forms of CGGBPs. Of the four forms tested, the least repressive effect was observed for the reptilian representative from *Anolis carolinensis*. This particular CGGBP1 has an anomalous N-terminal extension of the DNA-binding domain. Interpreted in the light of the findings of Morbia and colleagues (15), the DNA-binding of CGGBP1 seems important for repression by CGGBP1 through a mechanism that involves cytosine methylation restriction at TFBSs. In this investigation we have taken a non-candidate approach to independently assay cytosine methylation changes genome-wide by the same forms of CGGBPs as used for the gene expression assays. Neither the repressed genes nor the methylation changes brought about by alternative forms of CGGBPs showed any changes directed to achieve a particular function through regulation of a functional category of genes. Since these assays cannot be used to quantify global increases or decreases in transcriptional activity or cytosine methylation, we analyzed the qualitative features of these two datasets. We performed comparisons between the datasets obtained from different CGGBP forms to identify what DNA sequence features explain the differences in functions of the different evolutionary forms of CGGBPs. The experimental design and data analysis was tailored to define the current results in the light of the previous (4,9,18) and recent (15) reports of DNA sequence-dependent regulatory features of CGGBP1. The most relevant sequence-dependent feature of CGGBP1 is that various evolutionary forms of CGGBP have varying binding-site preferences. Although many of the binding sites identified *in vitro* for CGGBPs across most vertebrate lineages were GC-rich, there was a paucity of CpG dinucleotides. Another defining feature of CGGBP1 we factored in was the facultative nature of DNA-sequence dependence of CGGBP1 for gene repression. It has been previously reported that the CGGBP1-dependent repression of RNA Pol2 activity and occupancy in proximal promoters of target genes is abrogated upon heat shock. The TFBSs reported in these studies where CGGBP1 occupancy induced gene repression were derived from Alu-SINEs and consequentially GC-rich. Here we have used heat stress as an experimental condition under which we expected that any DNA-sequence dependence of gene repression by CGGBP1 would become redundant. It is important to note in this regard the non-detection of the human CGGBP1 in heat shocked samples; a finding previously reported to occur coincidentally in heat stressed samples with loss of a CGGBP1-repressive complex repressing HSF1 expression in a DNA sequence directed manner (18).

Global gene expression patterns under the influence of different evolutionary forms of CGGBP1 derived from independent experiments clustered Hs and Ac in two of the most distant groups. An underlying reason for this pattern of gene regulation could be that the Hs CGGBP1 is able to exert its repressive effect under the native conditions provided by the human cell whereas the non-human forms cluster away from Hs due to an incompatibility with the cellular milieu and resources to carry out gene repression. However, the anomalous N-terminal DNA-binding domain of Ac seemed to play a role in a relatively weaker repressive function as compared to the other forms of CGGBP1. Hence it is clear that even under the alien environment of a human cell the various non-human CGGBPs forms are able to exert transcriptional repression with some influence of their DNA-binding domain.

The incompatibilities between non-human CGGBPs and interaction partners of human origin can affect the functions of CGGBP1 which are not entirely DNA-sequence dependent. Those interactions between CGGBP1 and DNA which can occur directly and do not depend on the enabling interactions with other proteins could still materialize independent of the origin of CGGBP. This inference is based on the assumption that any PTMs required for DNA-binding of CGGBP1 take place in the human cells irrespective of the origin of CGGBP1. The N-terminal part of CGGBP1 preceding the DBD contains one such PTM site (Y20) necessary for CGGBP1 DNA binding and gene repression (1). The anomalous structure of Ac CGGBP1 in this part of the protein could explain why Ac CGGBP1 is the weakest repressor of gene expression. Our experiments capture repressive effects of various CGGBP forms which are not limited by interspecies incompatibilities between CGGBPs and interacting partners and these are likely to be direct DNA-CGGBP interactions. The DNA sequence preference would vary between different forms of CGGBP1 and the DNA sequence analysis of the promoters of CGGBP1- repressed genes supports this notion. The highest similarity in gene repression with respect to the repressive effects of Hs CGGBP1 in its native environment was observed for Gg CGGBP1, which amongst the representative CGGBPs has the lowest sequence difference from the Hs CGGBP1. Compared to Hs CGGBP1, with the highest sequence difference concentrated at its N-terminus flanking the DNA-binding domain, the Ac CGGBP1 was the least repressive. Also, upon expression of Lc CGGBP, which was the most different form of CGGBP compared to the Hs form, poor repression was observed. Such a dependence of gene repression on the extent of sequence difference from Hs CGGBP1, showed that the non-Hs forms of CGGBPs were not incapacitated in the human cellular context, nor were they toxic to the cells. It is important to note here that the only vertebrate lineage with no annotated CGGBP1 is that of the amphibians which are poikilothermic ectotherms. The only CGGBP in amphibians we could come across to employ in this study was from *Rhinatrema bivittatum*. We cloned and tried expressing it in HEK293T cells with no success. It is also noteworthy that the only poikilothermic and ectothermic CGGBP (Lc) maintained a very low expression level compared to other forms (Fig S3).

The expression data obtained from the 40°C experiments prove that the similarity between Gg and Hs CGGBP1 lead to highly similar gene repression patterns which stands in contrast with the lack of repression by Ac and Lc CGGBPs. The genes repressed at 40°C do not differ from those repressed at 37°C functionally and the only common feature identifying them which we have discovered is a pattern of the difference between GC content of the promoter and associated TFBSs. The abundance of GC-rich TFBSs in 37°C repressed genes but not in the 40°C repressed genes clearly show that different forms of CGGBPs target genes for repression specifically through GC-rich TFBSs in the 1 kb upstream promoters. These results are consistent with the proposition that the non-human forms of CGGBPs retain some of their gene repressive functions and continue to exert them specifically through regulation of GC-rich TFBSs. Interestingly, many of these TFBSs have been also identified as repressed by Hs CGGBP1 through its N-terminal DNA binding domain.

There are Hs-specific repressed genes but no Gg, Ac or Lc specific repressed genes.

The non-Hs forms might be unable to repress genes as the cellular environment and resources are alien and incompatible with respective CGGBP functioning. However, heat-stress removes such incompatibilities specifically for Gg, not Ac or Lc. Thus, CGGBP1 represses genes in a manner that is different at 37°C and 40°C conditions. At 37°C, taxon-specific differences between Hs and Gg manifest themselves.

Gene repression is a key function of CGGBP1 (6,15,18). Hs overexpression predominantly represses genes. Gene repression by Hs is shared only with Gg but not with Ac or Lc. Gene repression commonality between Hs and Gg increases at 40°C. Even under this condition Ac and Lc fail to repress genes. Evolutionary features of CGGBP1 in Gg and increasingly in Hs, enable them to act as gene repressors. Heat stress-responsive gene repression is a feature present in Gg and Hs both. Hence, it is either independently evolved in the two lineages or inherited from an ancestral form of CGGBP1 common to aves and mammals. But why these genes? These genes seem to be disparate with no commonality of functional categories. We tested if these genes are regulated by a common set of TFBSs highly prevalent in their promoter sequences. Through this analysis we found that although these genes are rich in a set of GC-rich TFBSs, these TFBSs are not restricted to these genes only. We concluded that the DNA sequence features of these promoters alone cannot explain the regulatory effects of different forms of CGGBP1 on their expression. As mentioned earlier the Hs CGGBP1- repressed genes are not only quantitatively rich in the hypomethylated TFBSs but are also compositionally rich in GC content, apparently due to restriction of methylation by CGGBP1. These orthologous promoter sequences allowed us to test some fundamental evolutionary properties of CGGBP1: Is methylation restriction a feature only of the human, and by extension mammalian, CGGBP1? What is the GC content profile of hypomethylated TFBSs in the orthologous promoters? Based on the premise that cytosine methylation restriction by CGGBP1 is a likely underlying mechanism for GC-richness of the hypomethylated TFBSs, we posited that TFBS GC contents in orthologous promoters would indirectly reveal cytosine methylation restrictive properties of CGGBPs from the respective species; a possibility that is very difficult to test experimentally.

One of the effects of CGGBP1 occupancy on GC-rich TFBSs could be a regulatable gene repression without constitutive silencing through cytosine methylation. Such a possibility has been shown to work in the repression of FMR1 gene through suppression of cytosine methylation in the CGG repeats (1–3,27,28). Evidence of such a phenomenon occurring genome-wide has been reported (6,7), including in some of our recent unpublished findings (15). Through an exhaustive combined analysis of the promoter sequences and cytosine methylation patterns of genes repressed by various forms of CGGBPs, their orthologs and their TFBS properties we have shown that the regulation of target promoters by CGGBP1 is not only in the form of gene repression rather also their regulatability by specific transcription factors through specific TFBSs. In addition to repression of promoters, CGGBP1 also contributes to the promoter sequence properties by preserving TFBSs through a restriction of cytosine methylation. A key feature of our orthologous sequence analysis is that we have imputed the effects of cytosine methylation associated mutations in TFBSs over and above the background changes in the promoter regions harboring them. These calculations fetch the effects of cytosine methylation restriction on evolution of the TFBS sequences themselves. CGGBP1 itself is not dependent on certain specific DNA sequence motifs and can bind to a variety of GC-rich DNA sequences. As such, the effects of CGGBP1 at the TFBSs, where the promoter functions are concentrated, are confounded by the effects at the rest of the promoter. The effects of CGGBP1 overexpression on TFBS methylation in the target promoters and associated GC content changes help resolve this. It can be inferred from our findings that the CGGBP1- repressed promoters have a significant amount of CGGBP1 occupancy. This occupancy duality between CGGBP1 and DNA keeps the methyltransferase activity under check, most likely sterically. At the TFBSs however the corresponding TFs would compete with CGGBP1 for occupancy. The stability of TF and TFBS complexes will depend on the actual TFBS sequence. Since a change in TFBS is a faster phenomenon with indirect influences on evolution of the TFs, it seems that CGGBP1 buffers this evolutionary process by preventing cytosine methylation at TFBSs more strongly than at the non-TFBS segments of the promoters. By preventing cytosine methylation CGGBP1 slows down the loss of C to T and as a complement, of G to A. The changes in specific TFBSs in the target promoters and their orthologs show clear instances of C-T transitions in the course of evolution which parallel the evolution of CGGBP1, especially its N-terminal DNA-binding part. Our recent work shows that GC-rich TFBSs in 1 kb promoters of CGGBP1-repressed genes are maintained as unmethylated through the DNA- binding of CGGBP1 (15). Apparently, the presence of N-terminal DNA-binding domain of CGGBP1 is sufficient to create steric hindrance to cytosine methylation at GC-rich TFBSs. The occupancy of the TFBSs by CGGBP1 is also likely to slow down the mutation rate in general and would reflect in the form of GC retention in ways other than those expected from a lack of cytosine methylation. We have observed several such instances.

It follows from our results that the methylation restriction of GC-rich TFBSs by Hs CGGBP1 is reflection of a larger process which might be occurring genome-wide which we only discover partially due to the limitations of our approaches. For instance, usage of RNA-sequencing and whole genome bisulfite sequencing could help us unveil the extensions of our currently presented findings to the TFBS evolution landscape of the entire genomes. A likely possibility is that CGGBP1 restricts cytosine methylation genome-wide and in turn allows retention of existing TFBSs through methylation prevention at specific GC-rich TFBSs. In achieving this, CGGBP1 also contributes to a phenomenon through which C-T transitions do not generate excessive levels of new TFBSs randomly which could be deleterious. Thus, CGGBP1 performs methylation restriction with a buffering effect on the TFBS evolution genome-wide.

Overexpression of CGGBPs in different forms just allows us to focus on specific genes and their promoters purely due to the sequence properties of their promoters. As such, these genes are no representation of the fraction of the genome to which CGGBP1 function is directed. By repressing transcription and at the same time suppressing methylation CGGBP1 creates a system of transcription repression which is not only less mutable, but also malleable and not as rigid as cytosine methylation. That CGGBP1 quickly loses its preference for the target promoters upon heat stress demonstrates this. The evolution of this function of CGGBP1 in the course of vertebrate evolution suggests that with the increasing stressful terrestrial habitats the retention of critical TFBSs in target promoters were an adaptive need to which CGGBP1 has contributed by enabling a mechanism of gene repression which prevents TFBS loss and at the same time is amenable to a rapid change upon heat stress, a common stressor in the terrestrial habitats. Through such functions the role of CGGBP1 in the evolution of various adaptive features of terrestrial vertebrates, including homeothermic endothermy, can not be discounted. The emergence of CGGBP1 and its evolutionary conservation in amniotes, and progressively more so in homeothermic endotherms and mammals only, supports this idea.

## Supporting information

Raw data file

## Limitations of the study

The cross-species comparisons conducted in this analysis do not encompass all potential species combinations. Additionally, the methylation data presented lacks base-level resolution.

## Ethics approval and consent to participate

Not applicable.

## Consent for publication

Not applicable.

## Resource availability

The data have been deposited at NCBI GEO (private).

## Authors’ contributions

PK and IM performed experiments, PK, IM, ALS and US analyzed the data, SD helped with experiments and data analysis, PK and ALS prepared the figures, PK and US wrote the manuscript. All authors reviewed the manuscript.

## Acknowledgement

The authors acknowledge Dr. Dhiraj Bhatia (IIT Gandhinagar), Dr. Noopur Thakur (Ahmedabad University) and Dr. Amit Mandoli (NIPER Ahmedabad) for helping with various experiments.

## Supplementary figures and legends

Fig. S1. CGGBP1 knockdown and sub-cellular localization of its various forms in HEK293T cells. A: shRNA-mediated knockdown of human CGGBP1 in HEK293T cells targeting four distinct regions of the ORF, as detailed in the Methods. HEK293T control (CT) cells express endogenous CGGBP1 at 20 kDa, while HEK293T knockdown (KD) cells show a reduction of CGGBP1 levels to approximately 70-80%. Beta-actin (42 kDa) was used as a loading control. B: Nuclear-cytoplasmic fractionation assays reveal that all vertebrate forms of CGGBP1 are predominantly localized to the nucleus after being synthesized in the cytoplasm, with little to no CGGBP1 detected in the cytoplasm. β-actin was used as a cytoplasmic marker, while total Histone H3 served as a marker for the nuclear fraction. Lc, Ac, and Gg CGGBPs were detected using a FLAG-tag antibody, and Hs CGGBP1 was probed with an anti-HA antibody. The lanes were rearranged to ensure proper alignment for β-actin. To account for the differing amounts of sample, the nuclear fraction was loaded at half the volume (7µl) compared to the cytoplasmic fraction (15µl), which revealed variations in expression levels between the samples.

Fig. S2: Global gene expression changes caused by expression of the indicated forms of CGGBP1 (Lc, Ac, Gg or Hs) when compared against the pattern expressing an empty vector (Ev) at 37°C. In the samples representative of non-amniotes (Lc) and lower amniotes (reptiles) widespread gene deregulation were observed with spikes of derepressed genes (visible as positive M-value spikes and modes of the density distribution at the top. In the representatives of higher amniotes (Gg and Hs) however the larger change in expression was repression. The extent of gene deregulation decreased with similarity between the forms of CGGBP1 representatives and the taxa they represent. Consistently differentially expressed genes (p- value < 0.01 in three replicate experiments) shown as green dots. The Y-axes show -log10 p- values and the X-axes show M values were calculated using the geometric means of the three replicate experiments.

Fig. S3: CGGBP1 overexpression under heat stress conditions. HEK293T cells were transfected with various CGGBP1 constructs and exposed to heat stress at 40°C for 48 hours (details in methods). Western blot analysis revealed multiple bands corresponding to different molecular weights of CGGBP1, aside from the typical 20 kDa species, as marked by red arrows, indicating potential heat-induced modifications such as ubiquitination. Non-human (non-Hs) CGGBP1 isoforms were detected using anti-FLAG antibodies. In contrast, human CGGBP1 (Hs CGGBP1) could not be detected with the HA-tag antibody, as reported previously. However, using a polyclonal CGGBP1 antibody, we identified Hs CGGBP1 running at a higher molecular weight than expected, while the conventional 20 kDa form was absent, despite using multiple antibodies. The polyclonal antibody also detected non-Hs CGGBP1 due to sequence similarity. Beta-actin was used as a loading control. CGGBP1 forms exhibit differential expression levels, with the non-amniotic form Lc being undetectable with the FLAG-tag unless the expression of the other samples reaches saturation, as indicated by the blue arrow.

Fig. S4: Global gene expression changes brought by the indicated forms of CGGBPs at 40°C. A-D: The global deregulation of gene expression was widespread at 40°C with no clear directions of deregulation and differences between species forms of CGGBP1 as observed at 37°C (Fig S1). Consistently differentially expressed genes (p-value < 0.01 in three replicate experiments) shown as green dots. The Y-axes show -log_10_ p-values and the X-axes show M- values were calculated using the geometric means of the three replicate experiments.

Fig. S5: Comparisons of gene expression changes caused by heat stress in the presence of various different forms of CGGBP1 reveal gene repression under heat stress as a feature of CGGBP1 from higher amniotes (Gg and Hs) not present in lower amniotes (Ac) or non-amniotes (Lc). Only consistently deregulated genes (p-value < 0.01 in 37°C versus 40°C comparisons) are plotted on the Y-axes and from Ev on the X-axes. Quadrants: Q1- significant induction in Ev as well as the indicated sample, Q2- significant repression in the indicated sample but induction in Ev, Q3- significant induction in Ev as well as the indicated sample, and Q4-significant repression in Ev but induction in the indicated sample. Red arrowheads mark the regions showing data points where significantly differentially expressed genes upon heat stress have opposing directions of change as compared to Ev. Lc and Ac show abnormal induction and Gg shows abnormal repression.

Fig. S6: An overall summary of MeDIP-seq. A: Frequency distribution of MeDIP signals in 0.2kb bins genome wide. Various samples are represented by different colors (Lc - brown, Ac - purple, Gg - red, Hs - green, Ev - orange and Input - Blue). All samples show enrichment over input across a wide range of signals. Hs mitigates cytosine methylation and shows a frequency distribution shift towards lower methylation signals whereas Gg follows the distribution pattern of Ev. Overall Hs and Gg restrain methylation genome-wide. In contrast, Ac and Lc expression increase methylation and the distribution shifts towards higher signal bins. B: PCA of MeDIP signal genome wide shows that Ev, Hs and Gg are more similar and cause similar patterns of methylation while Lc and Ac cluster together due to the similarity in methylation pattern caused by their expression.

Fig. S7: Overview of MeDIP signals on regulatory elements and promoters. A-D: Input shows no enrichment of methylation and all samples show enrichment of methylation as compared to input. MeDIP signals on conserved regulatory elements (CREs) (A), enhancers (B), all TSS (robust and permissive) (C) and the promoters (D) of probe set on the microarrays show a similar pattern of methylation. The genomic regions are sorted in descending order of MeDP signals.

Fig. S8: MeDIP reads show no major differences in presence of JASPAR (vertebrate) TFBSs. The Y axis represents TFBS occurrences (log scale) fetched through FIMO search on randomly sampled 1 million MeDIP seq reads from the indicated samples. Each TFBS is paired between different samples by horizontal gray lines.

Fig. S9: MeDIP signal on of 37ex, 40ex and 37∩40 gene promoters. A: Methylation in 1 kb upstream region of 37ex promoters recapitulates gene expression pattern at 37℃. The methylation pattern on these genes sets Hs apart from other samples and clusters Gg and Ev into a separate cluster while Lc and Ac cause a similar pattern of methylation. B and C: Such a strong distinction of methylation pattern is not manifested at 40ex as well as 37∩40 gene promoters. These methylation patterns closely resemble the gene repression patterns caused by Lc, Ac, Gg and Hs (Figure 2D-E).

Fig. S10: Correlations between methylation patterns at TFBSs in 1 kb promoters of genes repressed by CGGBP1 at 37∩40 gene promoters show no specific pattern which sets apart TFBSs methylation pattern induced by changing various forms of CGGBP1. It appears that these specific genes which flip for Gg when the heat-stress in play are moderately dependent on TFBS methylation of some if not all TFBSs.

Fig. S11: Analysis of TFBS occurrence and density in gene promoters. A: The analysis of TFBS within the 1 kb region upstream of the 37ex, 40ex, and 37∩40 gene promoters show no significant variation in their frequency. B: The density of TFBS per kilobase in these specific gene sets is not different from that of a randomly chosen set of genes on the microarray. C: The TFBS occurrence in the 37ex, 40ex, and 37∩40 gene promoters closely match the overall TFBS density observed in all genes on the microarray.

Fig. S12: A survey of GC contents of all known promoters in the species used for orthology analysis. A: In the genome-wide promoter sets known for four non-amniotes, 17 reptiles, 27 aves and 57 mammals shows that most promoters are in the GC content range of 30% to 70%. The analysis of effects of different forms of CGGBP1 on TFBS methylation in repressed gene promoters and their orthologs were restricted to this set of promoters in the 30% to 70% range for eliminating the outlier effects. B: The occurrence of the 36 hypomethylated TFBSs in the promoters genome-wide is higher in mammals and aves and clusters majority of them together. The lower amniotes (reptiles) and non-amniotes have lower abundance of these TFBSs with high variability. These hypomethylated TFBSs are also majorly GC-rich and the larger cluster of the rows classifies the high GC-content TFBSs together. It is the high GC-content hypomethylated TFBSs which cluster higher amniotes differently from the non-amniotes and reptiles.

Fig. S13: Patterns of GC, CpG and TpG contents in the promoters and TFBSs of all genes from 105 species used in the analyses. A: Frequency distributions of the promoter GC contents show that the GC content of promoters have increased in amniotes (top panel) with strong increases in higher amniotes. B: The GC content increase in amniotes is accompanied by a clear increase in hypomethylated TFBS GC contents over and above the promoter GC contents (calculated as TFBS GC content-promoter GC contents). The TFBS-Promoter GC content differentials are bimodal in amniotes with a conspicuous increase in the mode with higher differential value (X- axis). C: Similar to the increase in TFBS GC content, the CpG content has also increased in amniotes with strong CpG enrichment observed in aves and mammals. D: The CpG content increase in TFBSs in higher amniotes (C) is not accompanied by any discernible directional changes in TpG contents. These statistics show that the hypomethylated TFBSs have evolved a higher GC content in higher amniotes, of which a small part could be contributed to by lower rates of CpG to TpG transitions, a possibility we have tested further on.

Fig. S14: The significantly differentially repressed genes by Lc, Ac, Gg and Hs in replicate experiments show CpG-TpG differential profiles very different from the corresponding GC differential profiles (Fig 5A and B). A: The CpG-TpG differentials of hypomethylated TFBSs compared to the promoters in genes repressed at 37°C (n=3 replicates, p-value < 0.01). B: The CpG-TpG differentials of hypomethylated TFBSs compared to the promoters in genes repressed at 37°C (n=3 replicates, p-value < 0.01). Noticeably, only Hs, and to some extent Gg, showed a CpG-TpG differential pattern in 37°C repressed promoters which resembled that observed in 40°C repressed promoters. Unlike GC content in the TFBSs, these results indicated that the CpG content of the TFBSs is relevant to such a robust repression by Hs CGGBP1 that it is independent of the heat stress.

Fig. S15: Examples of GC retention in TFBSs in CGGBP1-repressed promoters and their orthologs which can not be explained by restriction of cytosine methylation. These findings suggest that cytosine methylation restriction is not the only possible mechanism contributing to GC retention in TFBSs under the influence of CGGBP1 of higher amniotes.

Fig. S16: The 36 hypomethylated TFBSs with methylation restriction and associated GC retention provided by CGGBP1 (28) were significantly enriched (p-value < 0.01, blue asterisks) in the publicly available CGGBP1 ChIP-seq dataset (PMID: 39605320). Seven TFBSs showed significant depletion (red asterisks), and one TFBS showed no significant change (grey ‘ns’). Comparisons were made against the input, using 10 independent read samplings (n = 1 million).

## Supplemental information

Supplementary figures: Fig S1-S16 in one PDF.

Supplementary tables: One compressed file containing Table S1-S15 and S17-S27 as Excel files, and, supplementary table S16 as a PDF.

## Additional supplementary file

One compressed file containing information about CGGBP1 constructs as a PDF and lists of orthologous genes as text files.

## Additional files

Additional files 1 to 6 are submitted as individual PDF files.

## Methods

### CGGBP transgenic forms constructs

Vertebrate-specific CGGBP constructs representing *Latimeria chalumnae* (Lc), *Anolis carolinensis* (Ac), *Gallus gallus* (Gg), and *Homo sapiens* (Hs) were cloned into the constitutive eukaryotic expression vector pcDNA3.1(+) acquired from Addgene using *Kpn*I and *Xho*I restriction enzymes. The protein sequences were obtained from UniProt, and the coding sequences (CDS) were analyzed for codon optimization. The following Uniprot IDs correspond to the selected CGGBP forms: Q9UFW8 (Hs), A0A1D5NUG1 (Gg), G1KEH3 (Ac) and H3AJH5 (Lc). Each construct included *Kpn*I and *Xho*I restriction sites at the 5’ and 3’ ends, respectively. An N-terminal HA-tag was added to the Hs CGGBP1 construct described elsewhere (29), while C-terminal FLAG-tags were incorporated for the non-mammalian constructs (Lc, Ac, Gg). The cloned plasmid constructs underwent sequencing and were confirmed for fidelity through analysis. Comprehensive details of the sequences can be found in the supplementary materials and additional files.

### Cell culture, shRNA transduction for knockdown

HEK293T cells were cultured in DMEM (AL007A) augmented with 10% FBS and grown at physiological temperature (37°C). CGGBP1 knockdown was done using CGGBP1-shRNA (targeting four different regions in the ORF) using lentiviral transduction by overexpression of CGGBP1-lentivirus constructs. The lentiviral constructs and packaging plasmids were mixed in equimolar concentration and transfected for the production of lentiviruses. Control - HEK293T (CT) and CGGBP1-shRNA transduced cells - HEK293T (KD) were selected using puromycin 1 μg/ml (Himedia) for stable transduction. The lentivirus constructs were acquired from Origene. The lenti-packaging plasmids: pRSV-Rev (12253), pMDLg/pRRE (12251) and pMD2.G (12259) were obtained from Addgene. Transfection was accomplished using JetOptimus (Polyplus #101000051). Polybrene (Sigma #TR-1003) was used for transducing cells.

### Over-expression and heat-stress

To assess the stability and expression of vertebrate forms of CGGBP, we conducted a study using HEK293T cells. First, the cells were transfected and incubated at a physiological temperature of 37°C. We allowed the transfection to proceed for 72 hours to ensure sufficient expression of the transgene. For the heat shock experiment, 24 hours after the initial transfection, during which we overexpressed various forms of CGGBP1, the transfected cells were incubated at 40°C for 48 hours.

### Nuclear cytoplasmic fractionation assay

HEK293T(KD) cells were utilized for the overexpression of transgenic CGGBP variants (Lc, Ac, Gg), whereas HEK293T(CT) cells were employed for Ev and Hs overexpression. Following cell pellet collection through centrifugation, the pellets were resuspended in 1x Cytoplasmic extraction buffer, which included 10 mM HEPES, 60 mM KCl, 1 mM EDTA, 0.075% v/v NP-40, 1 mM DTT, and 1 mM PMSF. The buffer was supplemented with 1x Halt Protease Inhibitor Cocktail - EDTA-free (Thermo Fisher Scientific #87785) and mixed using cut tips before a 2- minute incubation on ice. The resulting mixture was centrifuged at 1200 rpm to isolate the supernatant, which constituted the cytoplasmic fraction. The pellet was then washed with a 1x washing buffer (10 mM HEPES, 60 mM KCl, 1 mM EDTA, 1 mM DTT, 1 mM PMSF) to remove any remaining cytoplasmic material. Following this, the pellet was treated with a 1x nuclear extraction buffer (20 mM Tris HCl, 420 mM NaCl, 1.5 mM MgCl_2_, 0.2 mM EDTA, 1 mM PMSF, 25% (v/v) glycerol), with the NaCl concentration adjusted to 400 mM. The nuclei were vortexed and incubated on ice for 10 minutes with intermittent vortexing, producing the nuclear fraction. Both fractions underwent centrifugation to clear debris and were subsequently analyzed by SDS-PAGE and Western blotting.

### Western blot

The samples were subjected to electrophoresis using a 4% stacking gel and a 10% resolving gel, with subsequent transfer to a PVDF membrane. The membrane underwent a blocking step for 1 hour at room temperature using a blocking buffer of 5% (w/v) dry milk in 1x TBST. Primary antibody incubation was conducted either overnight at 4°C or for 1.5 hours at room temperature at a dilution of 1:1000 in blocking buffer. Following a wash with 1x TBST, the membranes were incubated with an HRP-conjugated secondary antibody (1:5000 dilution in blocking buffer) for 1 hour at room temperature, followed by another wash with 1x TBST. Signal development was carried out using an ECL substrate, and the resulting blots were imaged in chemiluminescence mode on the BioRad ChemiDoc MP Imaging System (BioRad #12003154).

### Antibodies

The following antibodies were used for establishing the expression of various forms of CGGBP at 37°C and 40°C, as well as for the nucleo-cytoplasmic fractionation assay: Invitrogen Mouse GAPDH Loading Control Monoclonal Antibody (GA1R, Catalog #MA5-15738), Proteintech Rabbit CGGBP1 Polyclonal Antibody (Catalog #10716-1-AP), Proteintech Mouse Anti-DYKDDDDK Tag Monoclonal Antibody (Catalog #66008-4-Ig), Proteintech Mouse Anti-Beta Actin Monoclonal Antibody (Catalog #66009-1-Ig), Invitrogen Rabbit Monoclonal CGGBP1 Antibody (Catalog #PA5-57916), Santa Cruz Biotechnology (SCBT) Mouse HA-Probe (F-7) Monoclonal Antibody (sc-7392, Catalog #E1718), and SCBT Mouse Histone H3 (1G1) Monoclonal Antibody (sc-517576, Catalog #H3018).

### Microarray-based gene expression analysis

Transgenic constructs of CGGBP (Lc, Ac, Gg) were overexpressed in HEK293-T(KD) cells, while HEK293-T(CT) cells were utilized for the overexpression of Ev and Hs. Each experiment was conducted in triplicates. Cell pellets were snap-frozen and subsequently stored at −80°C for RNA extraction, which was performed using Trizol, followed by purification with the Qiagen RNeasy mini kit (Qiagen #74106). The Agilent Quick-Amp labelling kit (Agilent #5190-0424) facilitated T7 promoter-based linear amplification, generating Cy3-labeled complementary RNA for the gene expression analysis. Hybridization was conducted using Agilent’s In Situ Hybridization kit (Agilent #5188-5242) on the Human GXP 8X60k microarray chip (AMADID: 072363). Data normalization included intra-array signal normalization through GeneSpring GX 14.5 Software, along with intra-group quantile normalization for triplicates across the five samples (Lc, Ac, Gg, Hs, Ev), followed by total intensity normalization prior to data analysis. A paired t-test was conducted to assess the differences in gene expression between transgenic forms induced by Hs and Ev across various comparisons. For each gene, the geometric mean was calculated from three replicates, and the fold change in gene expression was determined relative to Hs or Ev. A threshold of p-value < 0.01 was established to identify consistently deregulated gene sets. To mitigate the stochastic effects of gene expression at 40°C, differentially expressed genes were ranked based on their p-values, with the top 1000 deregulated genes selected for further analysis. To compare gene expression at 37°C and 40°C, two analytical approaches were utilized. First, an unsupervised clustering of post-normalized data through k-means clustering was used to delineate the overall expression patterns. We then retained 37ex (811), 40ex (1512) and 37∩40 (470) genes for further analyses. Second, the qualitative assessment of species-specific changes induced by CGGBPs at 40°C compared to 37°C was achieved by calculating the respective sample fold changes for various comparisons.

### Methylcytosine DNA immunoprecipitation (MeDIP)

Transgenic variants of CGGBP (Lc, Ac, Gg) were overexpressed in HEK293T (KD) cells, while HEK293T (CT) cells were utilized for the overexpression of Ev and Hs. After 72 hours of transfection, cellular lysates were harvested for genomic DNA extraction following the phenol-chloroform-isoamyl alcohol protocol. Sonication was optimized to produce DNA fragments of uniform size, specifically between 700 and 1000 base pairs. One microgram of the fragmented genomic DNA underwent end repair using the NEBNext Ultra II End Repair/dA-Tailing Module (NEB #E7546). Adapter ligation was performed by attaching Oxford Nanopore Technologies PCR adapters (ONT #SQK-PSK004) to the end-repaired DNA with NEB Blunt/TA Ligase Master Mix (NEB #M0367). The adapter-ligated DNA was treated with a 1x MeDIP master mix, which included 10 mM sodium phosphate buffer, 0.14 M NaCl, and 0.05% Triton X-100. This mixture was denatured at 95°C for 5 minutes and subsequently snap-chilled on ice. An antibody cocktail specific to 5-methylcytosine (EMD Millipore #MABE146, Sigma #SAB2702243, and Novus Biologicals #NBP2–42813) was added and incubated overnight at 4°C. Following this, Protein G Plus/Protein A Agarose beads (Merck Millipore #IP05) were added and incubated at room temperature for 2 hours. After centrifugation, the beads were washed three times with the 1x MeDIP master mix and digested with Proteinase K (Sigma #P2308) at 56°C for 2 hours. The supernatant containing the immunoprecipitated DNA was collected and amplified through 18 PCR cycles using whole genome primers (ONT #SQK-PSK004) and LongAmp Hot Start Taq 2X Master Mix (NEB #M0533). The final library was prepared with the NEBNext® Ultra™ II DNA Library Prep Kit for Illumina® (NEB #E7645, #E7103), and sequencing was carried out on the Illumina HiSeq 2500 platform.

### MeDIP-seq analysis

Our analysis commenced with adapter trimming and quality filtering, which were performed using fastp. The resulting clean reads were then aligned to the unmasked hg38 genome using Bowtie2, focusing specifically on reads longer than 100 bp for further evaluation. We used bigWig files of the data with deepTools to generate heatmaps and profiles that assess the overall distribution and identify common regions across samples. For the MeDIP data analysis, we employed bedtools to calculate signal distributions within 0.2 kb bins across multiple samples. Broad peaks were identified using MACS2 without any background correction. Finally, we used the Integrative Genomics Viewer (IGV) to visualize the genomic tracks of the MeDIP- seq data.

### In-depth comparative analysis of promoter sequences across 105 vertebrate species

We downloaded 105 vertebrate genomes from the Ensembl Genome Browser (version 112), along with their corresponding annotation files as Gene Transfer Format (GTF) files. To account for taxonomic diversity, our analysis included 4 non-amniotes, 17 reptiles, 27 aves, and 57 mammals. From the GTF files, we extracted the coordinates of transcription start sites (TSS) located at exon 1. For each species, we retrieved FASTA sequences corresponding to 1 kb upstream regions (promoters) of these TSS. The GC content of the promoters was calculated individually, and the promoters were categorized into 10 bins based on GC content, ranging from 0% to 100%. The percentage of promoters in each bin was computed as a fraction of the total promoters for each species and visualized (see supplementary figure). Notably, the majority of promoters across representative species fell within the GC content range of 30% to 70%. Consequently, we focused our further analyses on these promoters. To investigate the presence of 36 hypomethylated transcription factor binding sites (TFBSs) within these promoters, we utilized FIMO (Find Individual Motif Occurrences, MEME Suite). Each FIMO output row was tagged with the corresponding promoter FASTA sequence. The results for the GC content bins of 30% to 70% were concatenated for each species. We applied a stringent threshold (p-value < 0.00001) to refine the TFBS discovery. Additionally, we calculated and visualized the occurrence of the 36 TFBSs across different taxa using ClustVis, a web tool designed for clustering multivariate data.

### Comprehensive characterization of promoter sequences in 75 orthologous species

The identification of orthologs presented a significant challenge across the 105 species studied; we successfully identified orthologs for 75 species, comprising 4 non-amniotes, 14 reptiles, 12 aves, and 45 mammals. The heterogenous orthologous gene sets for 37ex, 40ex and 37∩40 lists can be found in additional supplementary files. Orthology information was manually retrieved from the Ensembl BioMart database for the human gene dataset (GRCh38. p2) for the available species within each taxon. Each raw orthology file included Gene ID, Transcript ID, their respective versions in humans, and the corresponding pairwise ortholog mapping. We retained only those entries for which complete information was available, subsequently eliminating duplicate entries by transcript ID versions for human genes. Utilizing the transcript IDs from specific gene sets (37ex, 40ex, and 37∩40), we extracted the corresponding heterogeneous ortholog sets from these files. Following the extraction of orthologs for each species, we retrieved the gene coordinates by pruning the transcription start sites (TSSs) from the raw GTF file, which contained TSS information from exon 1. In a manner similar to the analysis of vertebrate promoters, we retrieved 1 kb upstream FASTA sequences and conducted FIMO analysis to identify the presence of 36 hypomethylated transcription factor binding sites (TFBSs) within these divergent sets of orthologs. The same stringent threshold was applied as in previous analyses, and TFBS occurrences per kilobase were calculated and visualized using consistent methodologies, leading to specific observations.

### GC content differential calculation

For all 105 species and their corresponding available orthologs, we merged the pruned FIMO output (with a motif occurrence threshold of p-value < 0.00001) with the associated FASTA sequences. This merged dataset included a column containing species names and was organized into taxon-specific files for non-amniotes, reptiles, aves, and mammals. We then calculated the GC content for each transcription factor binding site (TFBS) and its corresponding promoter sequence. To analyze trends in GC content across vertebrate taxa, we generated density plots for the promoter sequences. Following this, we calculated the GC content differential as the difference between TFBS GC content and promoter GC content. These calculations were based on the actual nucleotide sequences provided by the FIMO program. To visualize the GC content differential across the four vertebrate taxa, we created density plots using R. Additionally, we conducted a Wilcoxon signed-rank test to assess the significance of differences in GC content among the taxa.

### CpG-TpG double differential calculation

We subjected the pruned FIMO output to a detailed analysis of CpG and TpG counts within the reported transcription factor binding sites (TFBSs) and their associated 1 kb upstream promoter sequences in orthologs, as well as in all vertebrate promoters for each taxon. To assess shifts in CpG and TpG counts throughout vertebrate evolution, we plotted density maps for both dinucleotide counts. Subsequently, we calculated the CpG minus TpG differential separately for TFBSs and promoters, normalizing these values by sequence length. We then derived a second differential, calculated as TFBS CpG-TpG minus promoter CpG-TpG, which was plotted to illustrate differences across various taxa and ortholog-specific gene sets. The significance of these differences was evaluated using the Wilcoxon signed-rank test for the respective ortholog gene sets and all promoters.

### STREME analysis

The final pruned FIMO output comprised all taxon-specific files associated with promoters exhibiting GC content ranging from 30% to 70%, including data on orthologs. For each of the 36 hypomethylated transcription factor binding sites (TFBSs), the FIMO output strings were extracted based on the frequency of reported TFBS occurrences. These sequences were then converted into FASTA format for four taxa: non-amniotes, reptiles, aves, and mammals. Subsequently, STREME was run using default parameters, with the taxon-specific FASTA files serving as background for the analysis of 37ex-specific TFBSs. The output was meticulously evaluated, and motifs reported by STREME were selected based on their E-values for further taxon-wise analysis.

### motifStack analysis

The STREME output for the 36 hypomethylated motifs was meticulously examined, and position weight matrices (PWMs) for these motifs were compiled from each taxon into a single file for comparative analysis. These text files were then utilized as input for motifStack, enabling the alignment of the motifs. The aligned images produced were analyzed, leading to conclusions about the variations observed among the different taxa.

### Promoter sequence characterization of target genes

In this analysis, FIMO was utilized alongside GC content differential assessments and CpG-TpG double differential evaluations to investigate consistently repressed genes affected by various forms of CGGBP1 at 37°C and 40°C. The objective was to define the characteristics of these gene sets and uncover taxon-specific alterations related to the different CGGBP1 forms under varying temperature conditions. The gene sets were categorized into three distinct groups - 37ex, 40ex, and 37∩40 through k-means clustering. The GC content of their individual promoters was computed, revealing no significant differences in total promoter GC content as confirmed by a Wilcoxon signed-rank test. Following this, FIMO analysis was conducted to explore the occurrence of transcription factor binding sites (TFBS) per kilobase within these promoters. To discern the impact of GC content on TFBS, analyses focused on high GC content (>50) and low GC content (<50) TFBS individually. A paired t-test was employed to compare the three gene sets (37ex vs.40ex, 37ex vs.37∩40, and 40ex vs.37∩40). Subsequent comparisons of TFBS GC content among the three gene sets were carried out using the Wilcoxon signed-rank test to highlight those TFBS with significantly higher GC content (p-value < 0.05). Commonalities among significant TFBS were illustrated using a Circos plot. Furthermore, the quantitative differences in GC content for these TFBS compared to JASPAR representative TFBS were assessed, leading to the creation of a mirror density plot that emphasized median values.

#### Softwares

The following software were used: fastp (version 0.23.4), Bowtie2 (version 2.5.1), deepTools (version 3.3.2), bedtools (version 2.27.1), MACS2 (version 2.2.7.1) and IGV. Motif discovery was performed with MEME Suite (version 5.5.1), using FIMO and STREME. motifStack (version 1.48.0) was used for visualization and comparison of identified motifs, ClustVis web tool (30) was used for generating heatmaps of motif occurrences. GraphPad Prism and R were used for generating other plots and conducting statistical analyses.

### Commands

The commands used for the data handling and analyses can be made available on request.

## Abbreviations

Ac: *Anolis carolinensis*
DBD: DNA Binding Domain
Ev: Empty vector
Gg: *Gallus gallus*
Hs: *Homo sapiens*
Lc: *Latimeria chalumnae*
MeDIP-seq: Methylated DNA Immuno-precipitation Sequencing
NLS: Nuclear localisation signal
TFBSs: Transcription Factor Binding Sites
37ex: Genes exclusively repressed by Hs CGGBP1 at 37°C
40ex: Genes exclusively repressed by Hs and Gg CGGBP1 at 40°C
37∩40: Gene repressed by Hs CGGBP1 at 37°C as well as 40°C and repressed by Gg CGGBP1 at 40°C

