## Supplementary figure for "CGGBP1 from higher amniotes restricts cytosine methylation and drives a GC-bias in transcription factor binding sites at repressed promoters"

Fig. S1

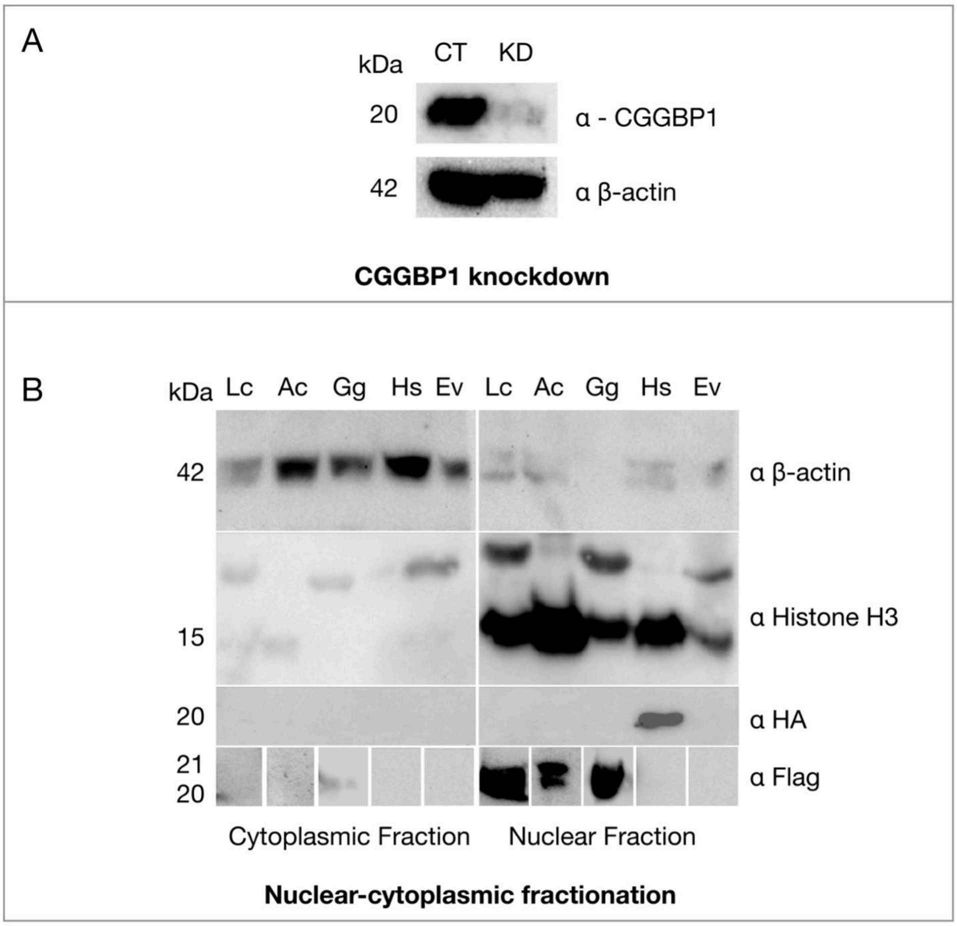


Fig. S1. CGGBP1 knockdown and sub-cellular localization of its various forms in HEK293T cells. A: shRNA-mediated knockdown of human CGGBP1 in HEK293T cells targeting four distinct regions of the ORF, as detailed in the Methods. HEK293T control (CT) cells express endogenous CGGBP1 at 20 kDa, while HEK293T knockdown (KD) cells show a reduction of CGGBP1 levels to approximately 70-80%. Beta-actin (42 kDa) was used as a loading control. B: Nuclear-cytoplasmic fractionation assays reveal that all vertebrate forms of CGGBP1 are predominantly localized to the nucleus after being synthesized in the cytoplasm, with little to no CGGBP1 detected in the cytoplasm. β-actin was used as a cytoplasmic marker, while total Histone H3 served as a marker for the nuclear fraction. Lc, Ac, and Gg CGGBP1 were detected using a FLAG-tag antibody, and Hs CGGBP1 was probed with an anti-HA antibody. The lanes were rearranged to ensure proper alignment for β-actin. To account for the differing amounts of sample, the nuclear fraction was loaded at half the volume (7μl) compared to the cytoplasmic fraction (15μl), which revealed variations in expression levels between the samples.

Fig. S2

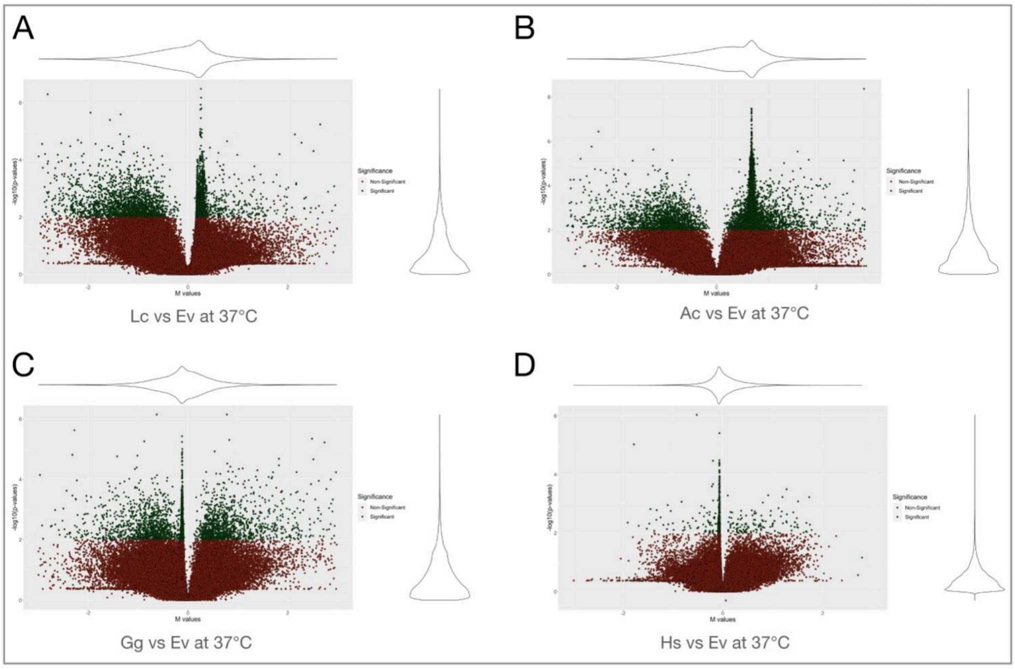


Fig. S2: Global gene expression changes caused by expression of the indicated forms of CGGBP1 (Lc, Ac, Gg or Hs) when compared against the pattern expressing an empty vector (Ev) at 37°C. In the samples representative of non-amniotes (Lc) and lower amniotes (reptiles) widespread gene deregulation were observed with spikes of derepressed genes (visible as positive M-value spikes and modes of the density distribution at the top. In the representatives of higher amniotes (Gg and Hs) however the larger change in expression was repression. The extent of gene deregulation decreased with similarity between the forms of CGGBP1 representatives and the taxa they represent. Consistently differentially expressed genes (p value <0.01 in three replicate experiments) shown as green dots. The Y-axes show -log10 p values and the X-axes show M values were calculated using the geometric means of the three replicate experiments.

Fig. S3

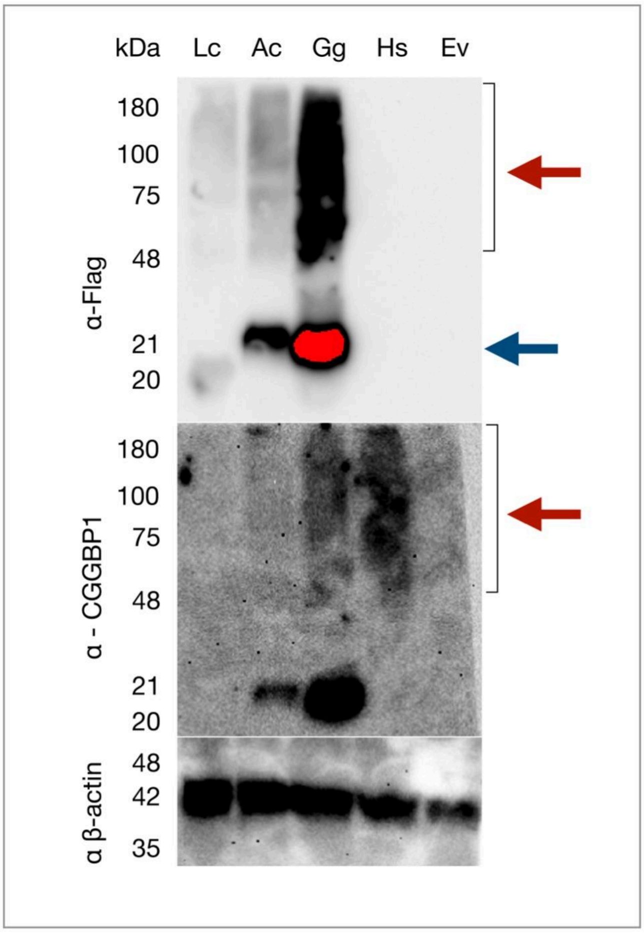


Fig. S3: CGGBP1 overexpression under heat stress conditions. HEK293T cells were transfected with various CGGBP1 constructs and exposed to heat stress at 40°C for 48 hours (details in methods). Western blot analysis revealed multiple bands corresponding to different molecular weights of CGGBP1, aside from the typical 20 kDa species, as marked by red arrows, indicating potential heat-induced modifications such as ubiquitination. Non-human (non-Hs) CGGBP1 isoforms were detected using anti-FLAG antibodies. In contrast, human CGGBP1 (Hs CGGBP1) could not be detected with the HA-tag antibody, as reported previously. However, using a polyclonal CGGBP1 antibody, we identified Hs CGGBP1 running at a higher molecular weight than expected, while the conventional 20 kDa form was absent, despite using multiple antibodies. The polyclonal antibody also detected non-Hs CGGBP1 due to sequence similarity. Beta-actin was used as a loading control. CGGBP1 forms exhibit differential expression levels, with the non-amniotic form Lc being undetectable with the FLAG-tag unless the expression of the other samples reaches saturation, as indicated by the blue arrow.

#### Fig. S4

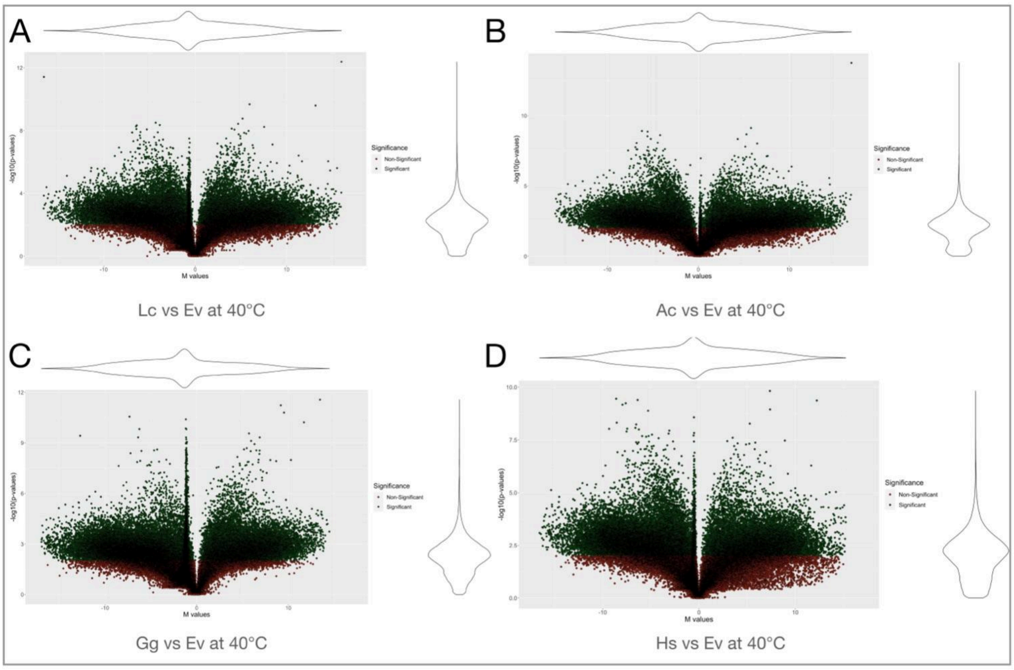


Fig. S4: Global gene expression changes brought by the indicated forms of CGGBPs at 40°C. A-D: The global deregulation of gene expression was widespread at 40°C with no clear directions of deregulation and differences between species forms of CGGBP1 as observed at 37°C (Fig S1). Consistently differentially expressed genes (p value <0.01 in three replicate experiments) shown as green dots. The Y-axes show -log10 p values and the X-axes show M values were calculated using the geometric means of the three replicate experiments.

Fig. S5

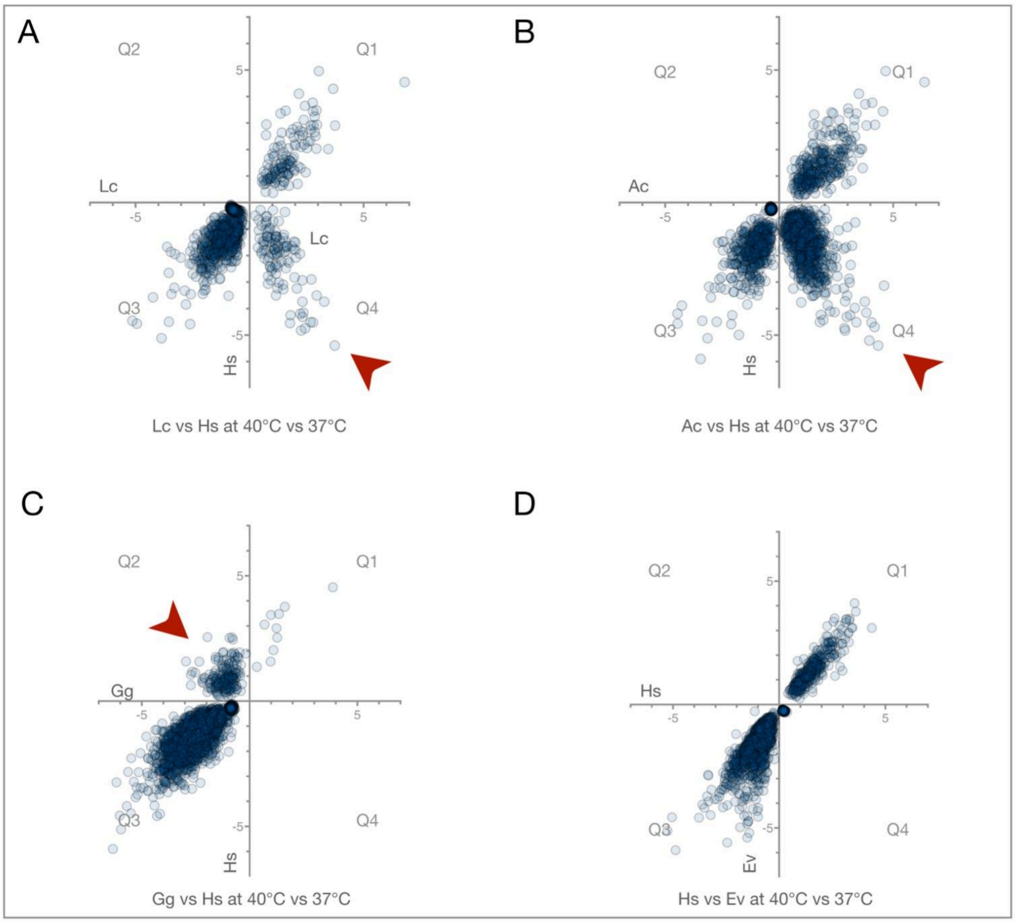


Fig. S5: Comparisons of gene expression changes caused by heat stress in the presence of various different forms of CGGBP1 reveal gene repression under heat stress as a feature of CGGBP1 from higher amniotes (Gg and Hs) not present in lower amniotes (Ac) or non-amniotes (Lc). Only consistently deregulated genes (p-value < 0.01 in 37°C versus 40°C comparisons) are plotted on the Y-axes and from Ev on the X-axes. Quadrants: Q1- significant induction in Ev as well as the indicated sample, Q2- significant repression in the indicated sample but induction in Ev, Q3- significant induction in Ev as well as the indicated sample, and Q4- significant repression in Ev but induction in the indicated sample. Red arrowheads mark the regions showing data points where significantly differentially expressed genes upon heat stress have opposing directions of change as compared to Ev. Lc and Ac show abnormal induction and Gg shows abnormal repression.

Fig. S6

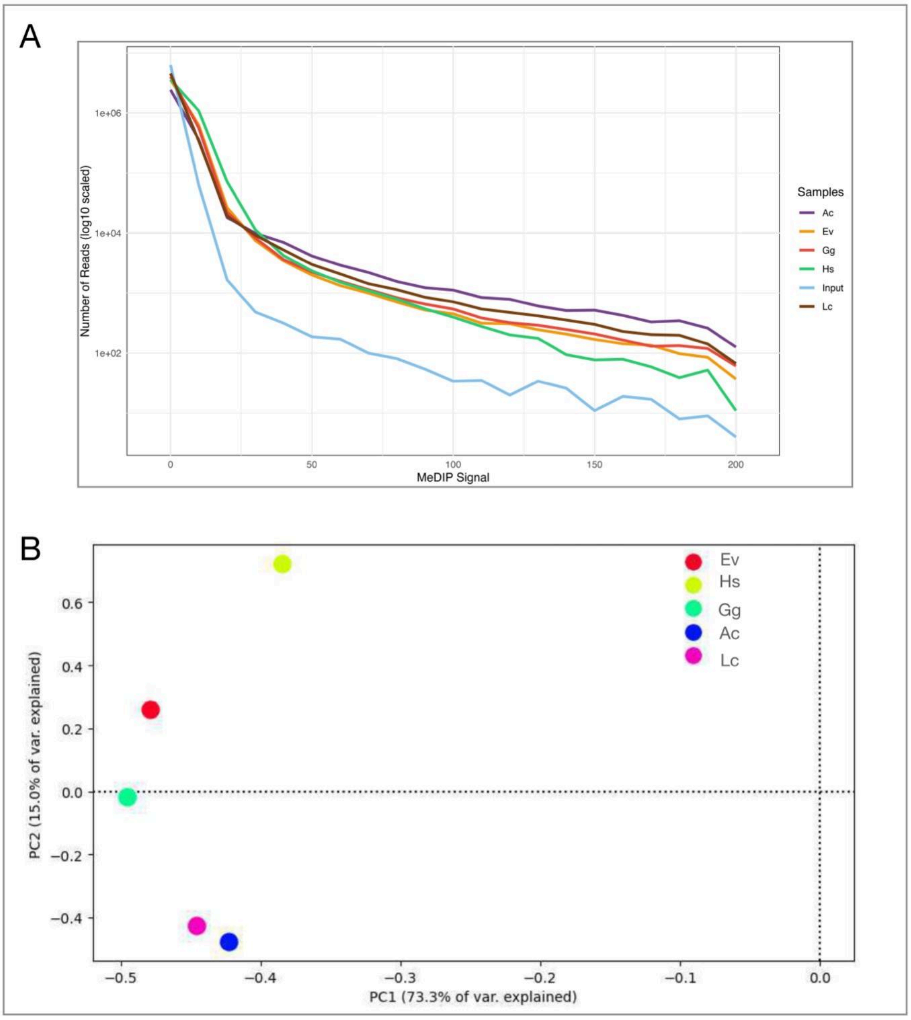


Fig. S6: An overall summary of MeDIP-seq. A: Frequency distribution of MeDIP signals in 0.2kb bins genome wide. Various samples are represented by different colors (Lc - brown, Ac - purple, Gg - red, Hs - green, Ev - orange and Input - Blue). All samples show enrichment over input across a wide range of signals. Hs mitigates cytosine methylation and shows a frequency distribution shift towards lower methylation signals whereas Gg follows the distribution pattern of Ev. Overall Hs and Gg restrain methylation genome-wide. In contrast, Ac and Lc expression increase methylation and the distribution shifts towards higher signal bins. B: PCA of MeDIP signal genome wide shows that Ev, Hs and Gg are more similar and cause similar patterns of methylation while Lc and Ac cluster together due to the similarity in methylation pattern caused by their expression.

Fig. S7

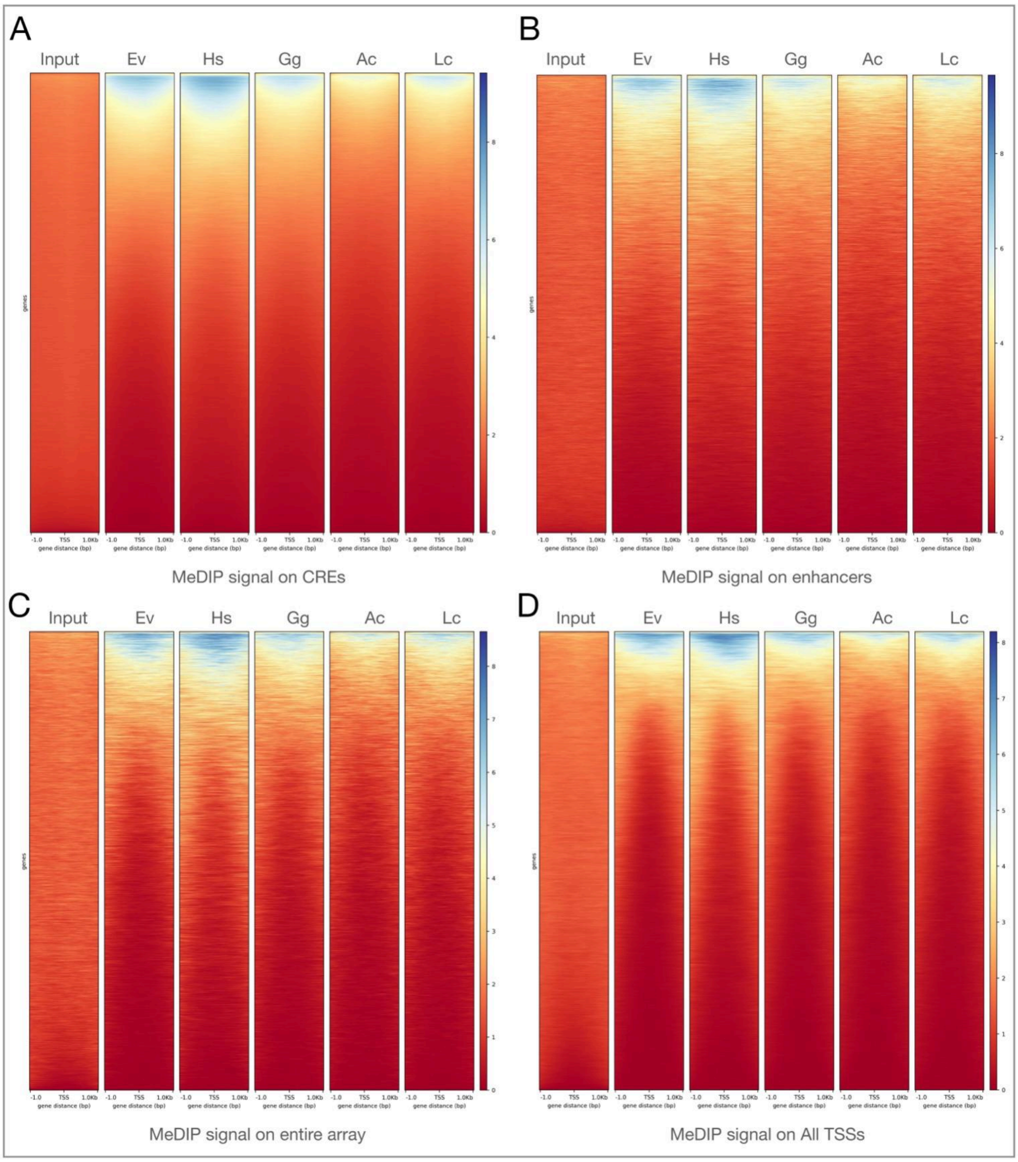


Fig. S7: Overview of MeDIP signals on regulatory elements and promoters. A-D: Input shows no enrichment of methylation and all samples show enrichment of methylation as compared to input. MeDIP signals on conserved regulatory elements (CREs) (A), enhancers (B), all TSS (robust and permissive) (C) and the promoters (D) of probe set on the microarrays show a similar pattern of methylation. The genomic regions are sorted in descending order of MeDIP signals.

Fig. S8

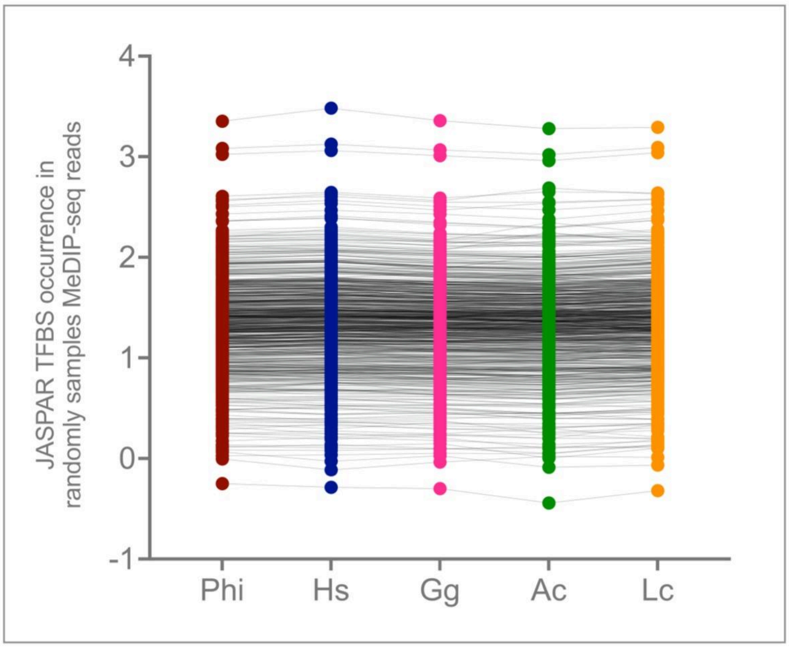


Fig. S8: MeDIP reads show no major differences in presence of JASPAR (vertebrate) TFBSs. The Y axis represents TFBS occurrences (log scale) fetched through FIMO search on randomly sampled 1 million MeDIP seq reads from the indicated samples. Each TFBS is paired between different samples by horizontal gray lines.

Fig. S9

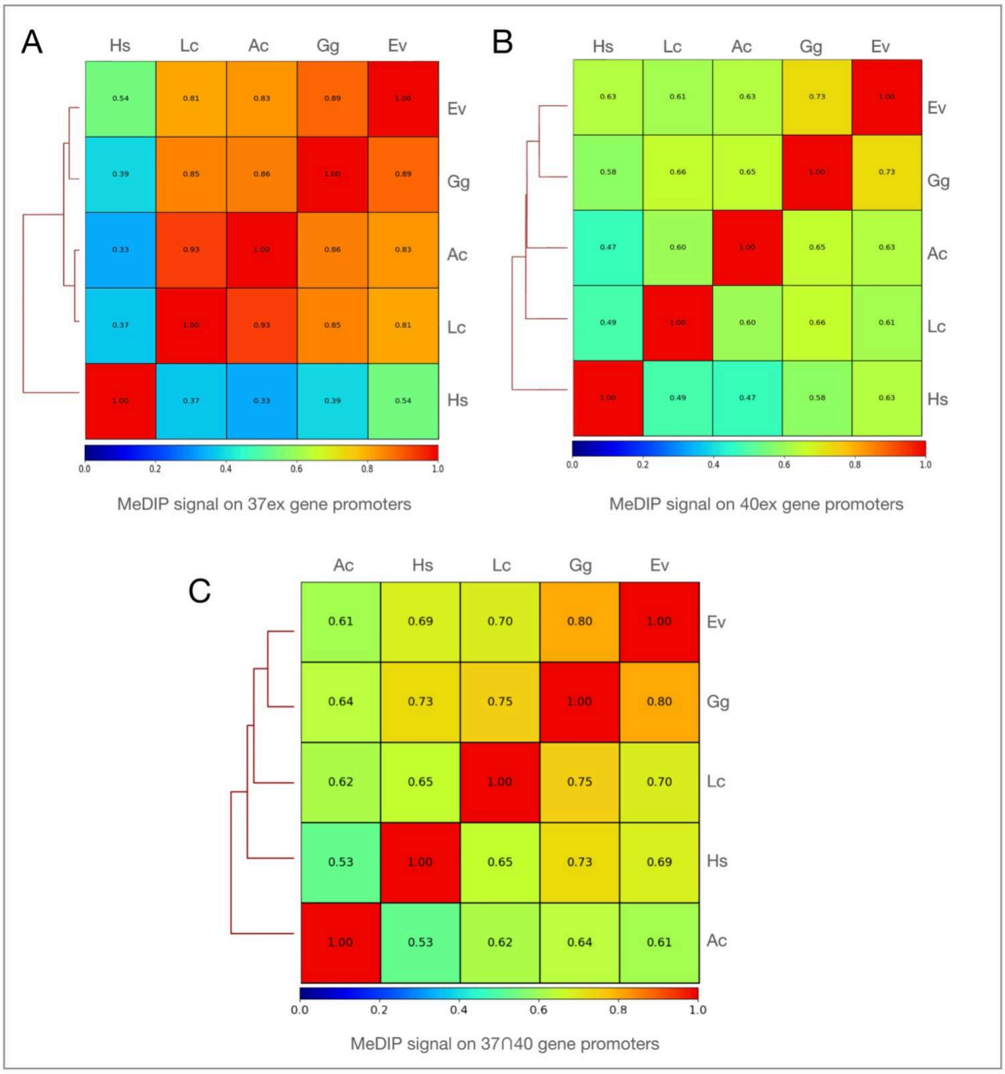


Fig. S9: MeDIP signal on of 37ex, 40ex and 37∩40 gene promoters. A: Methylation in 1 kb upstream region of 37ex promoters recapitulates gene expression pattern at 37°C. The methylation pattern on these genes sets Hs apart from other samples and clusters Gg and Ev into a separate cluster while Lc and Ac cause a similar pattern of methylation. B and C: Such a strong distinction of methylation pattern is not manifested at 40ex as well as 37∩40 gene promoters. These methylation patterns closely resemble the gene repression patterns caused by Lc, Ac, Gg and Hs (Figure 2D-E).

Fig. S10

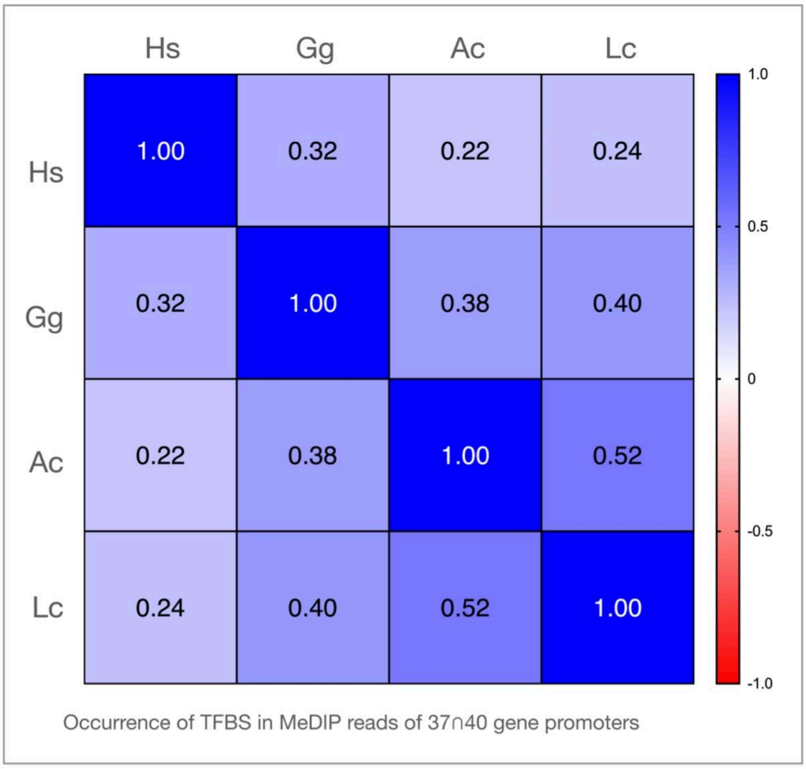


Fig. S10: Correlations between methylation patterns at TFBSs in 1kb promoters of genes repressed by CGGBP1 at 37∩40 gene promoters shows no specific pattern which sets apart TFBSs methylation pattern induced by changing various forms of CGGBP1. It appears that these specific genes which flip for Gg when the heat-stress in play are moderately dependent on TFBS methylation of some if not all TFBSs.

Fig. S11

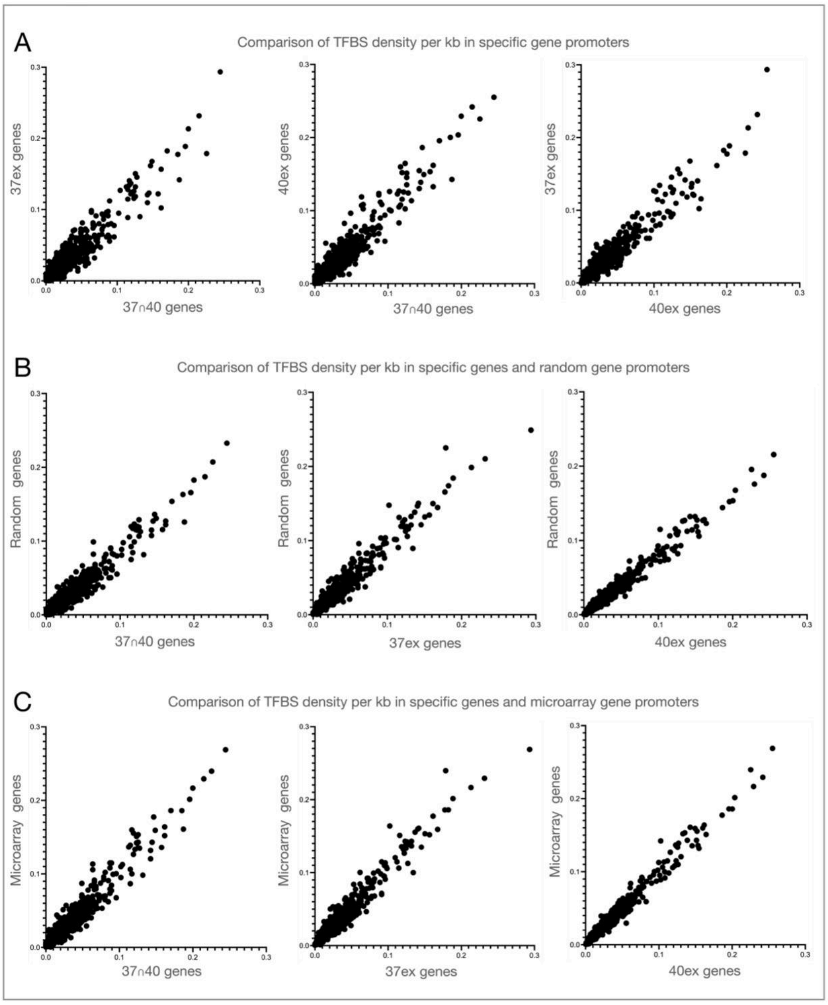


Fig. S11: Analysis of TFBS occurrence and density in gene promoters. A: The analysis of TFBS within the 1kb region upstream of the 37ex, 40ex, and 37∩40 gene promoters shows no significant variation in their frequency. B: The density of TFBS per kilobase in these specific gene sets is not different from that of a randomly chosen set of genes on the microarray. C: The TFBS occurrence in the 37ex, 40ex, and 37∩40 gene promoters closely matches the overall TFBS density observed in all genes on the microarray.

### Fig. S12

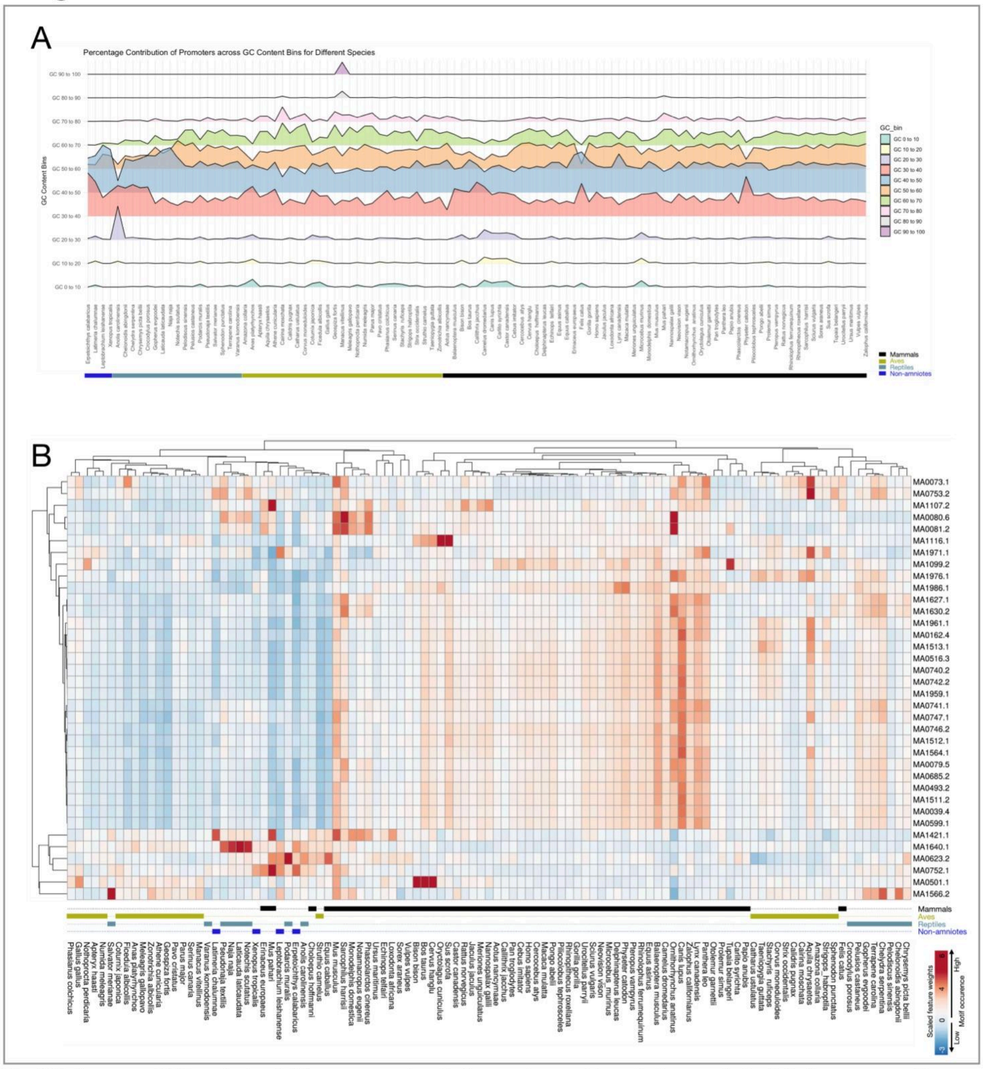


Fig. S12: A survey of GC contents of all known promoters in the species used for orthology analysis. A: In the genome-wide promoter sets known for four non-amniotes, 17 reptiles, 27 aves and 57 mammals shows that most promoters are in the GC content range of 30% to 70%. The analysis of effects of different forms of CGGBP1 on TFBS methylation in repressed gene promoters and their orthologs were restricted to this set of promoters in the 30% to 70% range for eliminating the outlier effects. B: The occurrence of the 36 hypomethylated TFBSs in the promoters genome-wide is higher in mammals and aves and clusters majority of them together. The lower amniotes (reptiles) and non-amniotes have lower abundance of these TFBSs with high variability. These hypomethylated TFBSs are also majorly GC-rich and the larger cluster of the rows classifies the high GC-content TFBSs together. It is the high GC-content hypomethylated TFBSs which cluster higher amniotes differently from the non-amniotes and reptiles.

### Fig. S13

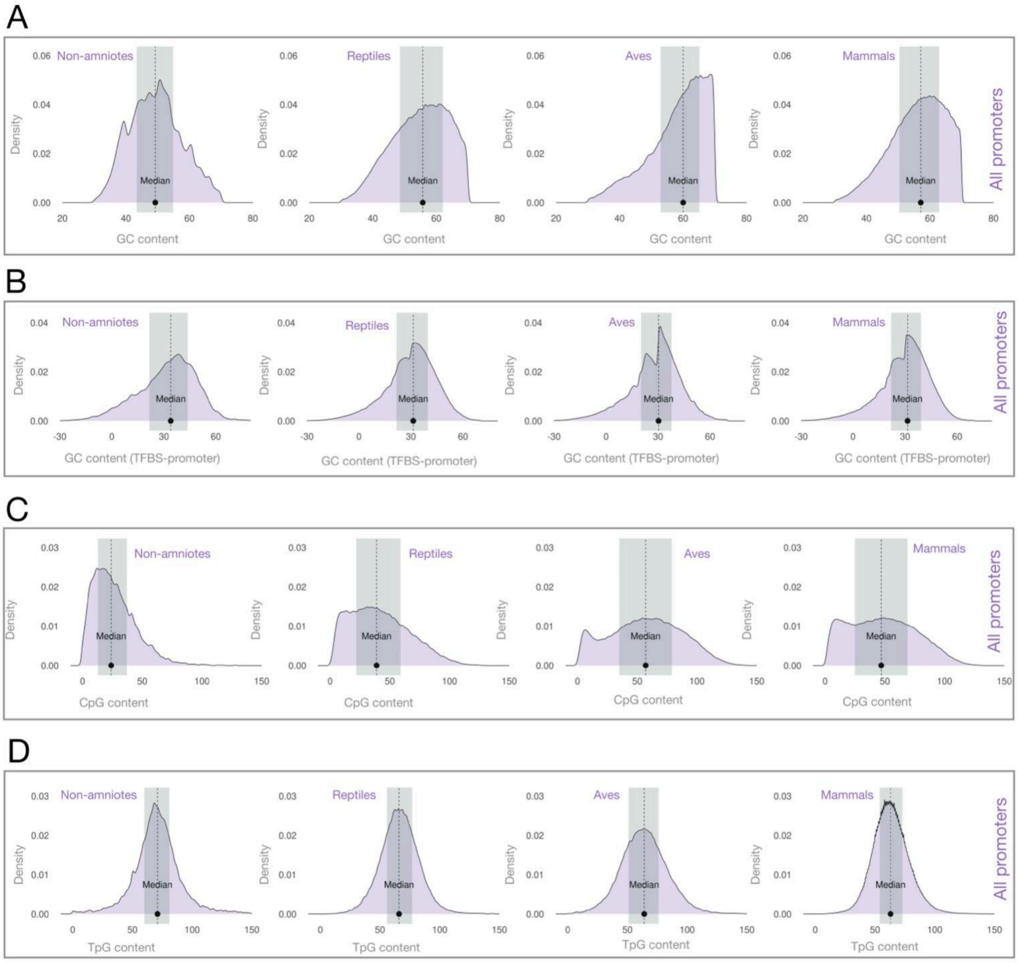


Fig. S13: Patterns of GC, CpG and TpG contents in the promoters and TFBSs of all genes from 105 species used in the analyses. A: Frequency distributions of the promoter GC contents show that the GC content of promoters have increased in amniotes (top panel) with strong increases in higher amniotes. B: The GC content increase in amniotes is accompanied by a clear increase in hypomethylated TFBS GC contents over and above the promoter GC contents (calculated as TFBS GC content-promoter GC contents). The TFBS-Promoter GC content differentials are bimodal in amniotes with a conspicuous increase in the mode with higher differential value (X-axis). C: Similar to the increase in TFBS GC content, the CpG content has also increased in amniotes with strong CpG enrichment observed in aves and mammals. D: The CpG content increase in TFBSs in higher amniotes (C) is not accompanied by any discernible directional changes in TpG contents. These statistics show that the hypomethylated TFBSs have evolved a higher GC content in higher amniotes, of which a small part could be contributed to by lower rates of CpG to TpG transitions, a possibility we have tested further on.

#### Fig. S14

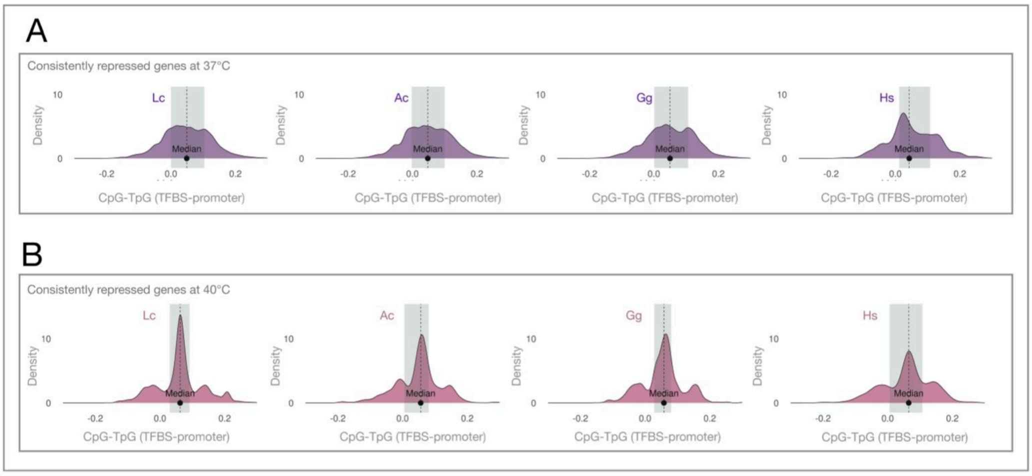


Fig. S14: The significantly differentially repressed genes by Lc, Ac, Gg and Hs in replicate experiments show CpG-TpG differential profiles very different from the corresponding GC differential profiles (Fig 5A and B). A: The CpG-TpG differentials of hypomethylated TFBSs compared to the promoters in genes repressed at 37°C (n=3 replicates, p<0.01). B: The CpG-TpG differentials of hypomethylated TFBSs compared to the promoters in genes repressed at 37°C (n=3 replicates, p<0.01). Noticeably, only Hs, and to some extent Gg, showed a CpG-TpG differential pattern in 37°C repressed promoters which resembled that observed in 40°C repressed promoters. Unlike GC content in the TFBSs, these results indicated that the CpG content of the TFBSs is relevant to such a robust repression by Hs CGGBP1 that it is independent of the heat stress.

### Fig. S15

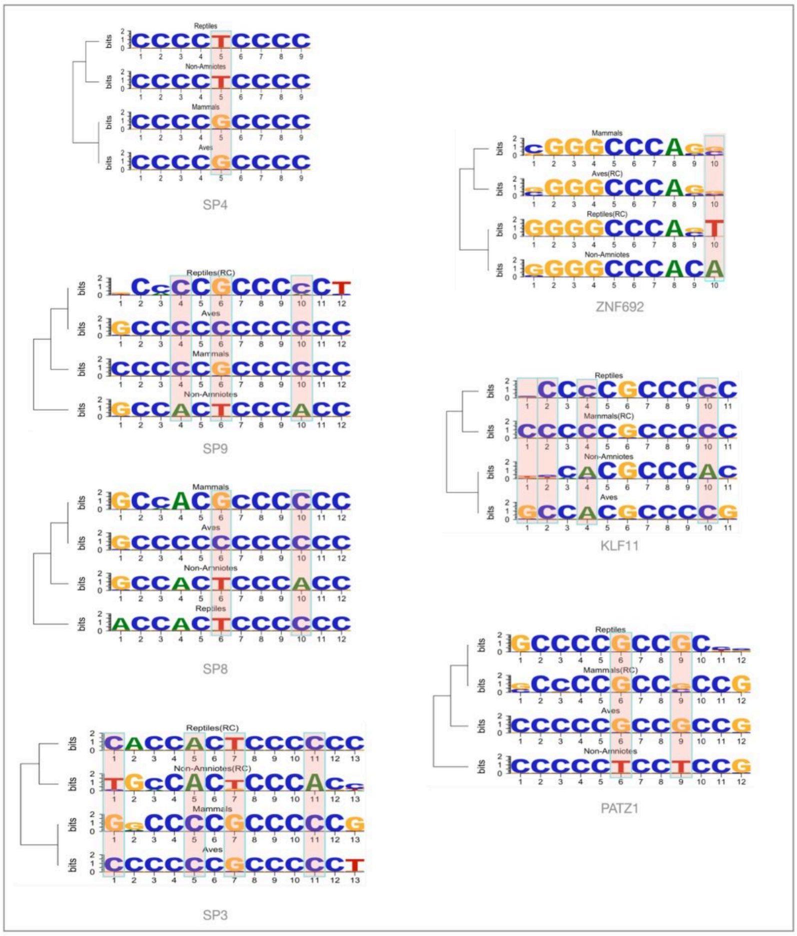


Fig. S15: Examples of GC retention in TFBSs in CGGBP1-repressed promoters and their orthologs which can not be explained by restriction of cytosine methylation. These findings suggest that cytosine methylation restriction is not the only possible mechanism contributing to GC retention in TFBSs under the influence of CGGBP1 of higher amniotes.

### Fig. S16

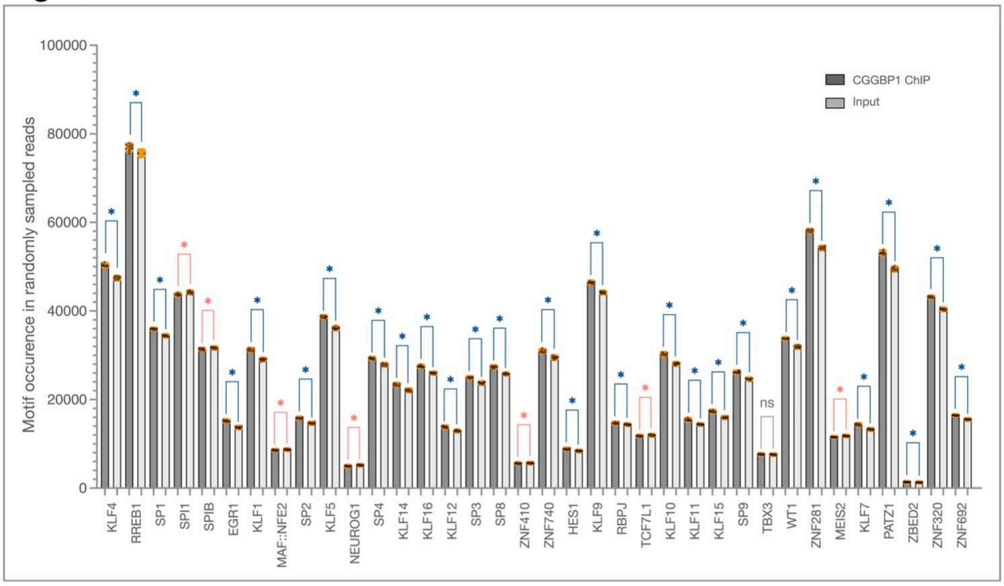


Fig. S16: The 36 hypomethylated TFBSs with methylation restriction and associated GC retention provided by CGGBP1 (28) were significantly enriched ($p < 0.01$, blue asterisks) in the publicly available CGGBP1 ChIP-seq dataset (PMID: 39605320). Seven TFBSs showed significant depletion (red asterisks), and one TFBS showed no significant change (grey 'ns'). Comparisons were made against the input, using 10 independent read samplings (n = 1 million).
