## Additional file 1 for "CGGBP1 from higher amniotes restricts cytosine methylation and drives a GC-bias in transcription factor binding sites at repressed promoters"

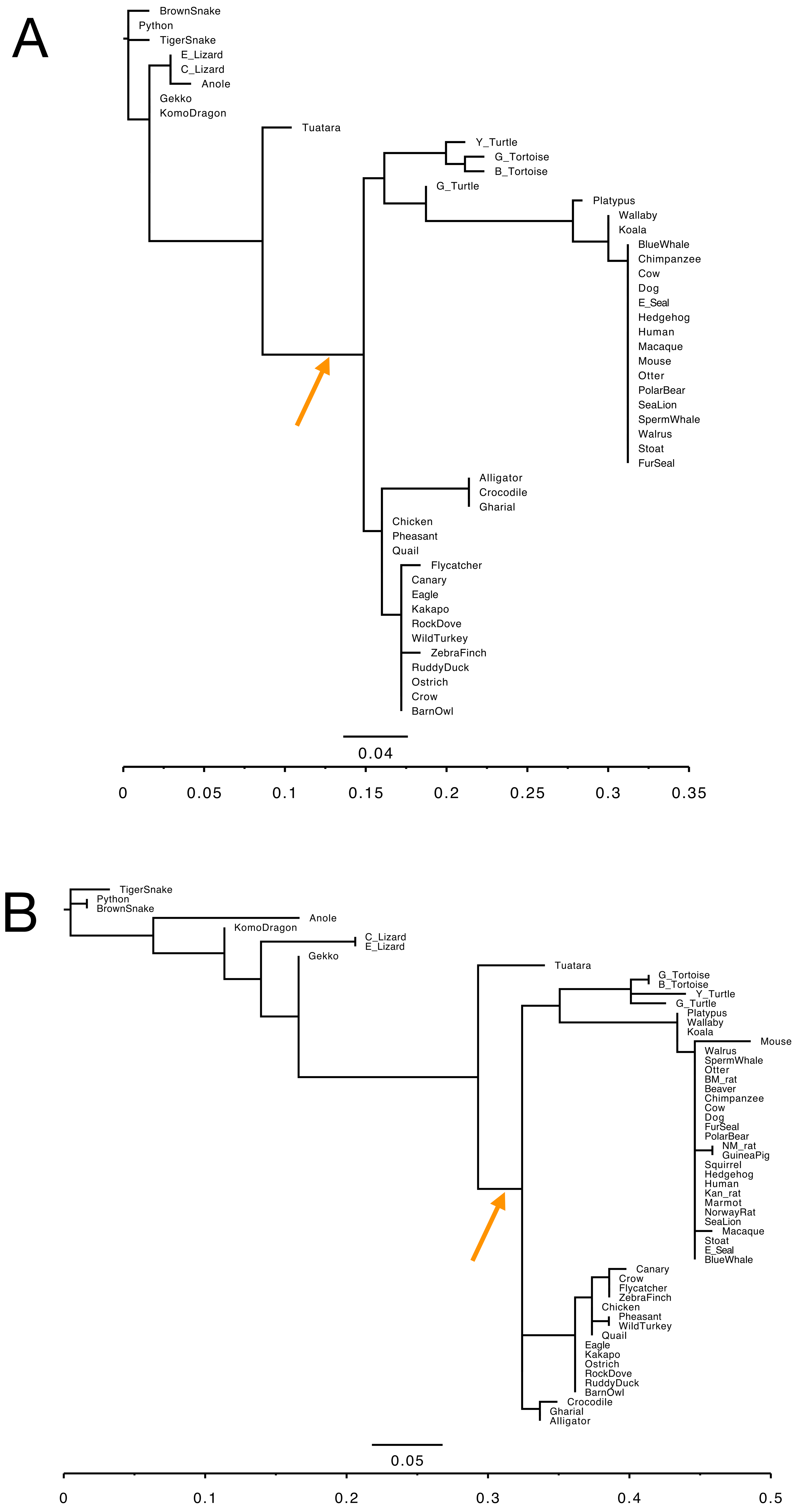


Additional file 1, fig 1. Phylogenetic trees derived from multiple sequence alignments of N-terminal (A) and C-terminal (B) parts of 49 amniotic CGGBP1s. The N- or C-terminal parts were identified as upto amino acid locations 79-RKKQR-84 (N-terminal part, A) or downstream sequence (B) of the human CGGBP1 sequence. For all other species the corresponding sites were selected through a pair-wise alignment against the human sequence. The common ancestral node of archosauromorpha and mammals is highlighted by an orange arrow. The testudines map variably as an outgrip in the lineage giving rise to the mammals. It is noteworthy that the N-terminal part is almost invariant in the eutherians, unlike the C-terminal part. A comparison of the divergences happen on the X-axes show that the overall changes in C-terminal part are higher than that  in the N-terminal part.

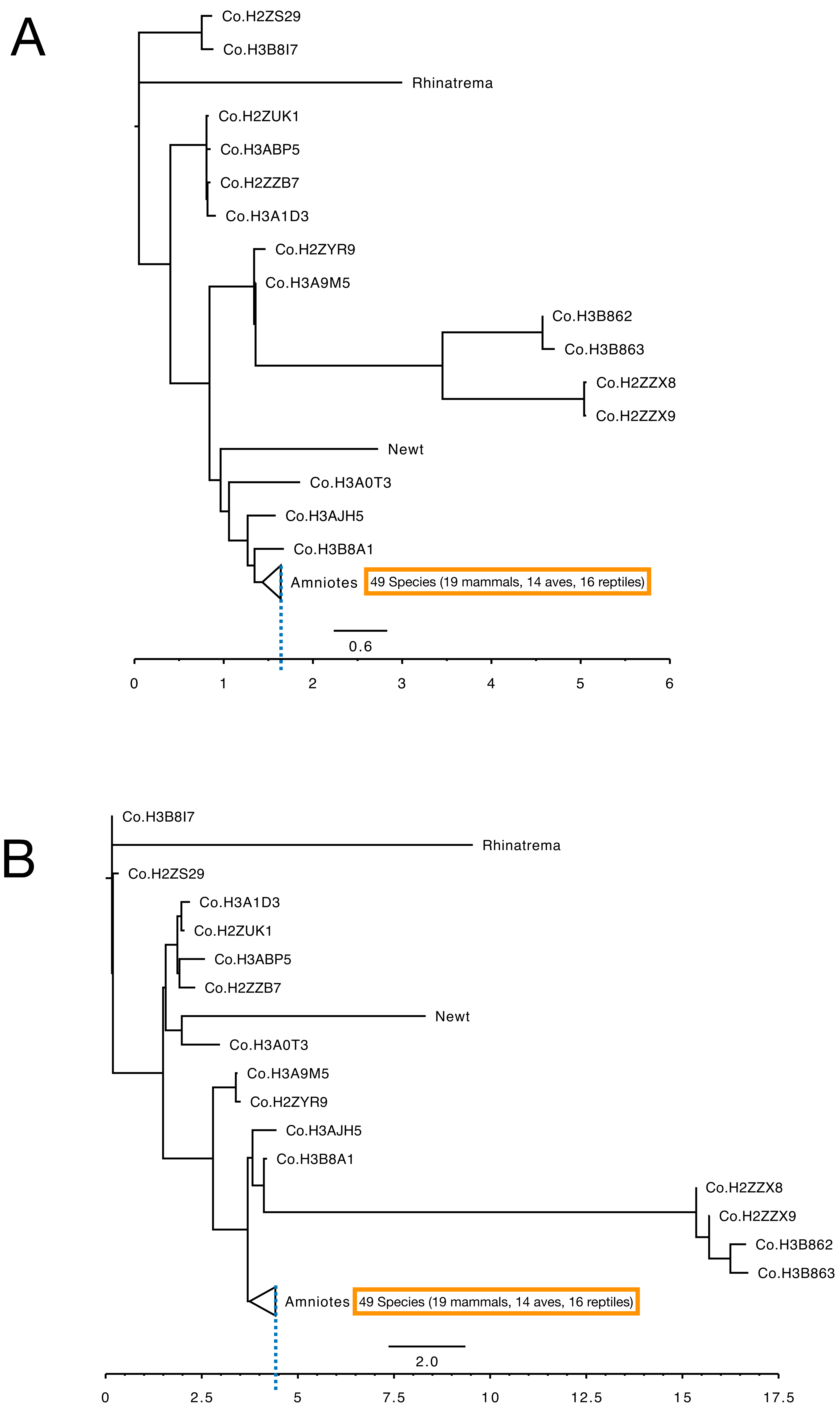


Additional file 1, fig 2: Phylogenetic trees based protein amino acid sequence similarities for the N-terminal (A) or the C-terminal parts of CGGBPs from (Latimeria (suffixed Co) and amphibians (Rhinatrema and Newt) as non-amniotic forms and 49 amniotic forms. The protein sequences were divided into N- or C-terminal parts at a location corresponding to the amino acid locations 79-RKKQR-84 of the human CGGBP1 sequence by a paired alignment. The X-axis indicates the relative divergence which is higher in the C-terminus (B) as compared to the N-terminus (A) in the non-amniotes. The highest divergence in all amniotes maps to (indicated by a blue dashed line) 1.5 to 2 units for N-terminus as 4.5 to 5 units for C-terminus.
