## Additional file 2 for "CGGBP1 from higher amniotes restricts cytosine methylation and drives a GC-bias in transcription factor binding sites at repressed promoters"

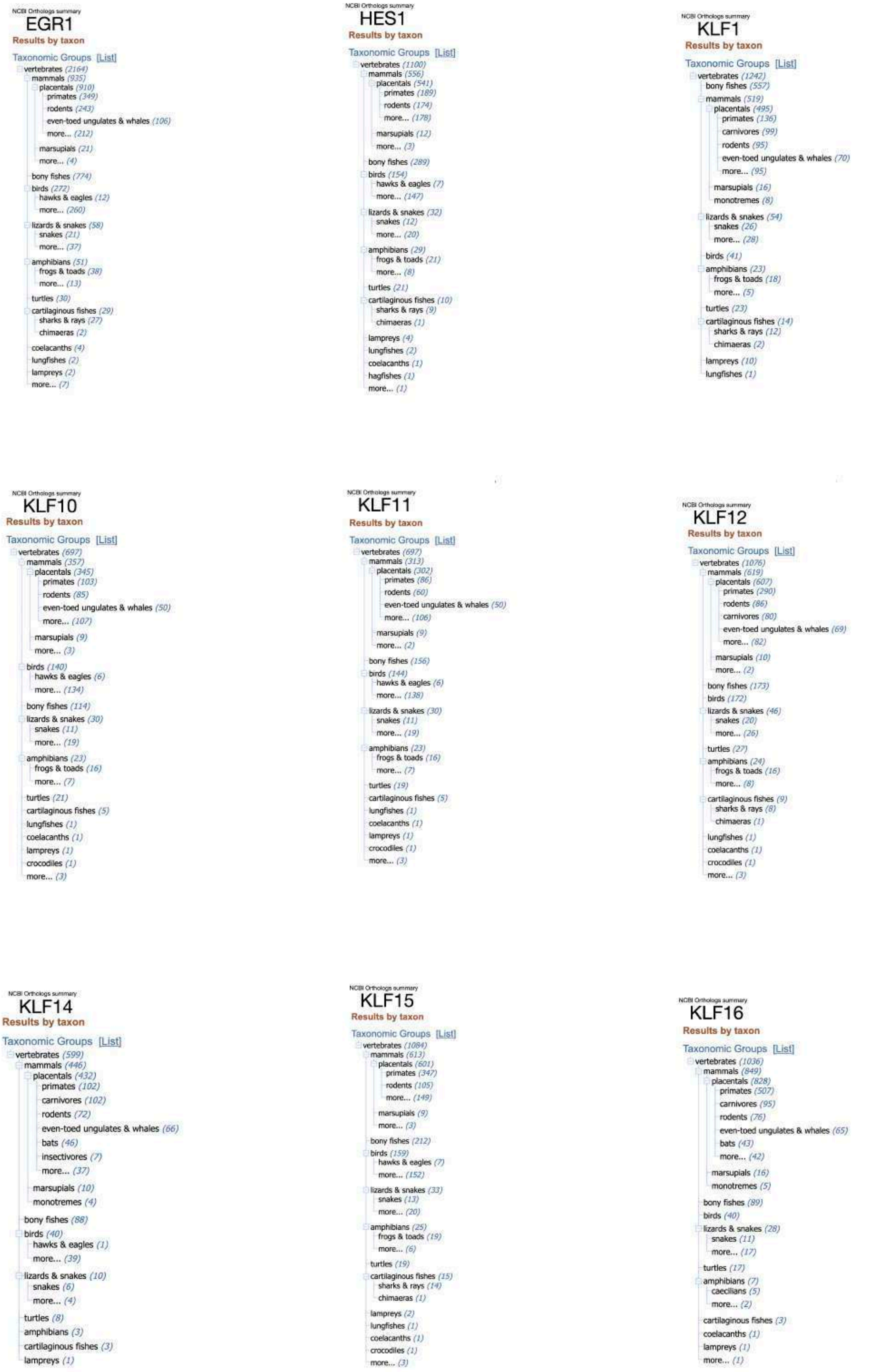


Additional file 2, page 1: A collage of NCBI Orthologs output for the 36 transcription factor genes corresponding to the factors binding to the hypomethylated motifs. Most of these factors have orthologs throughout the vertebrates.

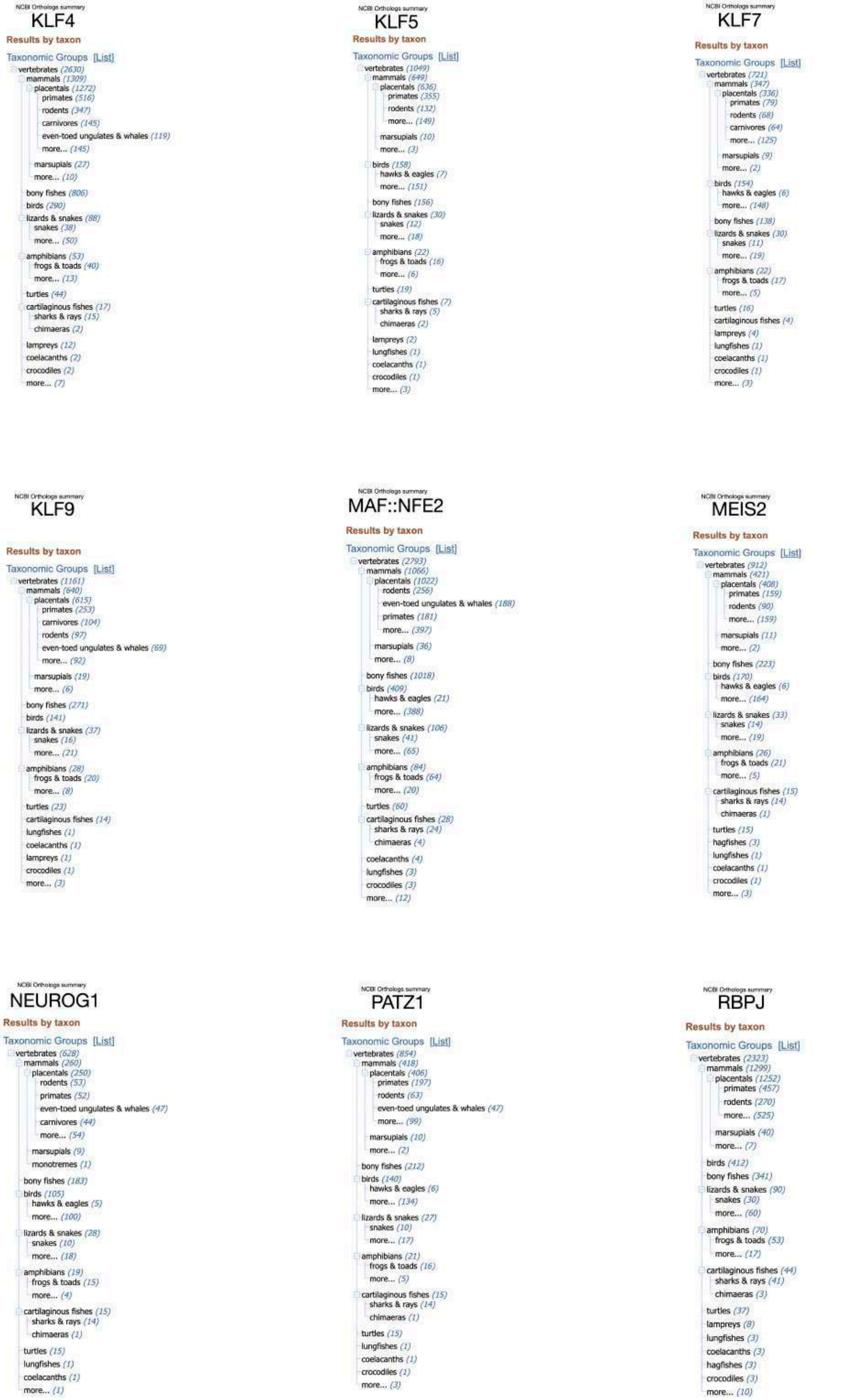


Additional file 2, page 2: A collage of NCBI Orthologs output for the 36 transcription factor genes corresponding to the factors binding to the hypomethylated motifs. Most of these factors have orthologs throughout the vertebrates.

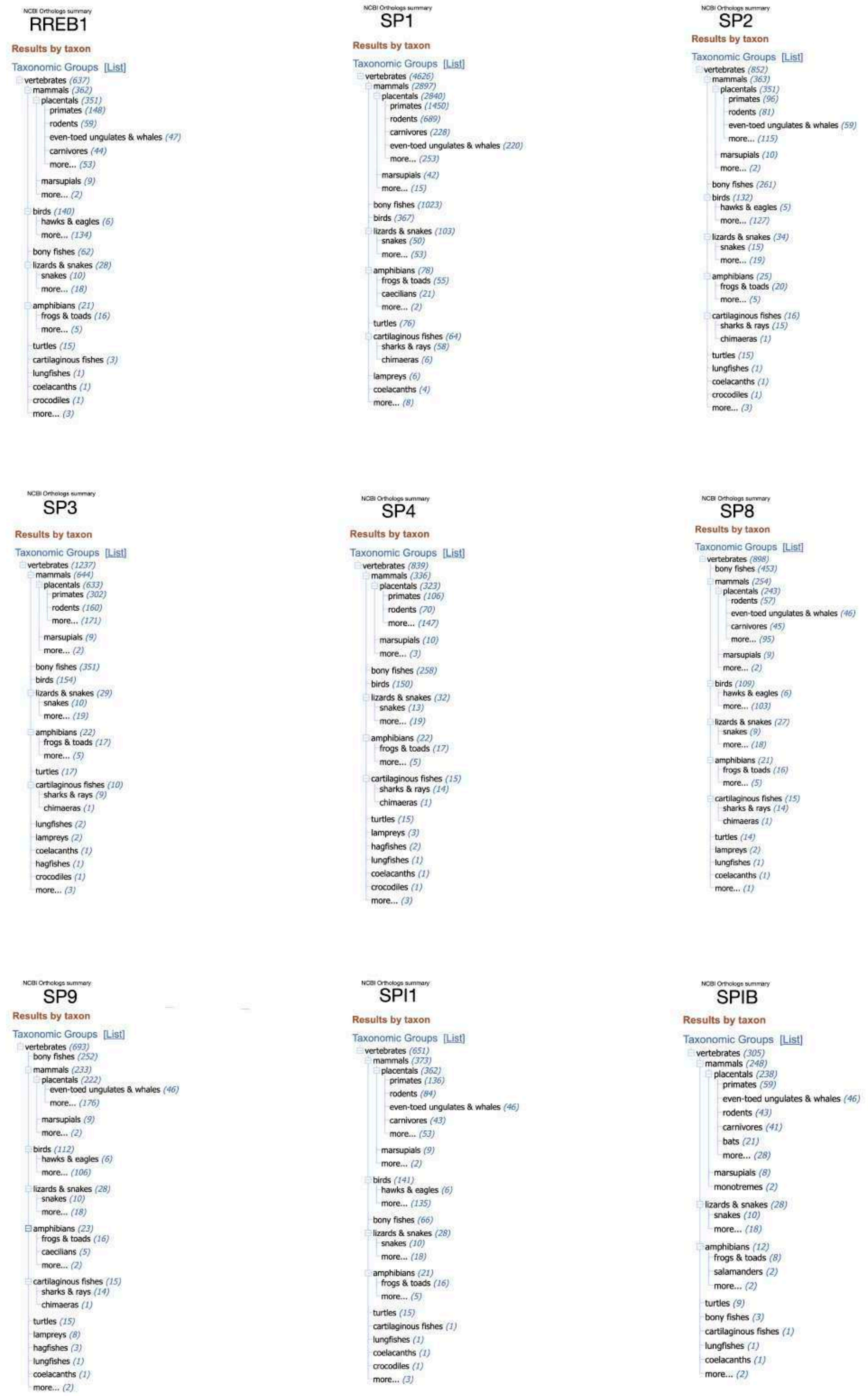


Additional file 2, page 3: A collage of NCBI Orthologs output for the 36 transcription factor genes corresponding to the factors binding to the hypomethylated motifs. Most of these factors have orthologs throughout the vertebrates.

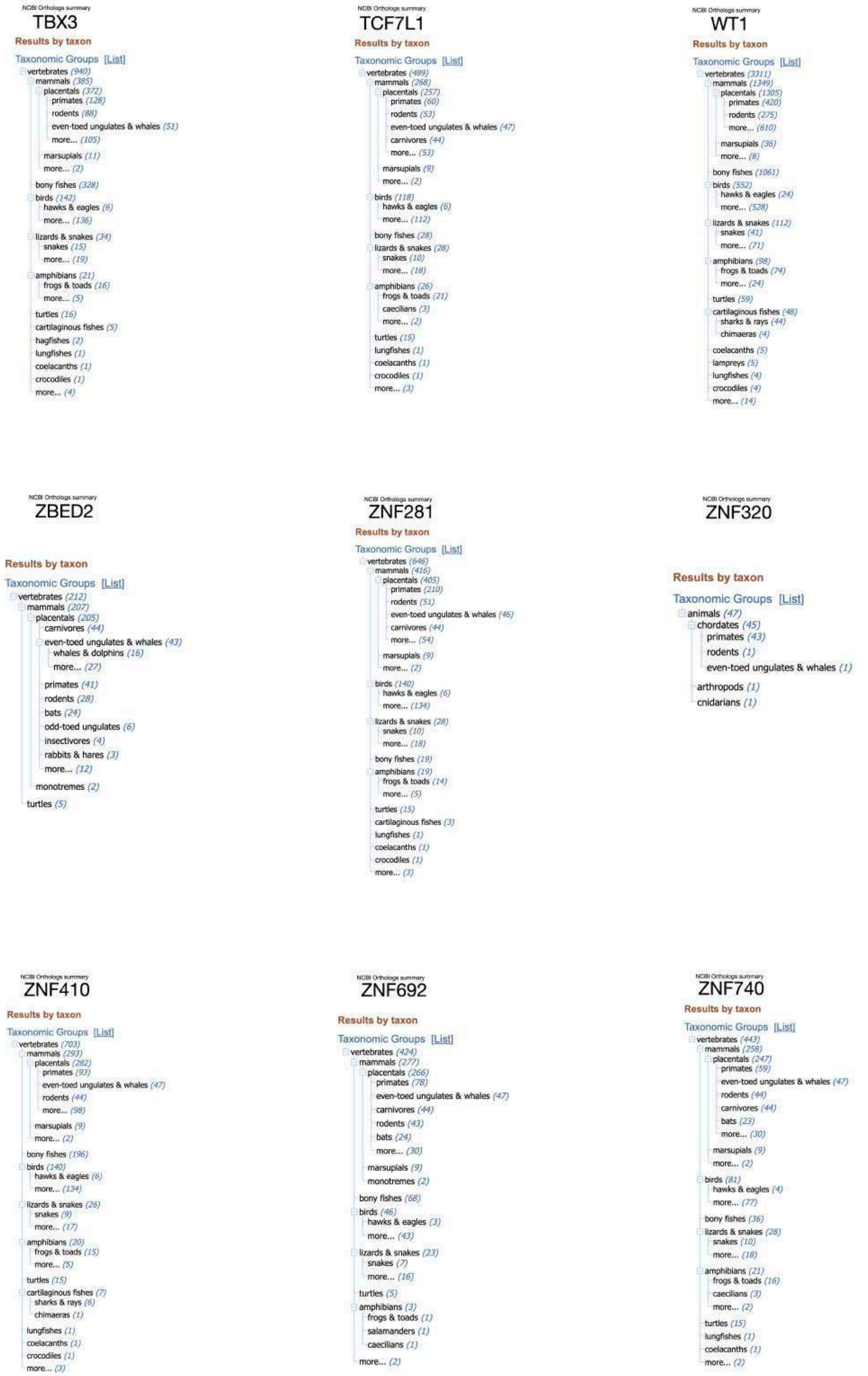


Additional file 2, page 4: A collage of NCBI Orthologs output for the 36 transcription factor genes corresponding to the factors binding to the hypomethylated motifs. Most of these factors have orthologs throughout the vertebrates.
