## Additional file 3 for "CGGBP1 from higher amniotes restricts cytosine methylation and drives a GC-bias in transcription factor binding sites at repressed promoters"

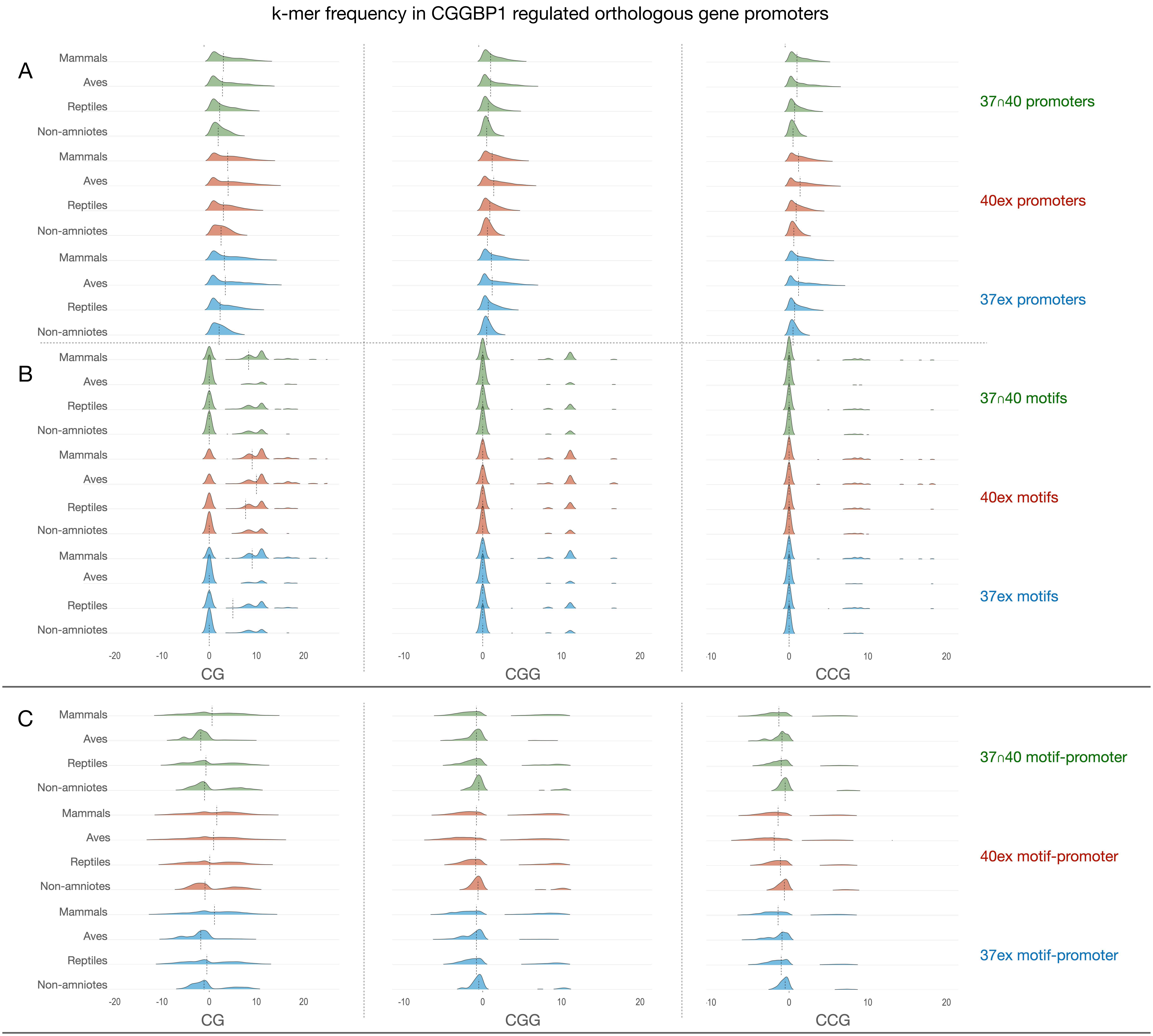


Additional file 3, fig 1: Frequency distributions of abundance of different k-mers (CG, CGG and CCG) calculated in promoters (A) or hypomethylated motifs (B) show a high value bias incrementally as the taxon groups transition from non-amniotes to mammals. The bias is clearly visible in promoters (A) and to a lesser extent in motifs (B). The effects of these compositional biases are eliminated by calculating the k-mer occurrences are motif-minus-promoter (motif-promoter) differentials (C). The description of motif GC content and CpT-TpG transition rates (main figures 3, 4 and 5) have all been calculated by using the motif-promoter methodology.
