## Additional file 4 for "CGGBP1 from higher amniotes restricts cytosine methylation and drives a GC-bias in transcription factor binding sites at repressed promoters"

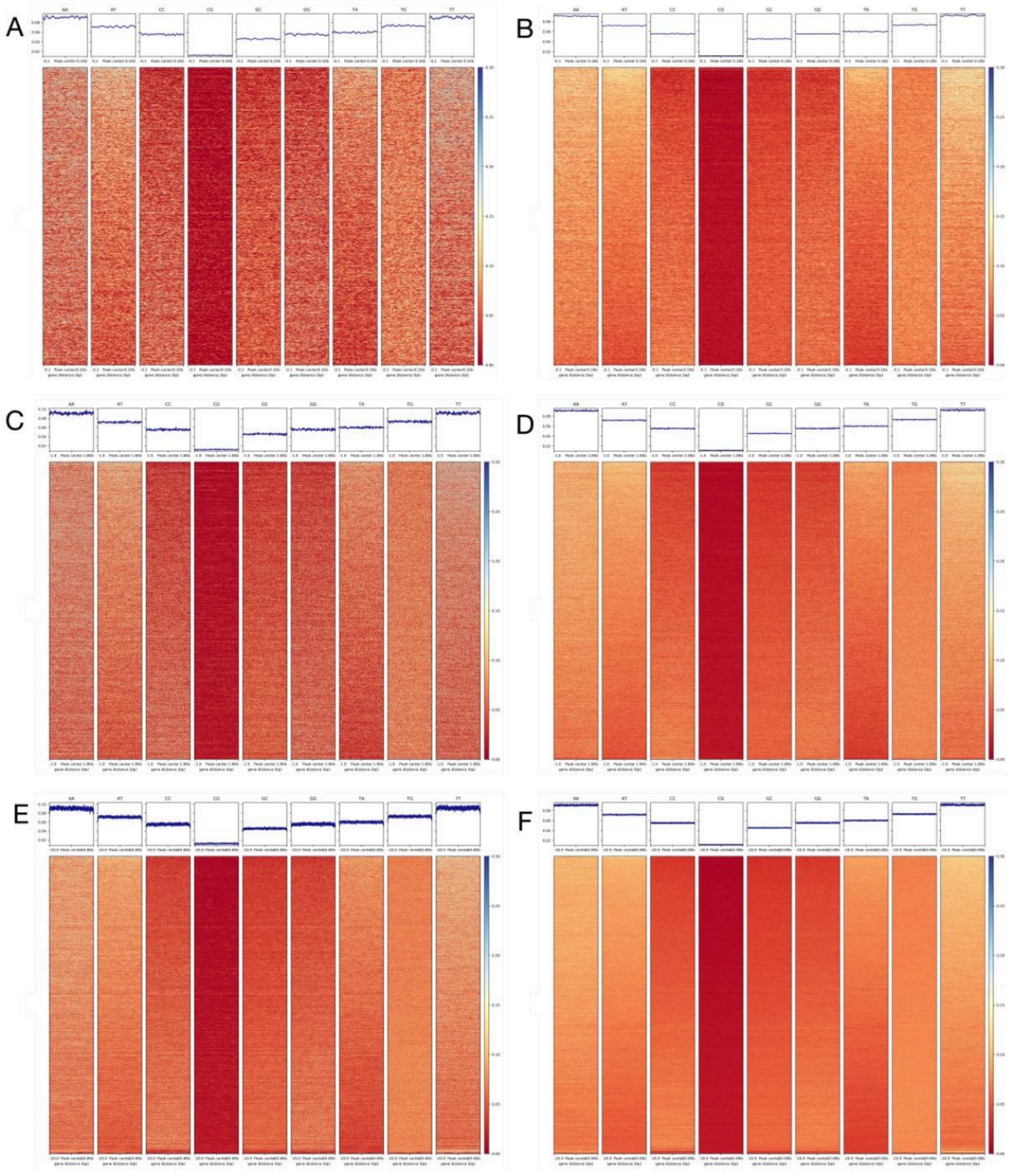


Additional file 4, fig 1: k-mer (dinucleotide) signals 150bp (A and B), 1kb (C and D) or 10kb (E and F) upstream and downstream from peak centers in ChIP-seq data GSE187851 (A, C and E) and ERR13661132 (B, D and F). The signals are plotted as mean and standard deviation of the mean.

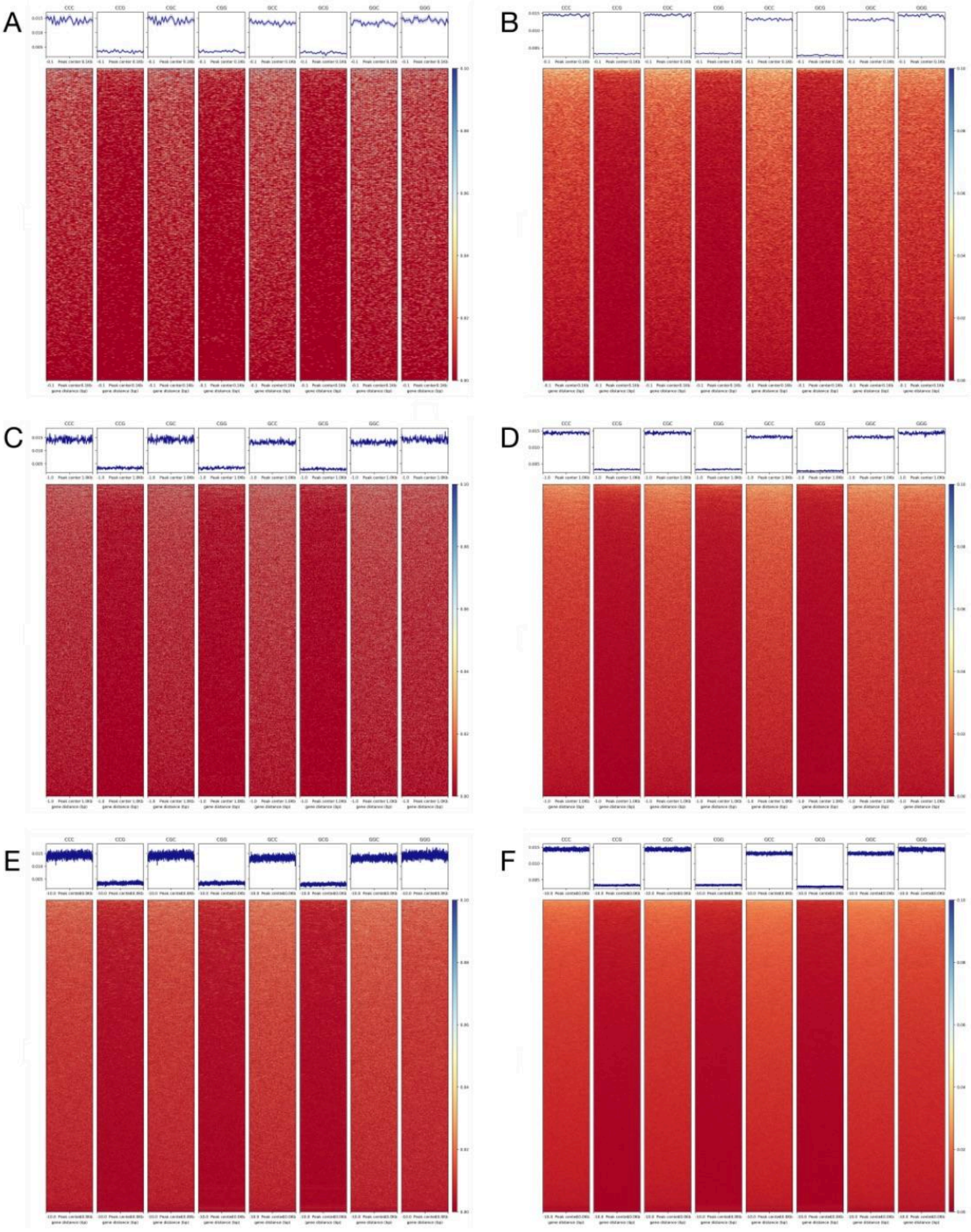


Additional file 4, fig 2: k-mer (trinucleotide) signals 150bp (A and B), 1kb (C and D) or 10kb (E and F) upstream and downstream from peak centers in ChIP-seq data GSE187851 (A, C and E) and ERR13661132 (B, D and F). The signals are plotted as mean and standard deviation of the mean.
