## Additional file 5 for "CGGBP1 from higher amniotes restricts cytosine methylation and drives a GC-bias in transcription factor binding sites at repressed promoters"

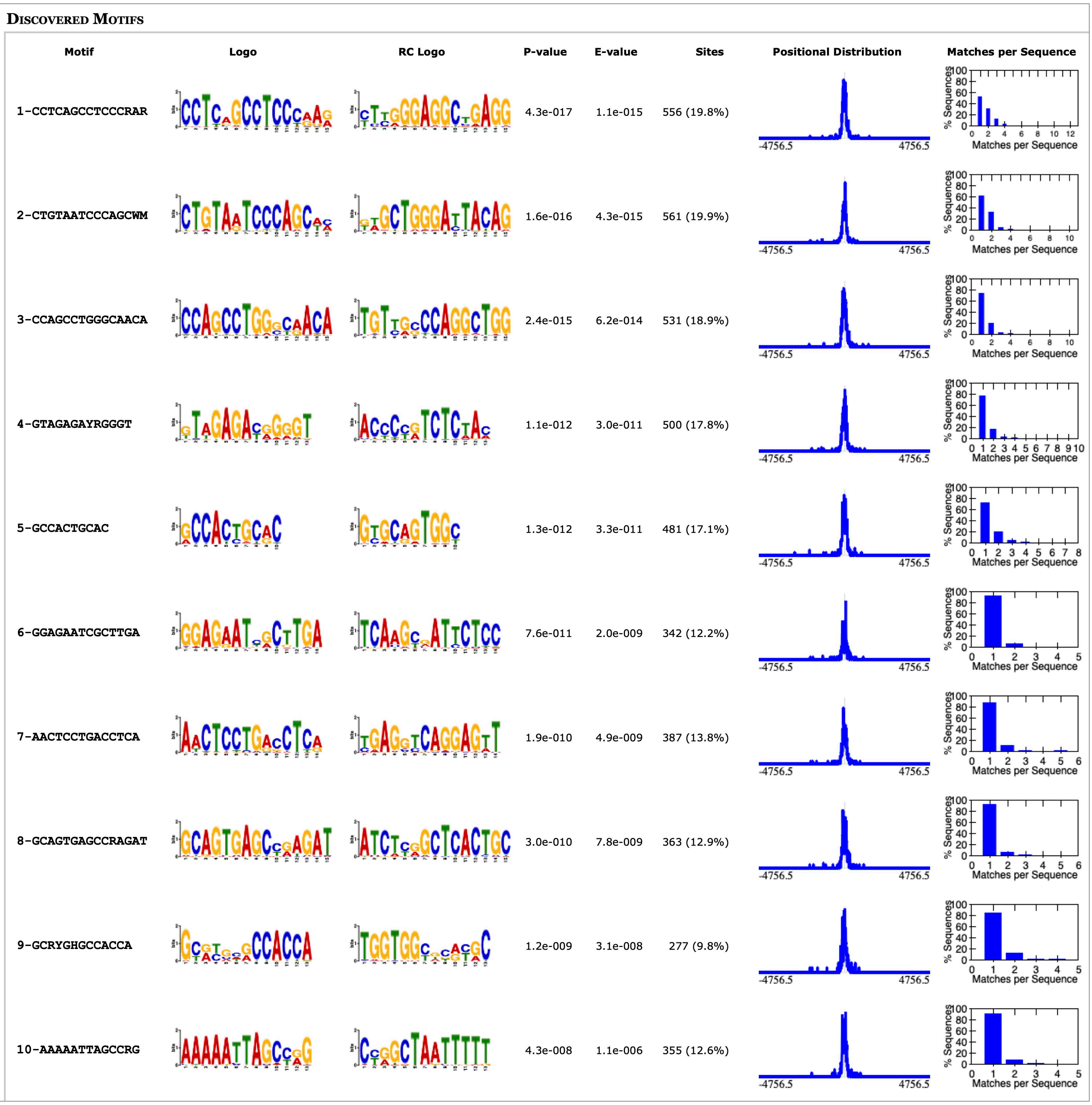


Additional file 5, fig 1: De-novo motifs identified using STREME in narrow peaks of ChIP vs Input (PMID: 22955616).

### Discovered Motifs

| Motif | P-value | E-value | Sites |
|---|---|---|---|
| 1-CCAGCCTGGGCAACA | 9.7e-163 | 2.2e-160 | 5389 (24.4%) |
| 2-GCTGGGATTACAGGC | 6.4e-161 | 1.4e-158 | 5578 (25.3%) |
| 3-CCTCAGCCTCCCRAR | 5.4e-160 | 1.2e-157 | 5437 (24.7%) |
| 4-GCCACCAYGCCYRGC | 6.4e-145 | 1.4e-142 | 5112 (23.2%) |
| 5-AGGAGAATCRCTTGA | 1.8e-141 | 4.0e-139 | 4790 (21.7%) |
| 6-GCAGTGAGCCRAGAT | 4.5e-131 | 9.9e-129 | 4175 (18.9%) |
| 7-CTCGAACTCCTGACC | 2.7e-129 | 6.1e-127 | 4507 (20.4%) |
| 8-CCRTCTCTACTAAAA | 2.0e-125 | 4.3e-123 | 4087 (18.5%) |
| 9-AGTGCAGTGGCRCR | 5.4e-112 | 1.2e-109 | 3912 (17.7%) |
| 10-AGYRAGACTCYGTC | 3.8e-108 | 8.4e-106 | 3874 (17.6%) |

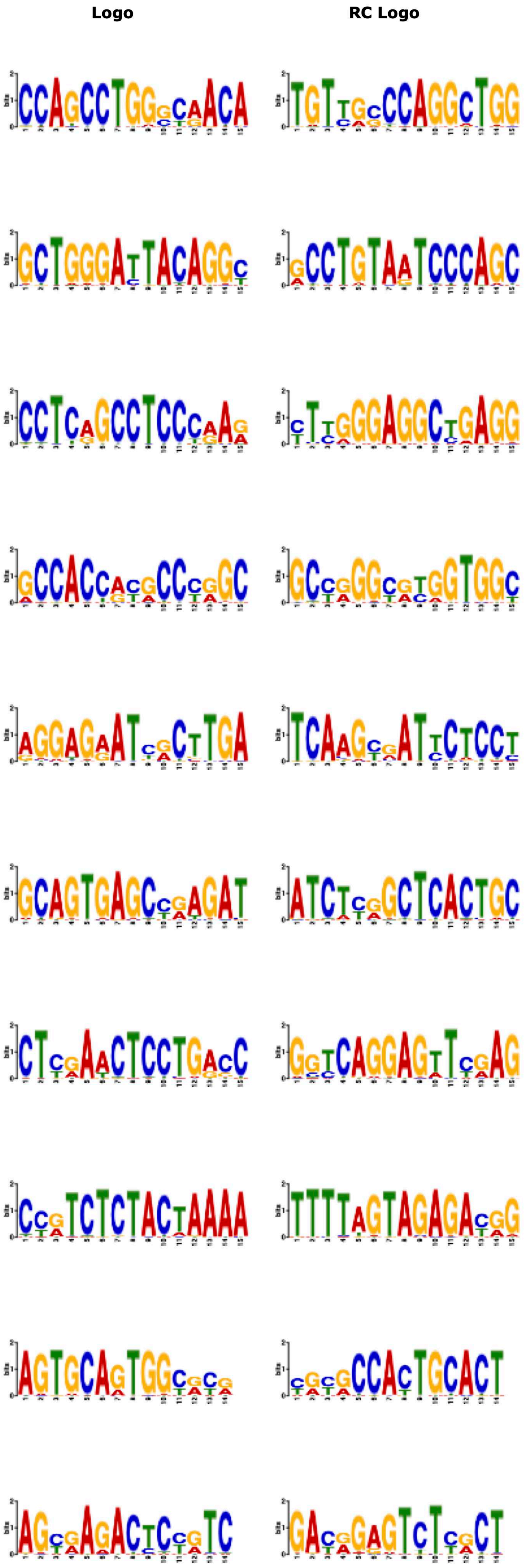


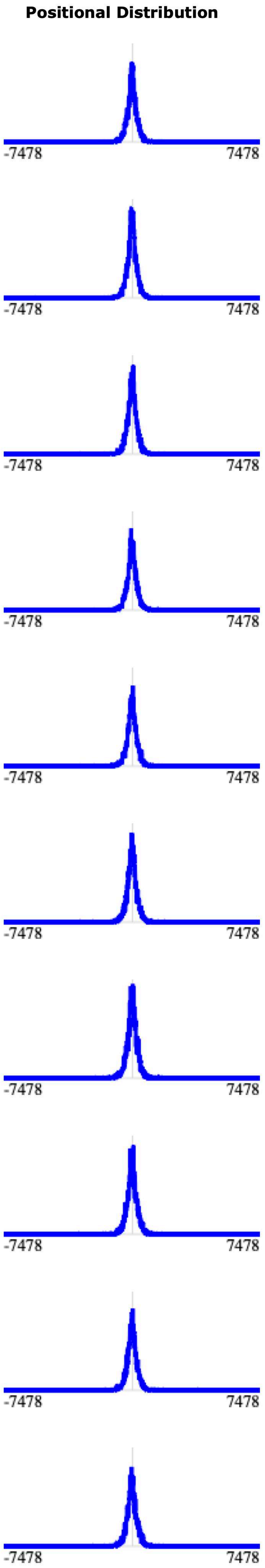


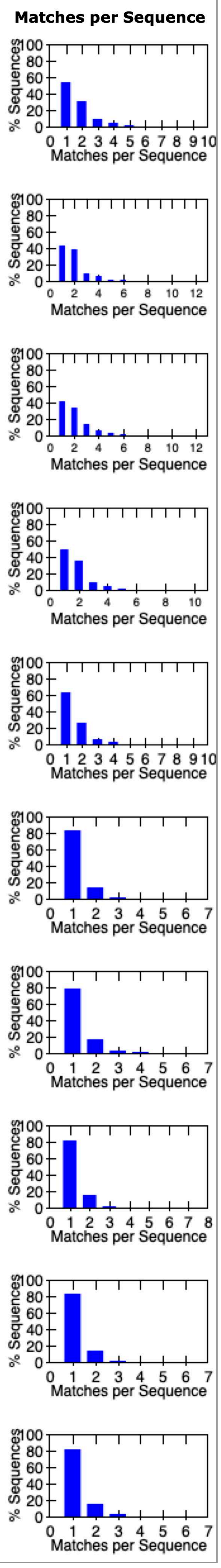


Additional file 5, fig 2: De-novo motifs identified using STREME in narrow peaks of ChIP rep1 vs Input (PMID: 39605320).

**Discovered Motifs**

| Motif | Logo | RC Logo | P-value | E-value | Sites | Positional Distribution | Matches per Sequence |
|---|---|---|---|---|---|---|---|
| 1-CTGGGATTACAGGCR | | | 4.2e-009 | 7.6e-008 | 285 (12.2%) | | |
| 2-CCAGCCTGGGCRACA | | | 2.3e-007 | 4.1e-006 | 270 (11.5%) | | |
| 3-CCTCAGCCTCCCRAR | | | 1.6e-006 | 3.0e-005 | 273 (11.7%) | | |
| 4-GCAGTGAGCCRAGAT | | | 3.3e-006 | 5.9e-005 | 175 (7.5%) | | |
| 5-GRATCRCTTGAR | | | 1.8e-005 | 3.2e-004 | 195 (8.3%) | | |
| 6-RTCTCTACTAAAAAT | | | 2.4e-005 | 4.3e-004 | 163 (7.0%) | | |
| 7-GTCTCAAAAAAAAA | | | 6.2e-005 | 1.1e-003 | 152 (6.5%) | | |
| 8-CTCGAACTCCTG | | | 9.0e-005 | 1.6e-003 | 186 (7.9%) | | |
| 9-CACCAYGCCYRGC | | | 4.4e-004 | 8.0e-003 | 276 (11.8%) | | |
| 10-AGTGCAGTGGYRC | | | 7.0e-004 | 1.3e-002 | 167 (7.1%) | | |

Additional file 5, fig 3: De-novo motifs identified using STREME in narrow peaks of ChIP rep2 vs Input (PMID: 39605320).
