## Additional file 6 for "CGGBP1 from higher amniotes restricts cytosine methylation and drives a GC-bias in transcription factor binding sites at repressed promoters"

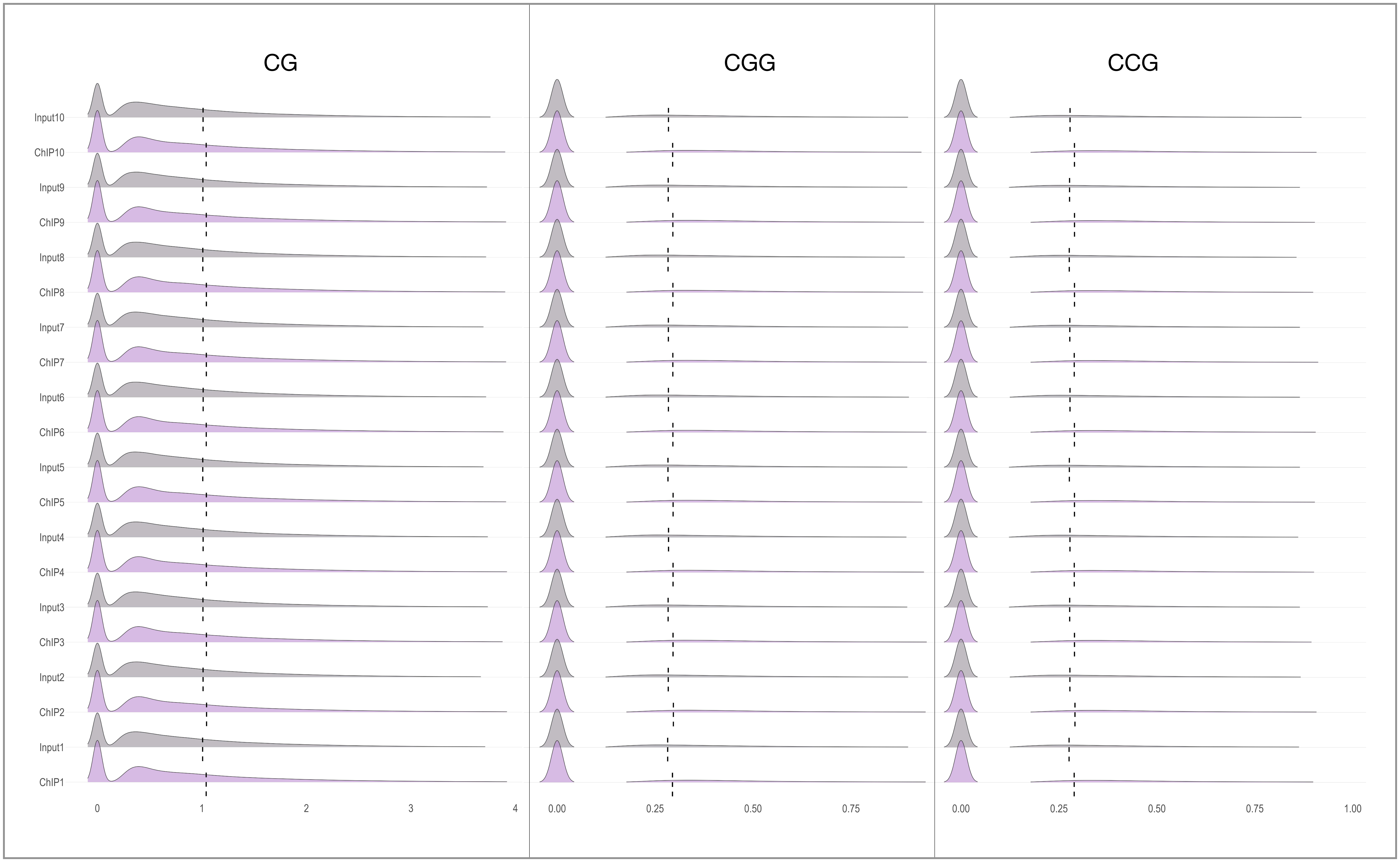


Additional file 6, fig 1: k-mer frequencies in randomly sampled 1 million reads from ChIP and Input (PMID: 39605320) over 10 independent iterations. The dotted lines show the means. The medians in these multimodal distributions are close to zero indicating the general paucity of these k-mers in the ChIP-seq and Input reads.

CG k-mer frequency analysis in random 1 million reads ChIP vs Input (PMID: 39605320)

| Pair | Mean ChIP | Mean Input | Median ChIP | Median Input | p-value | Significant | Test Performed |
|---|---|---|---|---|---|---|---|
| **ChIP1 vs Input1** | 1.04088358143253 | 1.00686130912804 | 0.680272108843537 | 0.674157303370786 | 1.51444121032334E-77 | Yes | Parametric Unpaired t-test |
| **ChIP2 vs Input2** | 1.04435828617597 | 1.01092749428045 | 0.680272108843537 | 0.677200902934537 | 1.04991024449284E-73 | Yes | Parametric Unpaired t-test |
| **ChIP3 vs Input3** | 1.04305614766619 | 1.00983298568487 | 0.680272108843537 | 0.675675675675676 | 4.11461751197382E-73 | Yes | Parametric Unpaired t-test |
| **ChIP4 vs Input4** | 1.04270476497347 | 1.01153161045208 | 0.680272108843537 | 0.675675675675676 | 1.15974500993924E-64 | Yes | Parametric Unpaired t-test |
| **ChIP5 vs Input5** | 1.04254256678485 | 1.00812410011454 | 0.680272108843537 | 0.675675675675676 | 1.12407818592124E-78 | Yes | Parametric Unpaired t-test |
| **ChIP6 vs Input6** | 1.04145276579996 | 1.01184095838028 | 0.680272108843537 | 0.677200902934537 | 1.39947462565872E-58 | Yes | Parametric Unpaired t-test |
| **ChIP7 vs Input7** | 1.04213807938901 | 1.01067360040536 | 0.680272108843537 | 0.677966101694915 | 5.31526177849543E-66 | Yes | Parametric Unpaired t-test |
| **ChIP8 vs Input8** | 1.04245338406195 | 1.00869094527666 | 0.680272108843537 | 0.675675675675676 | 1.48495413845044E-75 | Yes | Parametric Unpaired t-test |
| **ChIP9 vs Input9** | 1.04171547670381 | 1.00949199680579 | 0.680272108843537 | 0.675675675675676 | 5.89780336368967E-69 | Yes | Parametric Unpaired t-test |
| **ChIP10 vs Input10** | 1.04098785184545 | 1.01092686593721 | 0.680272108843537 | 0.675675675675676 | 4.13971850899907E-60 | Yes | Parametric Unpaired t-test |

Additional file 6, table 1: Unpaired t-test comparisons between ChIP and input for the occurrence of CG-mers; n = 1 million reads in each sampling.

CGG k-mer frequency analysis in random 1 million reads ChIP vs Input (PMID: 39605320)

| Pair | Mean ChIP | Mean Input | Median ChIP | Median Input | p-value | Significant | Test Performed |
|---|---|---|---|---|---|---|---|
| **ChIP1 vs Input1** | 0.29432077391269 | 0.282097133253989 | 0 | 0 | 9.03087024978376E-57 | Yes | Parametric Unpaired t-test |
| **ChIP2 vs Input2** | 0.296617675284415 | 0.283548289310366 | 0 | 0 | 8.15951494026756E-64 | Yes | Parametric Unpaired t-test |
| **ChIP3 vs Input3** | 0.295755273586543 | 0.283205214292923 | 0 | 0 | 5.01600384438823E-59 | Yes | Parametric Unpaired t-test |
| **ChIP4 vs Input4** | 0.295785870678016 | 0.284243304163105 | 0 | 0 | 4.93037871303606E-50 | Yes | Parametric Unpaired t-test |
| **ChIP5 vs Input5** | 0.295890246354339 | 0.28284618061992 | 0 | 0 | 1.66844642578297E-63 | Yes | Parametric Unpaired t-test |
| **ChIP6 vs Input6** | 0.295414370185227 | 0.283927622079972 | 0 | 0 | 1.18885014187528E-49 | Yes | Parametric Unpaired t-test |
| **ChIP7 vs Input7** | 0.29525183497008 | 0.283833365437294 | 0 | 0 | 2.68393383534222E-49 | Yes | Parametric Unpaired t-test |
| **ChIP8 vs Input8** | 0.295411599340409 | 0.283035632609593 | 0 | 0 | 1.72771384025251E-57 | Yes | Parametric Unpaired t-test |
| **ChIP9 vs Input9** | 0.295313913544289 | 0.283467290741997 | 0 | 0 | 1.09023013119488E-52 | Yes | Parametric Unpaired t-test |
| **ChIP10 vs Input10** | 0.294828266540746 | 0.284071858192011 | 0 | 0 | 1.03778876479542E-43 | Yes | Parametric Unpaired t-test |

Additional file 6, table 2: Unpaired t-test comparisons between ChIP and input for the occurrence of CGG-mers; n = 1 million reads in each sampling.

CCG k-mer frequency analysis in random 1 million reads ChIP vs Input (PMID: 39605320)

| Pair | Mean ChIP | Mean Input | Median ChIP | Median Input | p-value | Significant | Test Performed |
|---|---|---|---|---|---|---|---|
| **ChIP1 vs Input1** | 0.288843265107831 | 0.275757514994074 | 0 | 0 | 3.1166013442723E-65 | Yes | Parametric Unpaired t-test |
| **ChIP2 vs Input2** | 0.290812813920599 | 0.277881536901816 | 0 | 0 | 2.22415977574053E-62 | Yes | Parametric Unpaired t-test |
| **ChIP3 vs Input3** | 0.289591423384334 | 0.277335949020029 | 0 | 0 | 1.92677717365597E-56 | Yes | Parametric Unpaired t-test |
| **ChIP4 vs Input4** | 0.28903658646431 | 0.278057779289386 | 0 | 0 | 5.51093390398463E-46 | Yes | Parametric Unpaired t-test |
| **ChIP5 vs Input5** | 0.289542451080857 | 0.276478369004715 | 0 | 0 | 1.69126789799733E-64 | Yes | Parametric Unpaired t-test |
| **ChIP6 vs Input6** | 0.289396773132604 | 0.278158557936177 | 0 | 0 | 6.30299165041079E-48 | Yes | Parametric Unpaired t-test |
| **ChIP7 vs Input7** | 0.288989770334573 | 0.277637196850938 | 0 | 0 | 4.10736802498036E-49 | Yes | Parametric Unpaired t-test |
| **ChIP8 vs Input8** | 0.289619415092553 | 0.276275811860443 | 0 | 0 | 1.0159311351746E-66 | Yes | Parametric Unpaired t-test |
| **ChIP9 vs Input9** | 0.289650951162358 | 0.277013405707154 | 0 | 0 | 4.28637940444377E-60 | Yes | Parametric Unpaired t-test |
| **ChIP10 vs Input10** | 0.289541191158927 | 0.278290234579649 | 0 | 0 | 1.13967248808805E-47 | Yes | Parametric Unpaired t-test |

Additional file 6, table 3: Unpaired t-test comparisons between ChIP and input for the occurrence of CCG-mers; n = 1 million reads in each sampling.

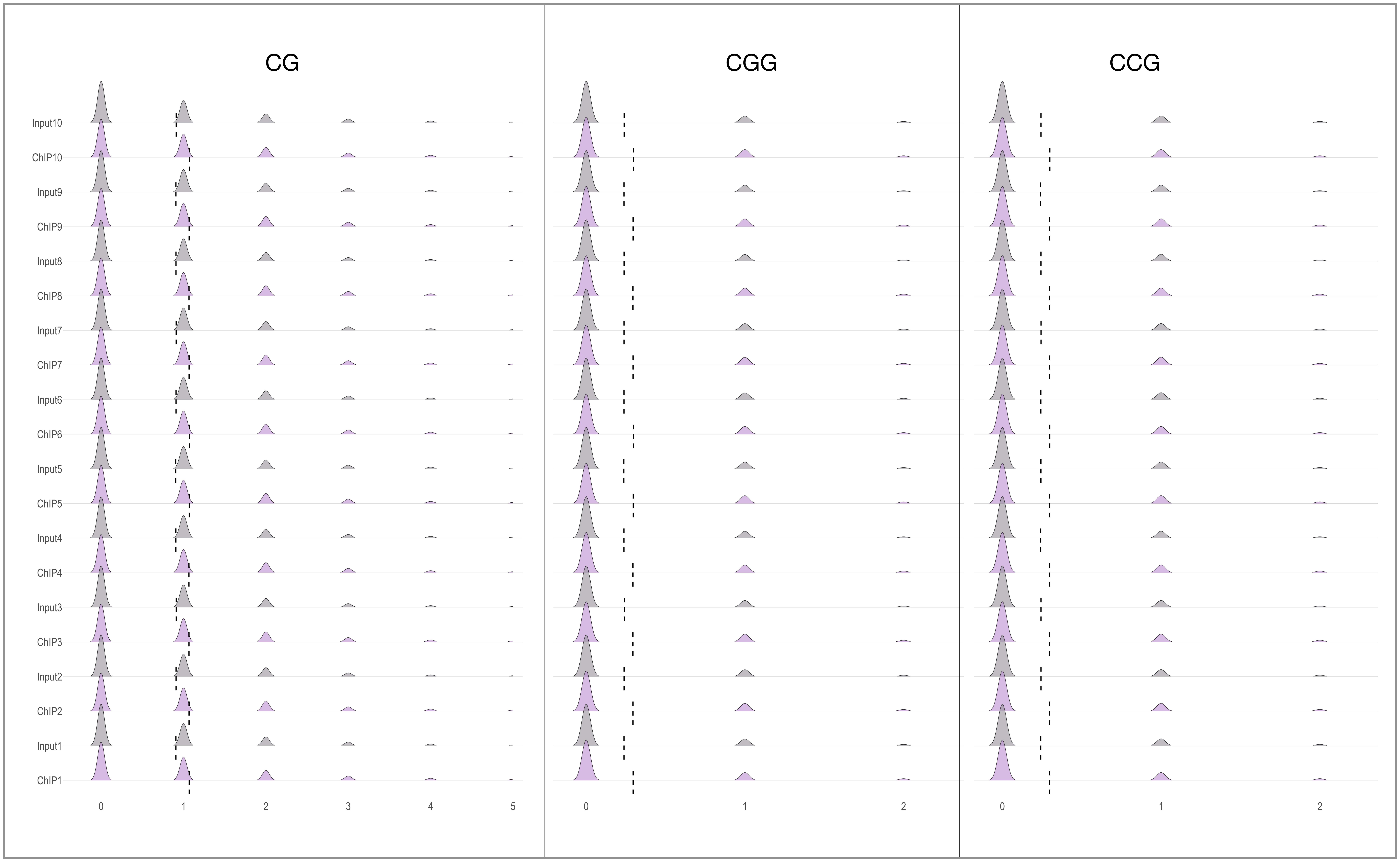


Additional file 6, fig 2: k-mer frequencies in randomly sampled 1 million reads from ChIP and Input (PMID: 22955616) over 10 independent iterations. The dotted lines show the means. The medians in these multimodal distributions are close to zero indicating the general paucity of these k-mers in the ChIP-seq and Input reads.

CG k-mer frequency analysis in random 1 million reads ChIP vs Input (PMID: 22955616)

| Pair | Mean ChIP | Mean Input | Median ChIP | Median Input | p-value | Significant | Test Performed |
|---|---|---|---|---|---|---|---|
| **ChIP1 vs Input1** | 1.06870270931601 | 0.907873768365778 | 1 | 0 | 0 | Yes | Parametric Unpaired t-test |
| **ChIP2 vs Input2** | 1.06707618536518 | 0.909251903508788 | 1 | 0 | 0 | Yes | Parametric Unpaired t-test |
| **ChIP3 vs Input3** | 1.06634756758584 | 0.91105829552345 | 1 | 0 | 0 | Yes | Parametric Unpaired t-test |
| **ChIP4 vs Input4** | 1.06599914757196 | 0.907282923963523 | 1 | 0 | 0 | Yes | Parametric Unpaired t-test |
| **ChIP5 vs Input5** | 1.06917071799213 | 0.905841800987181 | 1 | 0 | 0 | Yes | Parametric Unpaired t-test |
| **ChIP6 vs Input6** | 1.06997895731864 | 0.908857874609401 | 1 | 0 | 0 | Yes | Parametric Unpaired t-test |
| **ChIP7 vs Input7** | 1.06752958124866 | 0.910532248664532 | 1 | 0 | 0 | Yes | Parametric Unpaired t-test |
| **ChIP8 vs Input8** | 1.06655359842022 | 0.908428482020252 | 1 | 0 | 0 | Yes | Parametric Unpaired t-test |
| **ChIP9 vs Input9** | 1.06813409619257 | 0.908029895118872 | 1 | 0 | 0 | Yes | Parametric Unpaired t-test |
| **ChIP10 vs Input10** | 1.07065057261081 | 0.910836457467157 | 1 | 0 | 0 | Yes | Parametric Unpaired t-test |

Additional file 6, table 4: Unpaired t-test comparisons between ChIP and input for the occurrence of CG-mers; n = 1 million reads in each sampling.

CGG k-mer frequency analysis in random 1 million reads ChIP vs Input (PMID: 22955616)

| Pair | Mean ChIP | Mean Input | Median ChIP | Median Input | p-value | Significant | Test Performed |
|---|---|---|---|---|---|---|---|
| **ChIP1 vs Input1** | 0.295342901887829 | 0.23806960221655 | 0 | 0 | 0 | Yes | Parametric Unpaired t-test |
| **ChIP2 vs Input2** | 0.294637556620886 | 0.238535508522204 | 0 | 0 | 0 | Yes | Parametric Unpaired t-test |
| **ChIP3 vs Input3** | 0.294392647578179 | 0.239508100254735 | 0 | 0 | 0 | Yes | Parametric Unpaired t-test |
| **ChIP4 vs Input4** | 0.294513852326721 | 0.237591400996342 | 0 | 0 | 0 | Yes | Parametric Unpaired t-test |
| **ChIP5 vs Input5** | 0.295841867978988 | 0.237133669009456 | 0 | 0 | 0 | Yes | Parametric Unpaired t-test |
| **ChIP6 vs Input6** | 0.296296677272464 | 0.237983163580067 | 0 | 0 | 0 | Yes | Parametric Unpaired t-test |
| **ChIP7 vs Input7** | 0.294902598518234 | 0.238155296337567 | 0 | 0 | 0 | Yes | Parametric Unpaired t-test |
| **ChIP8 vs Input8** | 0.294541336839851 | 0.238437544838546 | 0 | 0 | 0 | Yes | Parametric Unpaired t-test |
| **ChIP9 vs Input9** | 0.295070069682153 | 0.238026027375618 | 0 | 0 | 0 | Yes | Parametric Unpaired t-test |
| **ChIP10 vs Input10** | 0.296563007074134 | 0.239174247209203 | 0 | 0 | 0 | Yes | Parametric Unpaired t-test |

Additional file 6, table 5: Unpaired t-test comparisons between ChIP and input for the occurrence of CGG-mers; n = 1 million reads in each sampling.

CCG k-mer frequency analysis in random 1 million reads ChIP vs Input (PMID: 22955616)

| Pair | Mean ChIP | Mean Input | Median ChIP | Median Input | p-value | Significant | Test Performed |
|---|---|---|---|---|---|---|---|
| **ChIP1 vs Input1** | 0.298089445179518 | 0.242002127751768 | 0 | 0 | 0 | Yes | Parametric Unpaired t-test |
| **ChIP2 vs Input2** | 0.29788693379568 | 0.243243956409611 | 0 | 0 | 0 | Yes | Parametric Unpaired t-test |
| **ChIP3 vs Input3** | 0.29745182371315 | 0.243266081790854 | 0 | 0 | 0 | Yes | Parametric Unpaired t-test |
| **ChIP4 vs Input4** | 0.296577609585546 | 0.241844518902874 | 0 | 0 | 0 | Yes | Parametric Unpaired t-test |
| **ChIP5 vs Input5** | 0.298164185603955 | 0.24286495167776 | 0 | 0 | 0 | Yes | Parametric Unpaired t-test |
| **ChIP6 vs Input6** | 0.298475066118889 | 0.242438541914306 | 0 | 0 | 0 | Yes | Parametric Unpaired t-test |
| **ChIP7 vs Input7** | 0.298231095931709 | 0.243492817774624 | 0 | 0 | 0 | Yes | Parametric Unpaired t-test |
| **ChIP8 vs Input8** | 0.297622656394522 | 0.243126952777319 | 0 | 0 | 0 | Yes | Parametric Unpaired t-test |
| **ChIP9 vs Input9** | 0.29816071930099 | 0.241438530671944 | 0 | 0 | 0 | Yes | Parametric Unpaired t-test |
| **ChIP10 vs Input10** | 0.299126286480242 | 0.243200975352108 | 0 | 0 | 0 | Yes | Parametric Unpaired t-test |

Additional file 6, table 6: Unpaired t-test comparisons between ChIP and input for the occurrence of CCG-mers; n = 1 million reads in each sampling.
