## Additional supplementary information file for "CGGBP1 from higher amniotes restricts cytosine methylation and drives a GC-bias in transcription factor binding sites at repressed promoters": CGGBP1_constructs_design.pdf

### ***Homo sapiens* (Hs)**

**Insert name: Hs**
**Uniprot ID:Q9UFW8**
**N-terminal cloning site: KpnI**
**C-terminal cloning site: XhoI**
**Size: 555 bp**
**Sequence including restriction sites:**

5’ - GGTACC(KpnI)GCCGCCACC(Kozak)ATG(start
codon)GGATATCCTTACGATGTACCAGACTATGCT(HA-Tag)GAGCGATTTGTAGTAACAGCA
CCACCTGCTCGAAACCGTTCTAAGACTGCTTTGTATGTGACTCCCCTGGATCGAGTCACTGAGT
TTGGAGGTGAGCTGCATGAAGATGGAGGAAAACTCTTCTGCACTTCTTGCAATGTGGTTCTGAA
TCATGTTCGCAAGTCTGCCATTAGTGACCACCTCAAGTCAAAGACTCATACCAAGAGGAAGGCA
GAATTTGAAGAGCAGAATGTGAGAAAGAAGCAGAGGCCCCTAACTGCATCTCTTCAGTGCAACA
GTACTGCGCAAACAGAGAAAGTCAGTGTTATCCAGGACTTTGTGAAAATGTGCCTGGAAGCCAA
CATCCCACTTGAGAAGGCTGATCACCCAGCAGTCCGTGCTTTCCTATCTCGCCATGTGAAGAAT
GGAGGCTCCATACCTAAGTCAGACCAGCTACGGAGGGCATATCTTCCTGATGGATATGAGAATG
AGAATCAACTCCTCAACTCACAAGATTGTTGA(Stop codon)CTCGAG(XhoI) - 3’

### ***Gallus gallus* (Gg)**

**Insert name: Gg**
**Uniprot ID: A0A1D5NUG1**
**N-terminal cloning site: KpnI**
**C-terminal cloning site: XhoI**
**Insert Size: 576 bp**
**Sequence including restriction sites:**

5’ - GGTACC (KpnI)GCCGCCACC (Kozak)ATG (start
codon)GAACGGTTCGGGGTGAAGTCCACTCCGTCACGTAACCGCTCGAAGACTGCTTTGTACG
TAACTCCTCAGGATCGGGTAACTGAGTTTGGCAGCGAGCTGCATGAAGATGGCGGAAAACTCTT
CTGCACTTCCTGCAACGTGGTCCTGAATCACGTCCGTAAATCGGCAATCAACGATCATCTTAAG
TCAAAAACGCATACGAAGCGGAAGGCGGAGTTTGAAGAGCAGAACGTCAGGAAGAAGCAGAGGA
CTCTGACCGCCTCCCTTCAGTGCAACAGCACTGCTCAGACGGAAAAGACTAGCGTCATACAAGA
CTTTGTGAAAATGTGCCTGGAAGCTAATATTCCCCTTGAGAAGGCTGACCATCCTTCTGTGCGA
GCCTTCCTGTCTCGCTACGTCAAAAACGGGAGTTCGATCCCCAAATCGGAGCAGTTAAGGAAAG
CATACCTGCCCGATGGGTATGACAATGAGAATCAACTCATCAATACCGAAGATCGC (stop
codon removed)GGG (linker glycine)GACTACAAAGACGATGACGACAAG
GACTACAAAGACGATGACGACAAG (FLAG 2x)TGA (stop codon)CTCGAG (XhoI)-
3’

### ***Latimeria chalumnae* (Lc)**

**Insert name: Lc**
**Uniprot ID:H3AJH5**
**N-terminal cloning site: KpnI**
**C-terminal cloning site: XhoI**
**Insert Size: 576 bp**
**Sequence including restriction sites:**

5′ - GGTACC (KpnI)GCCGCCACC (Kozak)ATG (start
codon)GAAAAGTCTGTGAAGAAGTCTGCCTTTTCTGGCCGTAAACGCAACAAAACTGCCTTGT
ATATAACAGCCCAGCAACGGTGTGAGCAGTTTGGGCCTCAGCTCCATGAGGATGGAGGAAAACT
CTTCTGCAGCGCGTGTAATGTGGTGCTAGACCATATCCGTAAATCGACGATCACCGATCATCTT
CGATCAAAGACCCACATGAAGAGGCTGGCAGAGTTTTCAGAAGAGTGCGTCAGGAAGAAGCAGA
AGACTTTAACGACTTCGTTGCAGTGCAACACTGTGGCTCAGGTGGAAAAGATAAGCGTCGTTCA
AGATTTTGTGAAAATGTGCCTGGAGGCAGGTATACCCCTGGAAAAAGCCAACCACCCATCGGTT
TGCACCTTTCTTTTGGCTCACGTCAAAAACAGTGATATCCCTCCATCTGACCAACTGAGAAAAG
TTGGCCTTCCTGATGTGCATAAAGCAAAACAAATATGTGTAAACCAAAAGTTTGTG (stop
codon removed)GGG (linker glycine)GACTACAAAGACGATGACGACAAG
GACTACAAAGACGATGACGACAAG (FLAG 2x)TGA (stop codon)CTCGAG (XhoI)-
3′

### ***Anolis carolinensis* (Ac)**

**Insert name: Ac**
**Uniprot ID:G1KEH3**
**N-terminal cloning site: KpnI**
**C-terminal cloning site: XhoI**
**Insert Size: 660 bp**
**Sequence including restriction sites:**

5′ - GGTACC (KpnI)GCCGCCACC (Kozak)ATG (start
codon)GAAGGTGATTCCTGTGGGATTCCTGAGCTTTTCCCCGTCACACCCATGGAGTCGTGGA
TGGAGTGTAACAGCTTACTGAAGACAACCATGGATCGGTTCGACGTGAAGCCGCCTCCGTCTCG
GAGCCGCTCTAAGACTGCTTTGTACGTGACGCCTCAGGACCGCGTCACGGAGTTTGGCAGCGAA
CTGTATGAAGACGGGGGGAAGTTGTATTGCACTTTCTGCAACGTGGTCTTGAACCACGTCCGAA
AATCCGCTATCAACGATCATCTGAAATCCAAGACTCACACCAAGCGGAAGGGGGAGTTTGAAGA
GCAAACCGTGCGGAAGAAGCCAAGGACCCTGACGGCCTCTCTGCAGTGCAACAGCGCGCCCCAG
ATTGAAAAGCCGAACGTCGTTCACGACTTTGTAAAAATGTTTCTGGAGGCCAGCATTCCCCTTG

AGAAGGCCGACCATCCCGCCGTGCGAGCGTTCCTCTCTCGGCACGTAAAAAACGGGAATTCTGT
CCCCAAAGCGGAGCAGCTCAGGAAGGCCTACCTGCCTGATGGGTTTACAAATCAGATGATCAAA
TCTGAAGACCAC (stop codon removed)GGG (linker
glycine)GACTACAAAGACGATGACGACAAG GACTACAAAGACGATGACGACAAG (FLAG
2x)TGA (stop codon)CTCGAG (XhoI)- 3’

### ***Rhinatrema bivittatum*** **(Rh)**

**Insert name: Rh**
**Uniprot ID: NA**
**N-terminal cloning site: KpnI**
**C-terminal cloning site: XhoI**
**Insert Size: 915 bp**
**Sequence including restriction sites:**

5’- GGTACC(KpnI)GCCGCCACC(Kozak)ATG(start
codon)CAGGCAGAGAAAGAACGTGTGGATATTTGTAAAGCATGGGTGAAGACACTCGCAGGTGCCAAT
ATCCCACTGTCCAAAATTGAACACCCACTTGTAAGAGAATTTCTCAACAGCAGGGTTCGGAATGGTGGT
GCAATTCCTGGAAGGACCCAACTGACTGAGACATACTTACCTGAAGTCTACCATGAAGAAAAAGAAAAG
TTACAACTGATGCTTAAAGATCAAAAAGTTGCAATTATCTTTGATGAAACATGTGATGATGATGCAAGA
TCAGTTCTGAATGTATTGTTTGCACCATTGGAACCTGATTGCACTGGGCATGTAAAATCATTTTTGGCA
AATATAGTGTTTTTAGATGTTGTTAATCACTTAATTGTTGCCCAAGCTGTAGTCAAAGCGATAAATGGT
TATCAGATTGATTACAACAGCAATTTGGTAATCGATACTGATAATGATAGTTATATGGAAAAAGCTTTT
AATTCTGTGCTCTCTACACTGTTTCCAAATTCTGTGCATATCACTTGTTTGGCACATATTGTTAACTTA
GTTGGTGAATCATTCAGAAAGCCTTTTCAGTTAATTGATACATTTGTAAGGAGCTTCAAAAACATGTTC
TATAACTCTAGTAGTAGGAAAGCGAAGTACCTGCGTTTCTTGAACAAAAAAGTACAAGAGAAATCTCTA
GCTGCTAAAGCATCCATGCCTCCAAGTCCTGTCGGAATACATTGGAACAGCTGGTTCCAATCTATTCAG
TATCATGTTAAGAATTTTCACTTTTACAAAGAATTCTTCATTGAAGAGTGTAACAGTAACTCCCCAGTT
TCTATGCACACTATT( Stop codon removed)GGG(linker
glycine)GACTACAAAGACGATGACGACAAGGACTACAAAGACGATGACGACAAG(FLAG2x)TAG
(Stop codon)CTCGAG(XhoI)-3’
