## Supplementary table for "CGGBP1 from higher amniotes restricts cytosine methylation and drives a GC-bias in transcription factor binding sites at repressed promoters": Table_S16.pdf

Table S16: Details of hypomethylated TFBSs (low MeDIP in Hs)

| Motif ID | Motif Name | GC content | Function | JASPAR vertebrate motif diagram |
|---|---|---|---|---|
| **MA0039.4** | KLF4 | 72.9239 | Activator or Repressor | 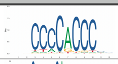 |
| **MA0073.1** | RREB1 | 62.2727 | Transcriptional activator | 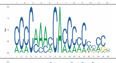 |
| **MA0079.5** | SP1 | 91.3502 | Transcriptional activator or repressor | 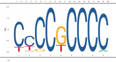 |
| **MA0080.6** | SPI1 | 45.688 | Transcriptional activator | 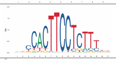 |
| **MA0081.2** | SPIB | 46.2878 | Transcriptional activator | 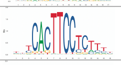 |
| **MA0162.4** | EGR1 | 75.1882 | Transcriptional activator | 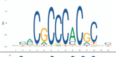 |
| **MA0493.2** | KLF1 | 83.7886 | Transcriptional activator | 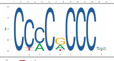 |
| **MA0501.1** | MAF::NFE2 | 42.9358 | Transcriptional activator or repressor | 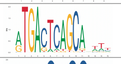 |
| **MA0516.3** | SP2 | 90.1108 | Transcriptional activator | 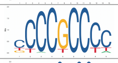 |
| **MA0599.1** | KLF5 | 79.2131 | Transcriptional activator | 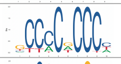 |
| **MA0623.2** | NEUROG1 | 40.7295 | Transcriptional activator | 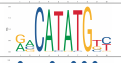 |
| **MA0685.2** | SP4 | 88.3852 | Transcriptional activator | 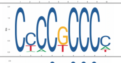 |
| **MA0740.2** | KLF14 | 87.1227 | Transcriptional repressor | 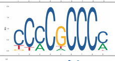 |
| **MA0741.1** | KLF16 | 78.4208 | Regulation of transcription | 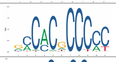 |
| **MA0742.2** | KLF12 | 91.9625 | Transcriptional repressor | 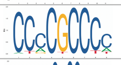 |
| **MA0746.2** | SP3 | 74.4037 | Activator or Repressor | 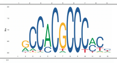 |
| **MA0747.1** | SP8 | 70.3579 | Transcriptional activator | 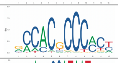 |
| **MA0752.1** | ZNF410 | 38.1187 | Regulation of transcription | 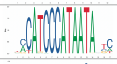 |
| **MA0753.2** | ZNF740 | 79.3626 | Regulation of transcription | 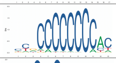 |
| **MA1099.2** | HES1 | 73.6828 | Transcriptional repressor | 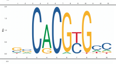 |
| **MA1107.2** | KLF9 | 66.1275 | Transcriptional activator | 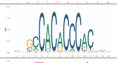 |
| **MA1116.1** | RBPJ | 52.8393 | Transcriptional repressor | 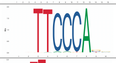 |
| **MA1421.1** | TCF7L1 | 35.3753 | Transcriptional activator or repressor | 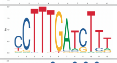 |
| **MA1511.2** | KLF10 | 90.6329 | Transcriptional repressor | 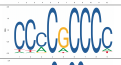 |
| **MA1512.1** | KLF11 | 74.933 | Transcriptional activator or repressor | 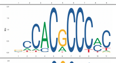 |
| **MA1513.1** | KLF15 | 90.155 | Transcriptional repressor | 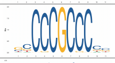 |
| **MA1564.1** | SP9 | 73.6818 | Transcriptional activator | 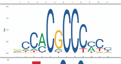 |
| **MA1566.2** | TBX3 | 46.737 | Transcriptional repressor | 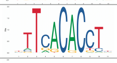 |
| **MA1627.1** | WT1 | 70.314 | Regulation of transcription | 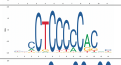 |
| **MA1630.2** | ZNF281 | 83.261 | Transcriptional repressor | 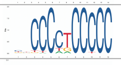 |
| **MA1640.1** | MEIS2 | 40.6485 | Transcriptional activator | 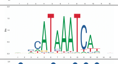 |
| **MA1959.1** | KLF7 | 85.3382 | Transcriptional repressor | 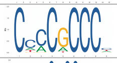 |
| **MA1961.1** | PATZ1 | 88.8517 | Regulation of transcription | 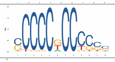 |
| **MA1971.1** | ZBED2 | 54.1855 | Transcriptional repressor | 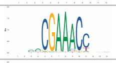 |
| **MA1976.1** | ZNF320 | 66.6178 | Regulation of transcription | 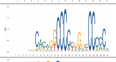 |
| **MA1986.1** | ZNF692 | 77.7124 | Transcriptional repressor | 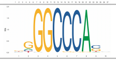 |
