## Supplementary material for "CGGBP1 from higher amniotes restricts cytosine methylation and drives a GC-bias in transcription factor binding sites at repressed promoters": Raw data file

Raw images corresponding to Fig 1B

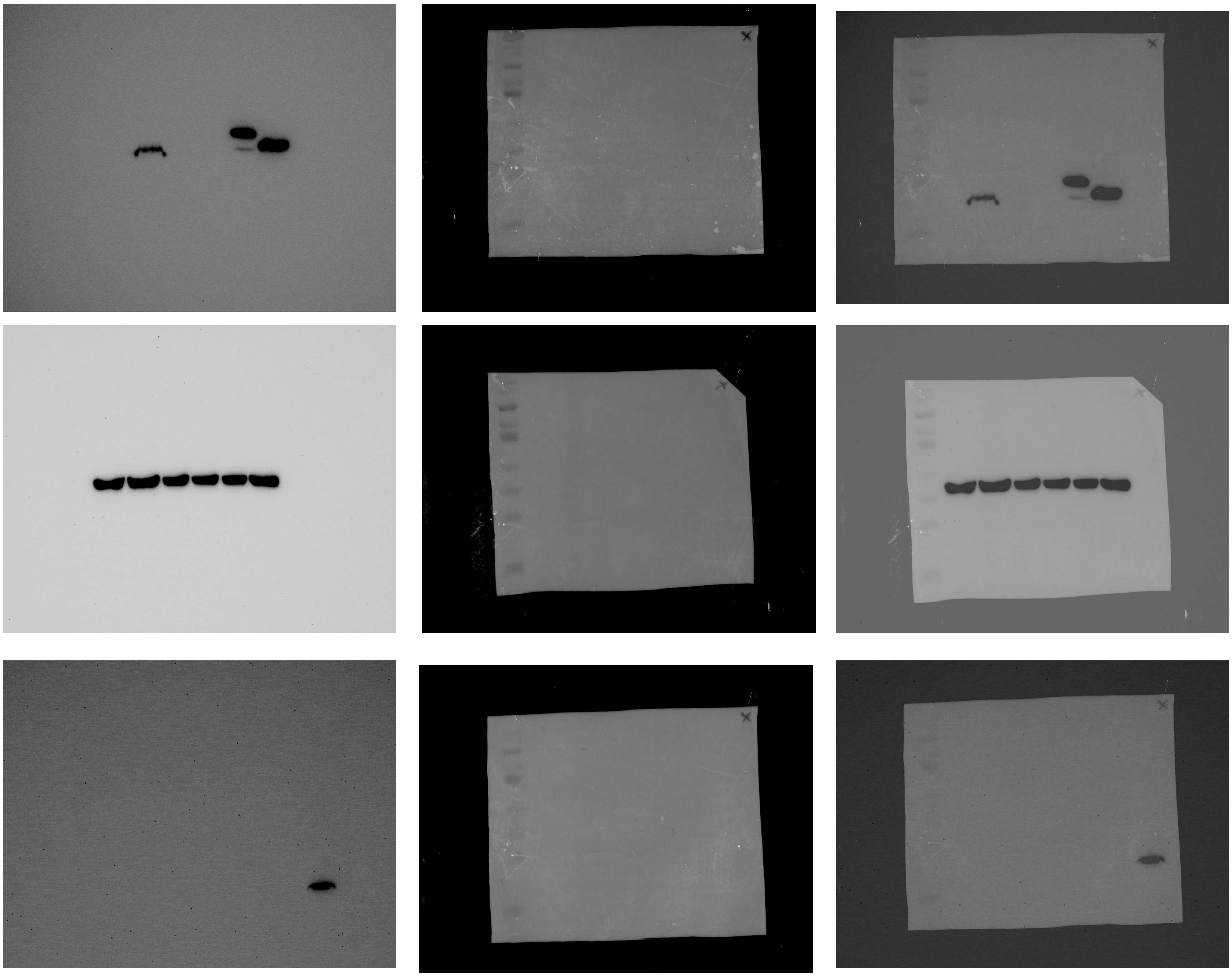

Raw images corresponding to Fig S1A

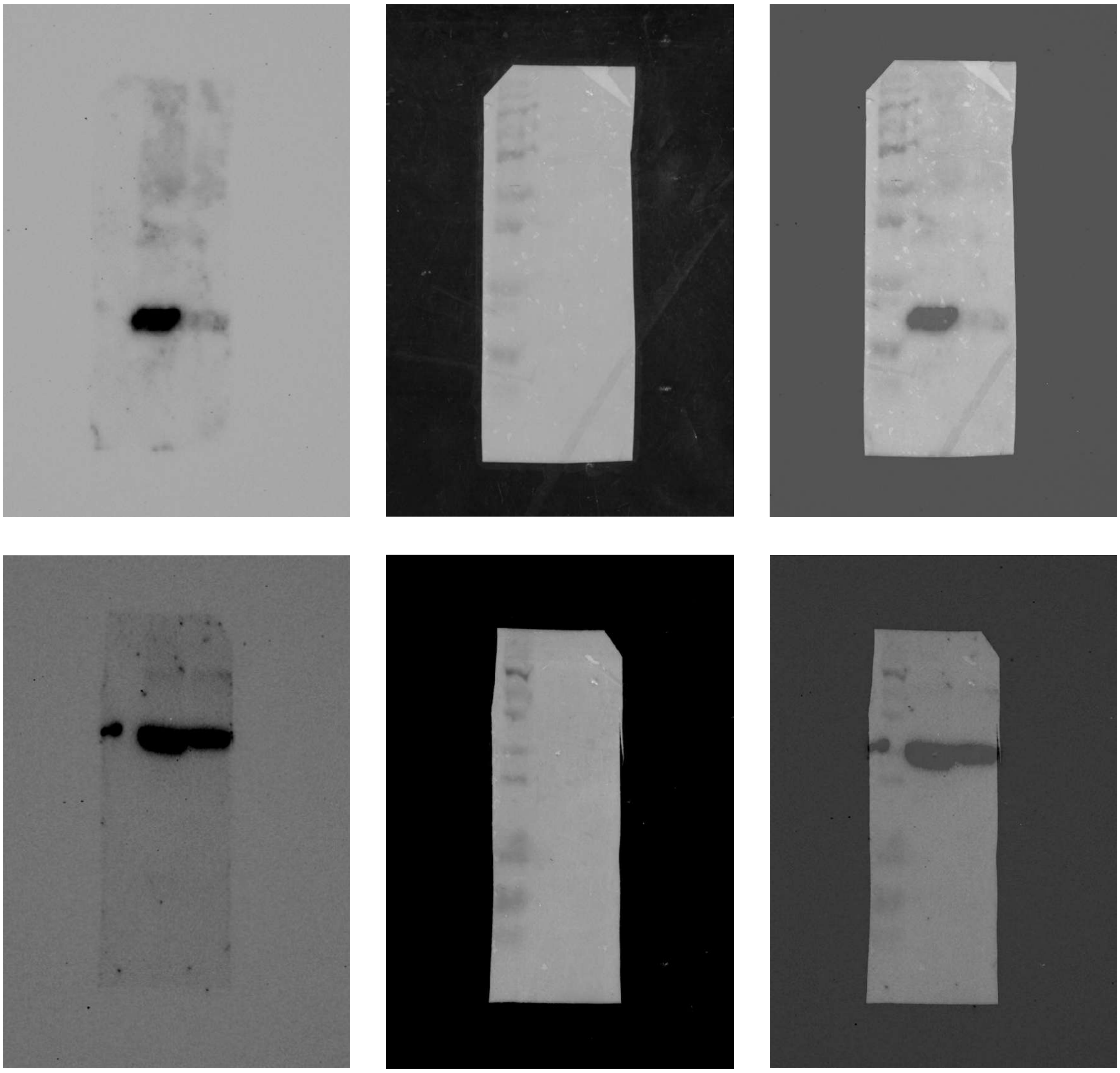

Raw images corresponding to Fig S1B

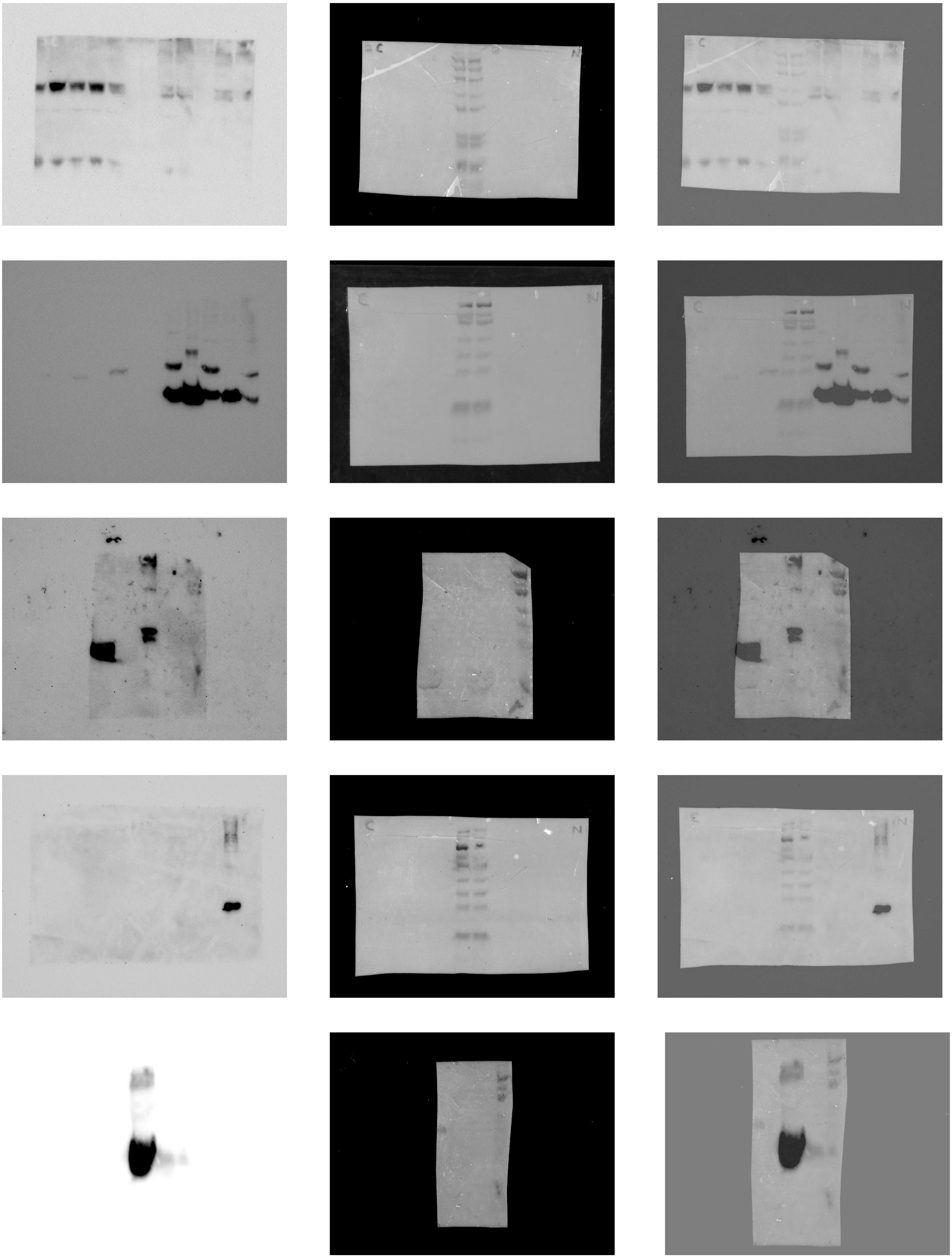

Raw images corresponding to Fig S3

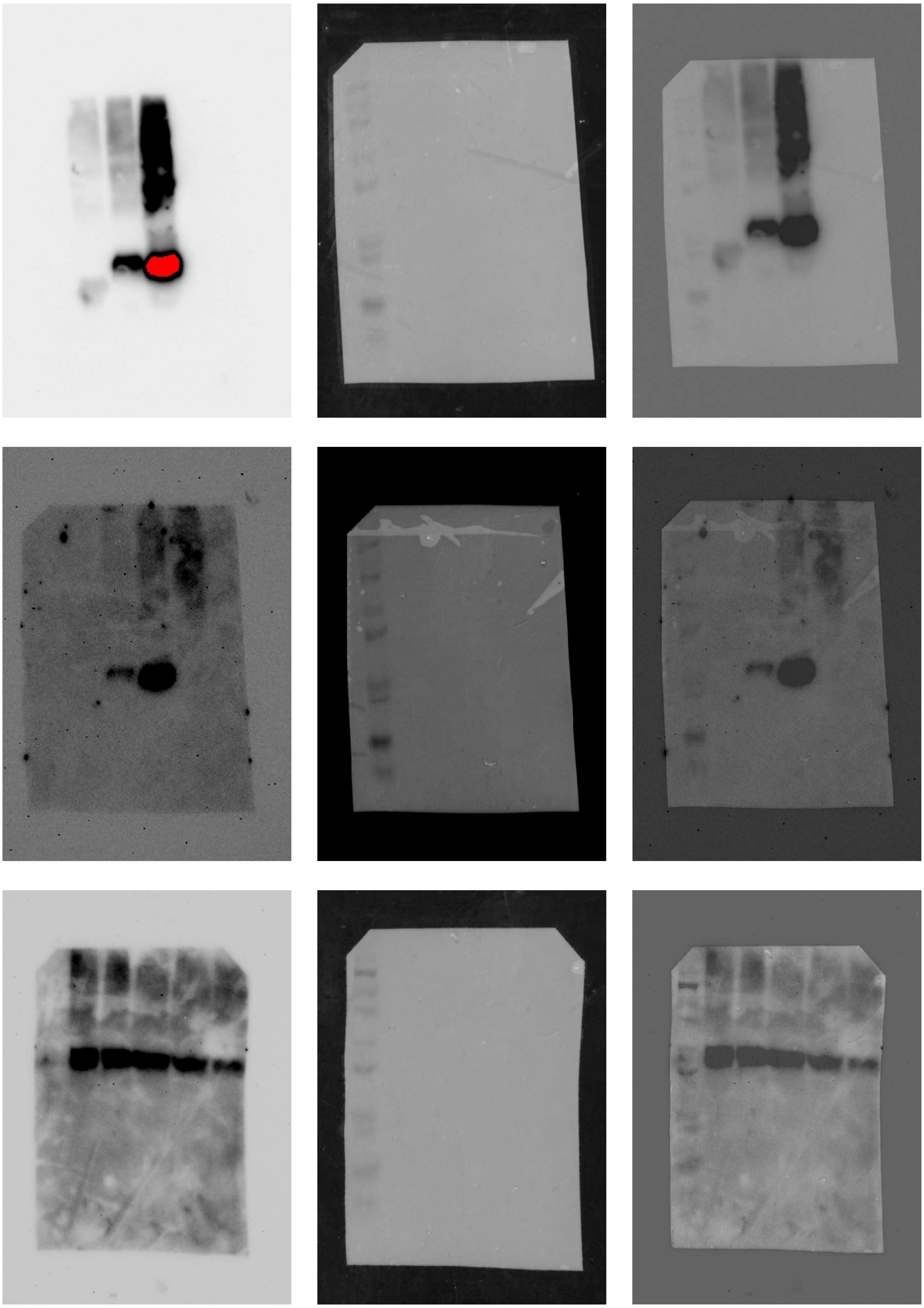
